# Comprehensive evaluation of methods for differential expression analysis of metatranscriptomics data

**DOI:** 10.1101/2021.07.14.452374

**Authors:** Hunyong Cho, Yixiang Qu, Chuwen Liu, Boyang Tang, Ruiqi Lyu, Bridget M. Lin, Jeffrey Roach, M. Andrea Azcarate-Peril, Apoena de Aguiar Ribeiro, Michael I. Love, Kimon Divaris, Di Wu

**Affiliations:** Department of Biostatistics, University of North Carolina, Chapel Hill, NC, United States; Department of Statistics, University of Connecticut, Storrs, CT, United States; School of Computer Science, Carnegie Mellon University, Pittsburgh, Pennsylvania, United States; Research Computing, University of North Carolina, Chapel Hill, NC, United States; Division of Diagnostic Sciences, University of North Carolina, Chapel Hill, NC, United States; Department of Genetics, University of North Carolina, Chapel Hill, NC, United States; Department of Medicine and Nutrition, University of North Carolina, Chapel Hill, NC, United States; Division of Pediatric and Public Health, University of North Carolina, Chapel Hill, NC, United States; Department of Epidemiology, University of North Carolina, Chapel Hill, NC, United States; Division of Oral and Craniofacial Health Sciences, Adam School of Dentistry, University of North Carolina, Chapel Hill, NC, United States; Lineberger Comprehensive Cancer Center, University of North Carolina, Chapel Hill, NC, United States

**Keywords:** metatranscriptomics, metagenomics, differential expression, benchmark, logistic-beta, early childhood caries

## Abstract

Understanding the function of the human microbiome is important; however, the development of statistical methods specifically for the microbial gene expression (i.e., metatranscriptomics) is in its infancy. Many currently employed differential expression analysis methods have been designed for different data types and have not been evaluated in metatranscriptomics settings. To address this gap, we undertook a comprehensive evaluation and benchmarking of ten differential analysis methods for metatranscriptomics data. We used a combination of real and simulated data to evaluate performance (i.e., model fit, type I error, false discovery rate, and sensitivity) of the methods: log-normal (LN), logistic-beta (LB), MAST, DESeq2, metagenomeSeq, ANCOM-BC, LEfSe, ALDEx2, Kruskal-Wallis, and two-part Kruskal-Wallis. The simulation was informed by supragingival biofilm microbiome data from 300 preschool-age children enrolled in a study of early childhood caries (ECC), whereas validations were sought in two additional datasets from an ECC study and an inflammatory bowel disease (IBD) study. The LB test showed the highest sensitivity in both small and large samples and reasonably controlled type I error. Contrarily, MAST was hampered by inflated type I error. Upon application of the LN and LB tests in the ECC study, we found that genes C8PHV7 and C8PEV7, harbored by the lactate-producing Campylobacter gracilis, had the strongest association with childhood dental diseases. This comprehensive model evaluation offer practical guidance for selection of appropriate methods for rigorous analyses of differential expression in metatranscriptomics. Selection of an optimal method increases the possibility of detecting true signals while minimizing the chance of claiming false ones.

## 1 Introduction

### 1.1 Significance

Microbiome has emerged as an undeniable cornerstone for a multitude of health and disease outcomes. Important new insights have been recently gained regarding the pivotal role of microbial dysbiosis in conditions such as obesity, gut disease, cancer, oral and dental diseases (Kaakoush et al., 2012; Tilg et al., 2011; Kilian et al., 2016; Gopalakrishnan et al., 2018). Contemporary investigations now seek to understand not only the composition of microbial communities (i.e., taxonomy) but also their functional activity. Taxonomy is typically ascertained by 16S rRNA or whole genome shotgun (WGS) sequencing with the latter offering several advantages over 16S sequencing, including a better phylogenetic resolution and information on genomic content (i.e., metagenomics). Microbial functional activity can be measured via RNAseq, i.e., metatranscriptomics (Visconti et al., 2019) and metabolomics. Microbial gene expression and metabolism are where the rubber meets the road, as they represent the viable and active members of the microbial community and reflect the biology underlying the microbiome’s interactions with the host and the environment.

Gastrointestinal and oral health, above and beyond their common anatomical, functional, and biological similarities, are now both better understood using models of microbial symbiosis and dysbiosis, while microbiome links between the two have begun to emerge (Olsen and Yamazaki, 2019). In both of these health and research fields, investigations involving metatrascriptomics have provided novel insights. An excellent example includes a multiomics study of inflammatory bowel disease (IBD) highlighting metatrascriptomics associations with the microbial communitys temporal variability, taxonomic, and biochemical shifts (Lloyd-Price et al., 2019). In the oral health domain, Peterson et al. used metatranscriptomics to identify dominant functions associated with dental caries within the dental biofilm of 19 twin pairs. These identified functions supported the microbial community’s biochemical activities related to sugar metabolism and resistance to acid and oxidative stress (Peterson et al., 2014). In another study, metatranscriptomics was used to identify differences in the activity of the subgingival biofilm microbiome between 7 individuals with periodontitis and periodontally-health controls (Duran-Pinedo et al., 2014). A notable recent review of metatranscriptomics analysis of the oral microbiome was recently reported by Duran-Pinedo (Duran-Pinedo, 2021).

Despite the increasing significance and availability of metatranscriptomics data, the development of tailored statistical analysis methods has not kept pace. There are few statistical analysis methods specifically designed to handle microbiome data, and a systematic evaluation of all existing methods that have been borrowed from other areas of high-throughput sequencing analysis (e.g., human studies) has yet to be undertaken.

### 1.2 State of existing differential microbial gene expression analysis methods

Issues and applications requiring special consideration in microbiome data analysis include but are not limited to data normalization (Weiss et al., 2017), clustering of species or genes (Holmes et al., 2012), alpha- and beta-diversity (Willis, 2019), differential abundance/expression (DA/DE) analysis (Chen and Li, 2016; Calgaro et al., 2020; Martin et al., 2020), and metabolome-based pathway analysis (Mallick et al., 2017). Most methods that address these aspects have either been borrowed from analytic pipelines used in other high-throughput sequencing technologies—e.g., bulk RNAseq—or were developed for 16S or WGS data. Most of these currently available methods to analyze metatranscriptome data rely on the joint mapping of microbial DNAseq and RNAseq data (Mallick et al., 2017; Niu et al., 2018; Narayanasamy et al., 2016). However, very few methods have been specifically developed for DE analysis of metatranscriptomics data (Westreich et al., 2018; Hickl et al., 2019). It follows that, while there is ample room for new metatranscriptomics analysis methods development, the existing approaches that have been developed for other data types must be benchmarked and validated prior to being rigorously applied for metatranscriptomics data analyses.

Although the data distributions are regarded as of the same class, either count, normalized count, or proportion with a zero mass, the metatranscriptomics data are more sparse, containing more zeros than metagenomics data. This higher sparsity is a likely result of some taxa, genes, or gene families not being expressed actively, but could also be attributable to some degree to less sensitive measurements of the community’s transcriptome due to technical reasons. It follows that, a systematic evaluation of the performance of existing methods is warranted, so that these methods can be recommended and put to use for the analysis of metatranscriptomics data.

In this article, we focus on DE analysis of metatranscriptomics data in the presence of nuisance information. The DE analysis is one of the fundamental analyses in other transcriptomics data—host bulk RNAseq and single-cell RNAseq, or scRNAseq. It often involves identifying genes whose expression levels are significantly associated with an outcome of interest (e.g., disease vs. health) after controlling for nuisance factors such as batch or block information and the host characteristics (e.g., demographics). While the metatranscriptomics DE analysis is meaningful by itself, it can provide a foundation for joint analysis of the metagenomics and metatranscriptomics data. In this article, the existence of metagenomics data is not required. Instead, the DE analysis methods for RNAseq data will be studied, accounting for the high percentage of zeros, overdispersion, possibly compositional data structures, and the presence of nuisance information.

The evaluation of methods can be done at a microbial species, microbial gene, or microbiome-inferred metabolic pathway level. In this paper, we focus on evaluating methods at the microbial gene level. While genes may have distinct roles within species, it is expected that the same genes across different species have similar functions. Furthermore, many genes are either unique to a species, or are mapped to yet-unclassified species. Metatranscriptomics provides the opportunity to measure the activity of genes, instead of inferring gene expression from microbial genomes. There have been attempts to evaluate differential abundance/composition/expression analysis methods at the species-or the taxon-level that have included many of the commonly used pre-processing and differential abundance analyses (Weiss et al., 2017; Calgaro et al., 2020; Paulson et al., 2013; Zhang et al., 2021). However, those studies did not consider the recently developed statistical models that were specifically designed for zero-inflated over-dispersed counts or compositional data such as MAST (McDavid et al., 2019) and logistic-beta test (Peng et al., 2016). Also, they did not evaluate performances of the methods at the gene level in metatranscriptomics (Weiss et al., 2017; Calgaro et al., 2020; Paulson et al., 2013). The number of species (typically several hundreds) and the number of genes (typically thousands, up to millions) are unique features of metatranscriptomics and metagenomics. The drawbacks become more salient when formally evaluating metatranscriptomics data analysis methods. Our paper aims at overcoming such limitations of previous attempts.

### 1.3 Outline

This paper is organized as follows. In Section 2.1 we list statistical analysis methods that could be used for microbial DE analyses. In the literature of metagenomics or 16S ribosomal RNA (rRNA) data analysis, popular DA/DE analysis approaches include rank-based methods for simple experimental designs, DESeq2 for bulk RNAseq data(Love et al., 2014), multi-dimensional ANCOM (Mandal et al., 2015), and linear regression after log-transformation. We consider the following methods in this simulation study: 1) log-normal (LN) test, 2) logistic-beta (LB) test, 3) Model-based Analysis of Single-cell Transcriptomics (MAST), 4) DESeq2 (Love et al., 2014), 5) metagenomeSeq (MGS) (Paulson et al., 2013), 6) ANCOM-BC (Lin and Peddada, 2020), 7) Linear discriminant analysis Effect Size (LEfSe) (Segata et al., 2011), 8) ANOVA-Like Differential Expression analysis (ALDEx2) (Fernandes et al., 2014), 9) Kruskal-Wallis (KW) test, and 10) two-part Kruskal-Wallis (KW-II) test (Wagner et al., 2011). Some of these methods are based on parametric models (LN, LB, MAST, DESeq2, MGS), some explicitly handle zero-inflation (LB, MAST, MGS, KW2), and one can handle compositional data (LB). Our simulations are more relevant than those of Weiss et al. (Weiss et al., 2017), wherein most of the methods being compared do not reflect the sparse nature of metatranscriptomics data.

To carefully reflect the distributional characteristics of metatranscriptomics data in evaluating those methods in terms of controlling false positive rate and optimizing power, we employed two unique simulation schemes. One is parametric simulations where the parameters were inspired by existing oral and gut microbial data sets. To ensure the similarity between the original and synthesized metatranscriptome data, we assessed the goodness of fit of different data generative models and preprocessing methods. The other scheme is semi-parametric simulations where the existing data sets are randomly sampled for evaluation and were given parametric synthetic disease effects. We have used metatranscriptomics datasets from two body sites for both simulation and application. Two of the datasets are from a genetic-epidemiologic study of early childhood oral health for Zero Out Early Childhood Caries (ZOE)) (Divaris et al., 2019, 2020) wherein oral microbial metatranscriptomics data were generated for over 400 children (297 actual study participants and 116 pilot study participants) aged 3– 5 years for seeking to test the association between microbiome features and early childhood caries (ECC). The third dataset was used to study the association between gut microbiome and inflammatory bowel diseases (“the IBD study”, Lloyd-Price et al. (2019)). We identified the genes of which the expression is significantly associated with ECC (Pitts et al., 2019) or IBD.

Overall, there is a need to comprehensively evaluate the DE methods for metatranscriptome data. To address this knowledge gap, we have designed metatranscriptome-specific simulations to systematically evaluate 10 popular methods. Based on our analysis results, we then offer guidance regarding the selection of appropriate statistical methods. In addition, we make available the code to replicate our work and for use in future studies. Finally, we present applications of the optimized DE methods for the identification of disease-specific microbial genes, potentially as biomarkers of diseases.

## 2 Material and Methods

### 2.1 Differential expression analysis methods

The ten DE analysis methods are evaluated in this simulation study are Log-normal test (LN), Logistic Beta test (LB) (Peng et al., 2016), Model-based Analysis of Single-cell Transcriptomics (MAST) (McDavid et al., 2019), DESeq2 (Love et al., 2014), metagenomeSeq (MGS) (Paulson et al., 2013), ANCOM-BC (Lin and Peddada, 2020), Linear discriminant analysis Effect Size (LEfSe), ANOVA-Like Differential Expression analysis (ALDEx2), Kruskal-Wallis test (KW), and two-part Kruskal-Wallis test (KW-II). Most methods are tailored so that they can control for batch effects while testing associations between gene expression and phenotypes of interest.

The differential ranking (DR) method (Morton et al., 2019) and ANCOM (Mandal et al., 2015) do not provide statistical significance for inference, and ANCOM was optimized to analyze data composed of a relatively small amount of taxonomic units. For these reasons, we do not include these methods. Instead, we consider ANCOM-BC that inherits the philosophy of DR methods and ANCOM but also provides p-values and is computationally efficient. Note that Linear discriminant analysis Effect Size (LEfSe) also does not provide statistical significance. However, it was included in this study due to its high frequent use in differential expression analyses. Since the original version of ANCOM-BC (“ANCOM-BC1”) considers structural zero detection that results in a significant inflation of type I error in our simulations, we consider another version (“ANCOM-BC2”) omitting the structural zero detection procedure and mainly report the results with the results for “ANCOM-BC1” relegated to Supplementary Figures 9–10.

Details such as distributional assumptions, test statistics, and inferential frameworks are reviewed in Supplementary Section 1 of the Supplementary Material.

The scope of our simulations is restricted to tests of differential expression of individual taxa, genes, or gene family levels. On the other hand, there is another group of methods that test the global effect of the whole microbiome, which will not be covered in this simulation study: PERMANOVA (Anderson, 2001), MiRKAT (Zhao et al., 2015), aMiSPU (Wu et al., 2016), and LDM (Hu and Satten, 2020).

Before each test, genes are screened out if they are expressed in only a few participants (the smaller of 2% of the samples or 10 samples). The rationale for such screening is twofold. First, models are not mathematically estimable, when the sample size is less than the number of parameters in the model. Second, even if they are estimable, many large-sample-based inferential procedures have non-negligible finite-sample bias when they are implemented in a small sample or when a gene expression is rare (King and Zeng, 2001). These issues become apparent when 1) the tests that involve logistic regression encounter rare events or 2) the non-zero statistics in two-part models are faced with only a small number of observations with nonzero expression. As evidenced in later sections, all methods except KW fall into either of these two scenarios. To allow for fair comparisons, we apply the same screening rule for each test.

### 2.2 Description of the three metatranscriptomics datasets

To understand and characterize the distributional features of metatranscriptomics data, we leverage three datasets made available by two recent studies involving the human microbiome. The first two datasets (namely, ZOE 2.0 and ZOE-pilot) were generated in a molecular epidemiologic study of early childhood caries (ECC; defined as dental cavities in children under the age of 6) (Pitts et al., 2019) called Zero Out Early Childhood Tooth Decay (ZOE) (Divaris et al., 2020, 2019). In that study, the association between the supragingival oral microbiome and the prevalence of clinically-determined ECC is investigated. The third dataset (namely, the IBD data) was generated in the context of a recent study of the gut microbial ecosystem and its association with Inflammatory Bowel Diseases (IBD) (Lloyd-Price et al., 2019). We base our analyses mainly on the largest (*n* = 300) oral microbiome dataset, or the ZOE 2.0 data, and use the other two datasets for the purposes of validation.

#### 2.2.1 Data One and Data Two: the pediatric dental caries datasets

One of the main aims of the ZOE project is to understand the biological basis of ECC, including the human genome and the oral microbiome. To-date, approximately 5% of the parent cohort (“ZOE 2.0”, 300/6,404) has been carried forward metagenomics, metatranscriptomics, and metabolomics analyses (Divaris et al., 2019). The microbiome samples from these 300 subjects in ZOE2.0 were all sequenced in 2018-19 but in two sequencing batches; the first sequencing batch in May 2018 included 52 samples and the second batch in November 2019 included the remaining 248 samples. In addition, the other 118 participants from the same population were included in a pilot study (“ZOE-pilot”) that included identical phenotyping and biofilm sequencing procedures and their microbiome samples were sequenced at different time points. Similarly, batch effects were evident in the ZOE-pilot data: 60 samples were sequenced in June and an additional 58 samples were sequenced in July of 2017, respectively. Therefore in sum, dental biofilm metatranscriptomics analyses have been done to-date on 418 children ages 3–5 (Divaris et al., 2019). Importantly, the average sequencing depth varied significantly across the sequencing dates, and thus the dates are considered as batches in the parameter selection procedure in Section 2.4.2.

Among the 418 subjects, three and two subjects were excluded for further analyses due to no or very low expression levels in the ZOE 2.0 and the ZOE-pilot data, respectively, resulting in effective sample sizes of 297 and 116. For the purposes of this analysis, ECC was defined as a dichotomous trait, healthy or diseased, based on modified International Caries Detection and Assessment (ICDAS) criteria (Divaris et al., 2020). ECC prevalence was similar in the two ZOE waves, i.e., ZOE 2.0: 49% (147/297) and ZOE-pilot: 50% (58/116). A detailed microbiome analysis protocol for this study has been reported recently (Divaris et al., 2019).

Adapter-trimmed, quality-controlled, demultiplexed Illumina HiSeq sequencing reads were aligned against the human hg19 reference to eliminate host derived reads. Generally speaking, the alignment and data pre-processing followed the procedure described previously (Cho et al., 2022; Divaris et al., 2019). For details, estimates of taxonomic composition, gene family, path abundance, and path coverage were produced from the remaining reads using HUMAnN2 (Abubucker et al., 2012). The resulting reads were scaled into reads-per-kilobase (RPK). We considered additional pre-processing methods including transcript-per-kilobase-million (TPM) and arcsine in Section 3.3.

The total number of gene-species combinations in the ZOE 2.0 metatranscriptome is 535,299; there are 204 distinct species, and 402,937 distinct genes. In the ZOE-pilot sample, there are 439,872 gene-species, 185 distinct species, and 342,004 distinct genes. Total RPKs per sample is on average 13,053,428 in ZOE 2.0 and 2,815,749 in ZOE-pilot. The RPKs are rescaled by dividing by the total RPKs per sample and then multiplying by 4.0 million in ZOE 2.0 and 3.4 million in ZOE-pilot, to make the total expression level for each subject to be 10 times the number of genes. This is a scaled version of TPM-normalized data. In this article, for notational convenience, this scaled version of TPM is referred to as TPM.

There were high proportions of zero gene expressions in both the ZOE 2.0 (80.4%) and the ZOE-pilot (87.9%) metatranscriptomics data. These high zero proportions are comparable and actually higher than what is encountered in the corresponding metagenomics data (75% in ZOE 2.0 and 68% in ZOE-pilot), as illustrated in Figure 1. While a significant number of genes in the metagenomics data are not sparse—one ninth in the ZOE 2.0 (one sixth in ZOE-pilot) of all genes have zero proportion smaller than 20%, virtually all genes, or 94% (97%) are sparse in the ZOE 2.0 (ZOE-pilot) metatranscriptomics data. Specifically, 54% (43%) of genes have ≥ 90% zero proportions in metagenomics compared to 59% (71%) in the metatranscriptomics data of the ZOE 2.0 (ZOE-pilot) data.

**Fig. 1:**
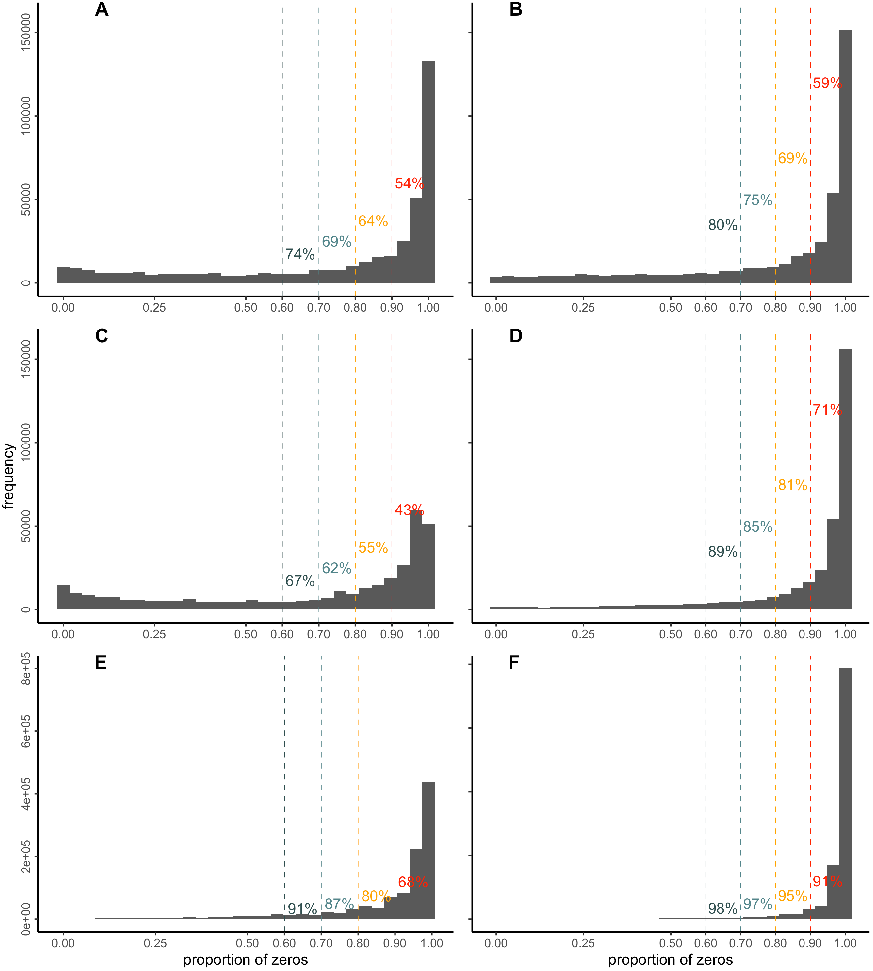
Histogram of zero-proportions at the gene-level in the metagenomics (LEFT) and the metatranscriptomics (RIGHT) data of the ZOE 2.0 (Row 1), the ZOE-pilot (Row 2), and the IBD (Row 3) studies. A. ZOE 2.0 metagenomics; B. ZOE 2.0 metatranscriptomics; C. ZOE-pilot metagenomics; D. ZOE-pilot metatranscriptomics; E. IBD metagenomics; F. IBD metatranscriptomics; Numbers on the histogram represent the proportion of genes of which the zero proportion is greater than or equal to the cutoff values, or the vertical bars left to the numbers.

#### 2.2.2 Data Three: the inflammatory bowel diseases dataset

The multi-omics dataset, including metatranscriptomes via fecal samples, was generated from 132 subjects for a one-year period, with repeated measurement along time per subjects, to study gut microbial ecosystem associated with IBD (Lloyd-Price et al., 2019). For metatranscriptomes, a modified RNAtag-seq protocol was used to create Illumina cDNA libraries which were sequenced on the Illumina HiSeq2500 platform yielding approximately 13 million paired end reads. In this IBD study, the authors have generated the taxonomic and functional profiles (http://huttenhower.sph.harvard.edu/biobakery). Taxonomic profiles of shotgun metagenomes were generated using MetaPhlAn. Functional profiling was performed by HUMAnN2 to quantify gene presence and abundance on a per-species basis (UniRef90s), for both metagenomics and metatranscriptomics. To ensure a reasonable read depth in each sample, the authors only used samples (metagenomes and metatranscriptomes) with at least 1 million reads (after human filtering) and at least one non-zero microbial abundance detected by MetaPhlAn. The dataset includes a total of 1,595 metagenomic and 818 metatranscriptomic samples.

We focus on the cross-sectional features of the metatranscriptomics data distribution, and thus we only examined the baseline information, or the first visit data, including a sample of 104 participants. We further dichotomized participants’ disease status as IBD (i.e., 50 Crohn’s disease and 26 ulcerative colitis cases) versus non-IBD (i.e., 28 ‘control’ participants). Clinic location was considered as a batch effect in that study, and thus was employed in our analyses after dichotomization (the pediatric versus the adult cohorts). The data are publicly available in a compositional format, where the gene expression sums up to one over genes and species for each measurement of a subject. The average proportion of zeros per gene in the IBD metatranscriptomics data is 96.3%, while that in the metagenomics data is 87.8% consistent with the trend of higher zero proportions in metatranscriptomics data over metagenomics data. Figure 1 illustrates the higher zero proportion in the metatranscriptomes over the metagenomes: 91% of genes have zero proportion ≥ 90% in the metatranscriptomics data, compared to 69% in the metagenomics data.

### 2.3 Data scaling and transformation

It is widely acknowledged that the DE analysis in metatranscriptomics depends not only on the DE methods but also on pre-processing of the data. For count data, scaling and transformation are commonly used as part of data normalization to make the data more comparable across samples and/or taxa or to remove distributional irregularities such as skewness. Reads-per-kilobase (RPK), transcripts-per-kilobase-million (TPM), rarefying, and upper-quartile log-fold change normalization (Robinson et al., 2010) are frequently used as the scaling techniques. The examples of transformation include arcsine, logarithm, and variance stabilizing transformation (VST) (Anders and Huber, 2010) among many others.

The strengths and the shortcomings of each scaling and transformation approach have been previously presented and discussed in the literature (McMurdie and Holmes, 2014; Willis, 2019), and also some simulation studies have been done to compare the methods in the differential abundance testing context (Weiss et al., 2017). However, in this paper, we do not pursue a comprehensive comparison of the scaling and transformation methods. Rather, we only consider the three widely used methods, RPK, TPM, and arcsine, because the DE methods evaluation is the main aim of our paper and, also, those scaling and transformation methods provide reasonably good distributional results in the example data.

Among the three pre-processing methods, TPM is defined as RPK divided by the sample sum of RPKs times a constant 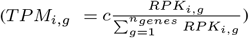 where the constant, *c*, was chosen to reflect the actual scale of the RPKs, or *c* = 5(20) million in the ZOE-pilot (2.0) study. Arcsine-transformation is defined as 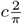 arcsin 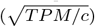. Note that the Beta distribution is for compositional data and thus RPK and TPM data are equivalent for the Beta distribution. The IBD data are available only in a compositional form, and thus, we do not consider the RPK-form of data.

### 2.4 Simulations

#### 2.4.1 Overview

To comprehensively evaluate the performance of the available methods, we considered two approaches: Simulation I. fully parametric simulations and Simulation II. semi-parametric simulations. In Simulation I, three generative models (zero-inflated log normal or ZILN and other two) were used with a comprehensive set of parameters that ranges over the most of the parameter estimates of the example data. The extensive range of parameter sets is to mitigate the potential model-misspecification issue, while we also provide goodness of fit for the generative models. In Simulation II, genes and their expression levels are sampled from the example data followed by artificial insertion of disease effects. The second simulation could provide a validation of the first simulation results, and, at the same time, is a means of appreciating the robustness of the DE methods acroess different datasets. In what follows, we describe how both simulations are done, sequentially.

Simulations are done in R 4.0.3 and the code is available at https://github.com/Hunyong/microbiome2020.

#### 2.4.2 Simulation I: parametric simulations

In Simulation I, multiple scenarios are defined by the following generative models with a nested factorial design. Three data generative model classes were used: zero-inflated log-normal (ZILN), zero-inflated negative binomial (ZINB) and zero-inflated gamma (ZIG) models. On top of each generative model, three factors are further considered: 1) baseline distribution, 2) disease effects, and 3) batch effects (without interaction with disease), each of which is further described in later sections. The parameters representing the three factors are selected after consideration of the estimated parameters of the gene expression in the ZOE 2.0 data as well as the other data sets.

We first obtain the distribution of the baseline parameters by estimating them in each of dental health and disease stratum (H: non-ECC and D: ECC) and sequencing batches. Next, we obtain the distribution of the estimated disease and batch effect parameters. Sets of parameters were chosen for this simulation study so that they suitably represent distributions of parameter estimates in the example data (The details of parameter selection are described in later paragraphs). Once the data distribution is defined, we generate random samples to mimic a small (*n* = 80) study and a large (*n* = 400) study, where all four subgroups of disease-batch combinations are equally sized and there are *n*gene = 10, 000 genes. Then we apply all tests listed in Section 2.1, and derive type I error, false discovery rate (FDR), sensitivity, and accuracy. For most of the methods, type I error (sensitivity) is defined as how often the p-values are less than the 5% significance level, or “the rejection rate at 5%”, in the health (disease) group, and the Benjamini-Hochberg procedure-based adjusted p-values, or “q-values,” are used to derive the FDR instead of type I error. Sensitivity is often called “power” or “true positive rate.” Accuracy is given by the group-size weighted average of 1− type I error and sensitivity. For LEfSe, however, since the p-values are not available, we do not obtain its FDR, although its type I error, sensitivity, and accuracy are derived using the set of the genes declared significant instead of using the 5% cut-off. Since, type I error, sensitivity, and the proportion of the signal genes are sufficient to derive accuracy, and with a large proportion, it is mostly driven by type I error, we do not always present and discuss the results, although all the results are provided in the Supplementary Materials.

##### Generative models

Three generative models are considered: zero-inflated log-normal (ZILN), zero-inflated gamma (ZIG), and zero-inflated negative binomial (ZINB). We do not include the zero-inflated beta (ZIB) distribution, as only a few methods, such as the LB test, model relative gene expressions or abundances rather than their absolute quantities. Furthermore, because ZIB can be considered a compositional transformation of independent ZIG distribution, ZIG-based results should serve as a good proxy for ZIB-based simulations. ZILN is a mixture of log-normal distribution and zeros, where the proportion of zeros is parametrized by *π* and the non-zero values, after the log transformation, follow normal distribution with mean *μ* and variance *σ*^2^. Often in the literature, the model is parametrized using the over-dispersion parameter, *θ*, instead of the variance so that *var*[*Y* |*Y >* 0] = *μ*^2^*θ*. ZILN is frequently used in practice both as a data generating model (Weiss et al., 2017) and as a testing model (e.g., MAST and MGS). A brief description of ZINB and ZIG can be found in Supplementary Section 2.

##### Baseline parameters

A set of baseline parameters uniquely defines a null distribution where there is neither disease nor batch effects. Based on each of these baseline distributions, the disease and/or batch effects are added to form alternative distributions.

The ZILN parameters were estimated for the genes in the ZOE 2.0 dataset for each of the disease and batch subgroups. The method of moments was used for estimation. See Figure 2A for the estimate distribution of 300 randomly selected genes. The first, the second, and the third quartiles of the *μ* estimates are 7.4, 19.7, and 52.6, respectively. Those for the *θ* estimates are 0.7, 1.2, and 1.8. Those for the *π* estimates are 0.34, 0.64, and 0.83. Based on the parameter distribution, we selected sets of baseline parameters for the ZILN model as in Supplementary Table 1 of the Supplementary Material, which is a set of combinations of *μ* ∈ {1, 5, 10*}, θ* ∈ {0.5, 2.0*}, π* ∈ {0.3, 0.6, 0.65, 0.7, 0.75, 0.8, 0.85, 0.9, 0.95}. The parameter estimates for the ZIG model are identical to those of the ZILN model and for this reason are not presented. ZINB model parameter estimates are provided in Supplementary Section 3.3.

**Fig. 2:**
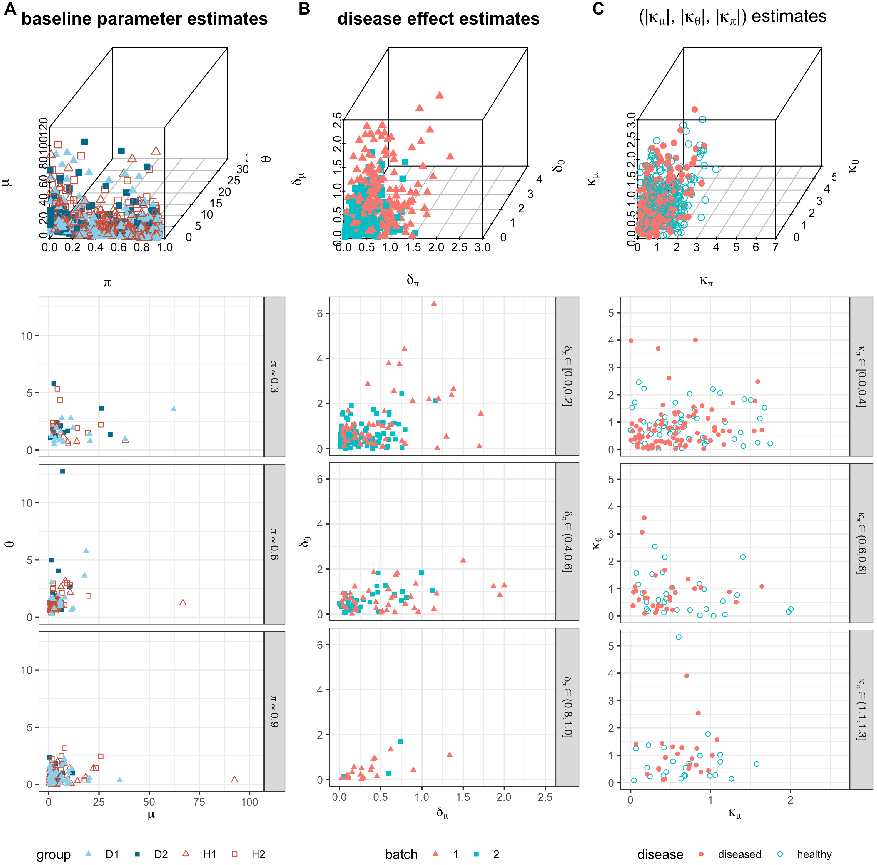
Column A: Parameter estimates of baseline ZILN distributions obtained from the ZOE 2.0 data with the 3-dimensional scatter plot on the top row and each of the subsequent rows representing *π* estimates being within 0.03 from 0.9, 0.6, and 0.3.Column B: Disease effect estimates based on ZILN models obtained from the ZOE 2.0 data in absolute values (|*δ*_*μ*_|, |*δ*_*θ*_|, |*δ*_*π*_|)Column C: Batch effect estimates based on ZILN models obtained from the ZOE 2.0 data in absolute values (|*κ*_*μ*_|, |*κ*_*θ*_|, |*κ*_*π*_|)

To add to our understanding of metatranscriptomics data distributions generated under realistic conditions, we followed the same procedures to estimate the parameters using the ZOE-pilot and the IBD data. Because the ZILN model is the main focus of our simulation study and the expression of the IBD data was only provided in a compositional form, we use these validation data to estimate the ZILN model parameters only. In the ZILN model, *μ* is the only parameter that is affected by scale transformations and most test results are thus invariant to scale transformations except for the NB- and the ZINB-based tests. These estimated parameters from the ZOE-pilot and the IBD data are presented in Supplementary Section 3.4. In Supplementary Figure 2A and 3A, the estimated parameters of the gene expression distribution for genes in the ZOE-pilot and the IBD data have a range that overlaps reasonably with Figure 2A and the sets of parameters in Supplementary Table 1 of the Supplementary Material.

Of note, in this paper we focus on total gene expression for each gene aggregated over all the species in the sample. We further consider parameters for i) expression of each gene-species combination (i.e., the joint data) and ii) species expression marginally across genes (i.e., the species marginal data). The corresponding distributions are provided in Supplementary Section 3.5 of the Supplementary Material. The distributions from the gene-species joint data and the species marginal data are reasonably covered by the parameter sets chosen in this section for all the settings except the ZINB model for the marginal species data, where the parameters are often either not estimable or outlying. In other words, the results are generalizable to the other aspects of data and in most settings, except for the species data with the ZINB model.

##### Disease effects

For each of the baseline distributions, disease effects are further considered to construct the alternative distributions. The disease effects are added in 10% of the genes with randomly perturbation of the direction—i.e., 5% of the genes have higher (lower) expressions are given to the disease group compared to the healthy group, and 90% of the genes do not have any disease effects. Let *δ* ≡ (*δ*_*μ*_, *δ*_*θ*_, *δ*_*π*_) denote the disease effects such that 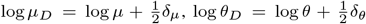, and 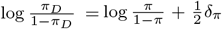, where *ξ*_*D*_ ≡ (*μ*_*D*_, *θ*_*D*_, *π*_*D*_) is the parameter for the diseased group. We simply denote such operation as *ξ*_*D*_ = *g*(*ξ, δ*). The parameter for the healthy group is *ξ*_*H*_ ≡ (*μ*_*H*_, *θ*_*H*_, *π*_*H*_) = *g*(*ξ*, −*δ*).

The disease effect estimates for the ZILN model are estimated from the genes from the ZOE 2.0 dataset assuming that there are no batch effects. These estimates for the 300 randomly selected genes are presented in Figure 2B. The quartiles of the *δ*_*μ*_ estimates are 0.1, 0.2, and 0.5 in the order. Those for the *δ*_*θ*_ estimates are 0.3, 0.5, and 0.9. Those for the *δ*_*π*_ estimates with finite values are 0.2, 0.3, and 0.6. Based on the parameter estimates, we select sets of disease effects for ZILN model as in Table 1 LEFT. Note that the *π* effect of scenario D4 is −1 to maintain consistency of the direction of the effects. The parameter estimates for the ZINB are provided in Supplementary Figure 1B. The corresponding ZILN parameter estimates for the gene-species joint data and the species marginal data are presented in Supplementary Figure 4B and 6B. The estimated ZILN parameters of the gene expression distribution for genes, gene-species, and species have a range that overlaps reasonably with Figure 2B and the sets of parameters in Table 1. The corresponding ZILN parameter estimates for gene expression levels in ZOE-pilot and the IBD data are given in Supplementary Section 3.4, and the selected sets of parameters in Table 1 cover the distributions well.

**Table 1.** Disease effects (LEFT) and batch effects (RIGHT). Additive effect size on log-(for *μ* and *θ*) or logit-(for *π*) transformed scale; e.g., under *μπ* effect I (D6) without batch effects, the disease (healthy) group has one unit higher (lower) nonzero mean on the log scale and one unit lower (higher) zero proportion on the logit scale than the baseline. Under a large effect I (K3), within a disease group, a batch group has 2 *×* 1 higher nonzero mean and dispersion on the log scale and 2 *×* 1 unit lower zero proportion on the logit scale compared to the other batch group.

| No. | $\delta_\mu$ | $\delta_\theta$ | $\delta_\pi$ | name | No. | $\kappa_\mu$ | $\kappa_\theta$ | $\kappa_\pi$ | name |
| --- | --- | --- | --- | --- | --- | --- | --- | --- | --- |
| D1 | 0 | 0 | 0 | null effect | K1 | 0 | 0 | 0 | null effect |
| D2 | 1 | 0 | 0 | $\mu$ effect | K2 | 0.5 | 0.5 | -0.5 | small effect I |
| D3 | 0 | 1 | 0 | $\theta$ effect | K3 | 1 | 1 | -1 | large effect I |
| D4 | 0 | 0 | -1 | $\pi$ effect | K4 | 0.5 | -0.5 | -0.5 | small effect II |
| D5 | 1 | 1 | 0 | $\mu \& \theta$ effect I | K5 | 1 | -1 | -1 | large effect II |
| D6 | 1 | 0 | -1 | $\mu \& \pi$ effect I | | | | | |
| D7 | 0 | 1 | -1 | $\theta \& \pi$ effect I | | | | | |
| D8 | 1 | 0 | 1 | $\mu \& \pi$ effect II | | | | | |
| D9 | 1 | -1 | 0 | $\mu \& \theta$ effect II | | | | | |
| D10 | 0 | -1 | -1 | $\theta \& \pi$ effect II | | | | | |

##### Batch effects

For each of the alternative distributions, we further considered batch differences. As batch effects are in most cases nuisance parameters, we considered limited settings and only binary effects were modeled. Batch effects are reflected on parameters for each health and disease group in a similar way as that of disease effects.

Let *κ* ≡ (*κ*_*μ*_, *κ*_*θ*_, *κ*_*π*_) denote the batch effects such that *ξ*_*d*,1_ = *g*(*ξ*_*d*_, *κ*) and *ξ*_*d*,2_ = *g*(*ξ*_*d*_, −*κ*) are the distribution parameters for disease group *d*(*d* = *D*or*H*) in batches 1 and 2, respectively. Alternatively, we denote *ξ*_*D*,1_ = *g*(*ξ, δ, κ*).

The batch effects for the ZILN model are estimated for the genes in the ZOE 2.0 dataset assuming that there are no batch effects. A random sample of these are presented in Figure 2 C. The quartiles of the *κ*_*μ*_ estimates are 0.3, 0.6, and 1.0 in the order. Those for the *κ*_*θ*_ estimates are 0.3, 0.7, and 1.2. Those for the *κ*_*π*_ estimates with finite values are 0.4, 0.8, and 1.4.

Based on the ZILN parameter estimates, we select sets of batch effect parameters for the ZILN model as in Table 1 RIGHT. The parameter estimates for the ZINB are provided in Supplementary Figure 1C. The corresponding ZILN parameter estimates for the gene-species joint data and the species marginal data are given in Supplementary Figures 4C and 6C. The estimated ZILN parameters of the gene expression distribution for genes have a range that overlaps reasonably with Figure 2C and the sets of parameters in Table 1. The corresponding ZILN parameter estimates for gene expression levels in ZOE-pilot and the IBD data are given in Supplementary Section 3.4, and the selected sets of parameters in Table 1 provide good coverage of the distributions.

#### 2.4.3 Simulation II: semi-parametric simulations

For each of the three datasets—ZOE 2.0, ZOE 2.0 pilot, and IBD— we randomly sample 10,000 genes among the prevalence-filtered (≥ 10%) genes. For a prespecified number (100 and 1000) of genes, synthetic disease effects are added, and we call them “signal genes.” The manipulated data are analyzed with the ten methods, and their performances are evaluated the same way as in Simulation I over 10 replicates.

The semi-parametric simulations were done for each of the ZOE 2.0, the ZOE 2.0 pilot, and the IBD data. For each data, *n*_genes_ = 1,000, 10,000 genes were randomly selected among which *n*_signal_ = 100, 1,000 signal genes were further selected for differential (*μ, π*) disease effects. Note that we did not consider the *θ* effects, as they are not of interest in general, and *μ* and *π* are the parameters that quantify the overall mean expression levels. For this task, we utilize the set of disease effects estimates obtained in Section 2.4.2 and choose only the subset tuples (*δ*_*μ*_, *δ*_*π*_) such that |*δ*_*μ*_| ≥ 2. From this large effect size tuples, *n*_signal_ tuples were randomly selected and were assigned to each of the signal genes. This effects size tuples were then applied to the disease groups only with the following rule.

Let (*δ*_*μ,g*_, *δ*_*π,g*_) be the effect size assigned to a signal gene *g*. Then *δ*_*π,g*_ is first applied to flip a certain number of zeros to non-zeros or non-zeros to zeros, and then *δ*_*μ,g*_ is applied to rescale the non-zero values. Let 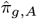 denote the sample zero proportion of the disease group *A* =D, H. Also, let 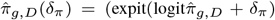 denote the differentiated zero proportion of gene *g* in the disease group *D*, where 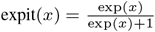 and 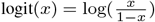. Let *Z*_*g,D*_ be a random realization from the binomial distribution with *n*_*D*_ trials and the success probabilities 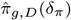, where *n*_*D*_ is the sample size of the disease group *D*. The health group counterparts are defined similarly: 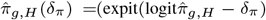 is the differentiated zero proportion, and *Z*_*g,H*_ is a binomial random draw with *n*_*H*_ trials. If 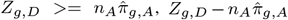 many non-zero-expressed subjects are randomly chosen and are replaced with zeros. On the contrary, if 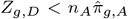, each of the randomly chosen 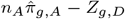 many zero-expressed subjects are given the expression values of one of the randomly chosen non-zero subjects. Once the *π*-disease effects are manipulated, the non-zero expression values, *Y*_*g,i*_ > 0, are transformed into exp(log(*Y*_*g,i*_) *± δ*_*μ*_) with + (−) for *i* in group *D* (*H*). Note that the signs of (*δ*_*μ*_, *δ*_*π*_) are preserved, so that for some genes *δ*_*μ*_ (*δ*_*π*_) is positive while for other genes it can be negative.

Then each of the ten analysis methods was applied to each of the semi-parametric simulated datasets and its performances were evaluated under the same metrics as being used in Simulation I (parametric simulations).

## 3 Results

### 3.1 Overall results

We present the results with respect to two main aims: 1) Goodness of fit of the different data generative models and preprocessing methods at the microbial gene level, and 2) the performances of the 10 DE methods in the real data-based simulations in both the parametric setup and the semi-parametric setup. The results suggest that the log-normal distribution fits reasonably well to the ZOE 2.0 data after the TPM transformation. We find, in both the parametric simulations and the semi-parametric simulations, the LB tests showed the highest sensitivity while controlling for the type I error in a large sample. In addition, MGS has comparably high sensitivity with a well-controlled type I error. MAST and ANCOM-BC often suffers from inflated type I error even with a large sample size. We apply the statistical analysis methods that show the highest performance to the sizeable metatrascriptomics dataset generated in two aforementioned studies: ZOE 2.0 of oral microbiome and the IBD study of gut microbiome, to identify significant genes that are associated with the disease phenotypes. It’s worthy mentioning that, to coordinate the data format output from mapping software, we follow a reasonable approach to obtain the expression data at the gene level in RPKs as the summed (or marginalized) RPK over all species per gene before DE analysis. Without losing the generality of our study, other aspects of microbial activities—the gene expression per gene per species (“gene-species joint data”) and the gene expression per species as aggregation over all genes in one species (“species marginal data”)—are also considered.

### 3.2 Goodness-of-fit of the generative models

#### 3.2.1 Goodness-of-fit in the ZOE data

We use the Lilliefors procedure (Lilliefors, 1967) to evaluate the goodness-of-fit of the generative models (ZILN, ZIG and ZINB) in the ZOE data. The Lilliefors procedure is a data-adaptive version of the Kolmogorov-Smirnov (KS) test (Smirnov, 1948), where the empirical distribution function (EDF) of a specific gene is compared to the cumulative distribution function (CDF) of the estimated model rather than to a fixed CDF. We randomly select 300 genes and the maximal difference of the EDF and the CDF for each gene is calculated. A p-value is calculated for each gene based on a null distribution generated by Monte Carlo simulations, and the histogram of the p-values and the proportion of the p-values less than the 0.05 threshold are reported.

The Lilliefors procedure is only applied to the non-zero values and ZINB is not evaluated with this procedure. This is because the KS test and the Lilliefors procedure are designed for continuous distributions, while the zero-inflation components in ZILN and ZIG have virtually perfect goodness-of-fit. The Beta distribution, the continuous part of the ZIB model, is also considered in this evaluation. For ZINB distributions, a graphical comparison is provided as an alternative to the quantitative procedure. Estimation of distribution parameters is based on three scaling/transformation methods: RPK, TPM, and arcsine.

Figure 3B suggests that for a small sample, all three generative models appear to have a reasonable fit to the data—the rejection rates are at most 10%. However, the overall high rejection rates in Figure 3A (ZOE 2.0) suggest that the reasonable high rejection rates in Figure 3B (ZOE-pilot) are probably due to typical lower testing power when sample size is small. Despite the high rejection rates in many settings in Figure 3A, the log-normal model shows a consistently good fit. In both Figures 3A and 3B, the TPM normalization is shown to provide a better fit compared to the RPKs, and the arcsine transformation further enhances the rejection rate. It is noteworthy that the results are almost identical between the TPM and the arcsine transformations for the log-normal distribution. This is because, with a large number of genes in the data (*n*_*genes*_ > 300, 000 for ZOE 2.0), most compositional (*T PM/c*) values are close to zero and, consequently, the arcsine transformation is essentially equivalent the square root transformation with scaling:

**Fig. 3:**
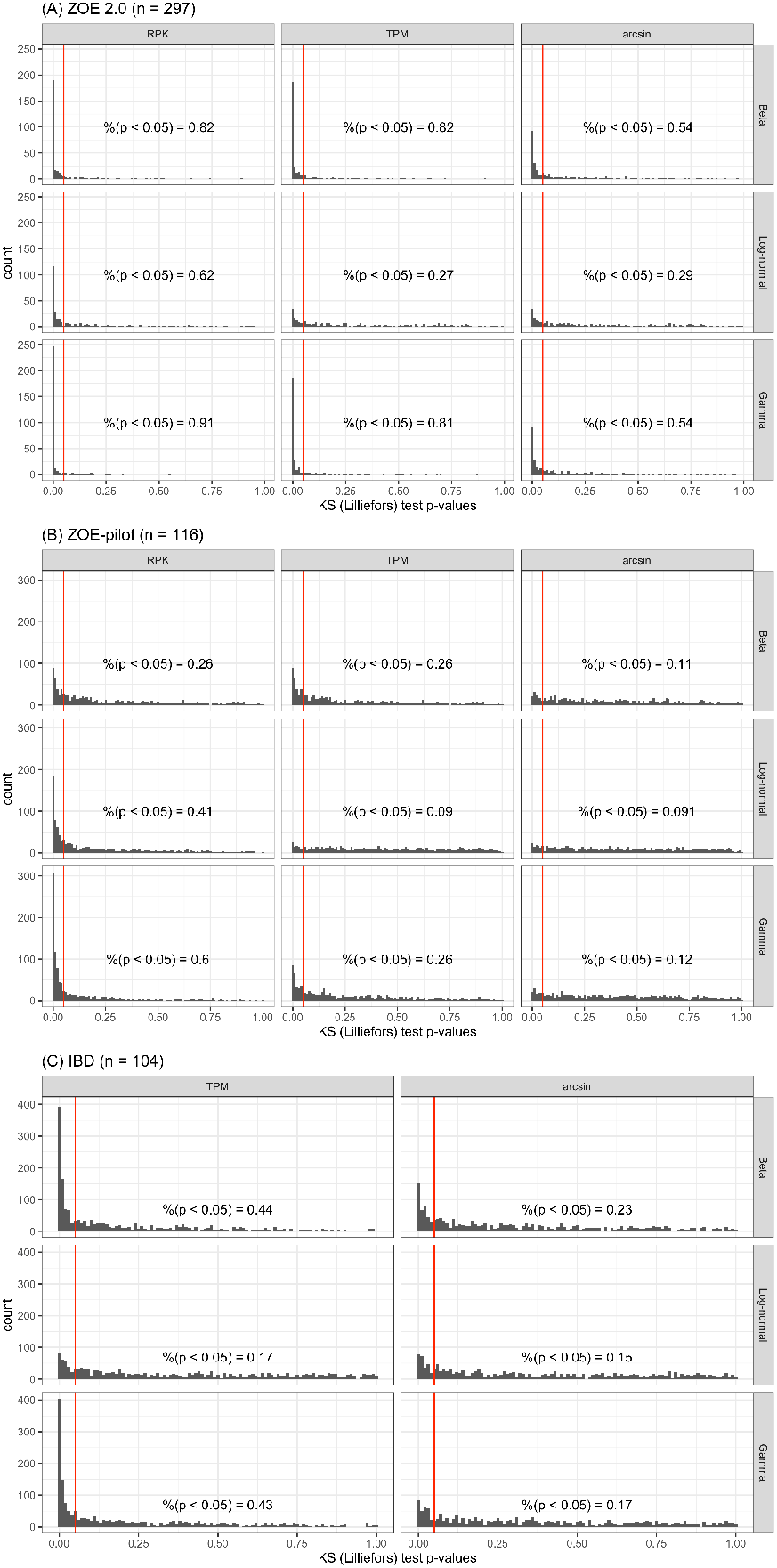
Goodness of fit (Kolmogorov-Smirnov) test results for Beta, Log-normal, and Gamma distributions (rows) with different scaling/transformation methods (columns). The top nine histograms (A) are based on the ZOE 2.0 data (*n* = 297), the middle nine graphs (B) are based on the ZOE-pilot data (*n* = 116), and the bottom six graphs (C) are based on the IBD data (*n* = 104).

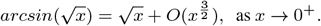

This implies that the arcsine transformation is merely a location-shift transformation in the log-normal model for most of compositional values.

The model fit of the ZINB distribution is illustrated in Supplementary Section 4.1 for a couple of randomly chosen genes after rounding values to nearest integers. The results suggest that the ZINB distribution has overall a reasonable fit to the RPK or the TPM transformed data, but has a poor fit to the arcsine transformed data.

#### 3.2.2 Goodness-of-fit of the generative models – the IBD data

Figure 3C shows that the goodness of fit of the IBD data is overall worse than that of the ZOE-pilot data that have a similar sample size. However, for both data, the log-normal distribution with either the TPM or the arcsin transformation yields the best fit.

### 3.3 Simulation I (parametric simulations) results

#### 3.3.1 Type I error and false discovery rate (FDR)

For the ZILN model, we analyze type I error and FDR results under D2 (*μ*_*D*_ *> μ*_*H*_) scenario as presented in Figure 4. Overall, type I error and FDR are well or reasonably controlled for several methods such as MGS, KW, KW-II, LN, ALDEx2. On the contrary, LEfSe with its default test thresholds—0.05 for Kruskal-Wallis and 2 for LDA—has higher than 5% type I error frequently, implying that LEfSe requires tuning of the thresholds to avoid high type I error, potentially with lower thresholds.

**Fig. 4:**
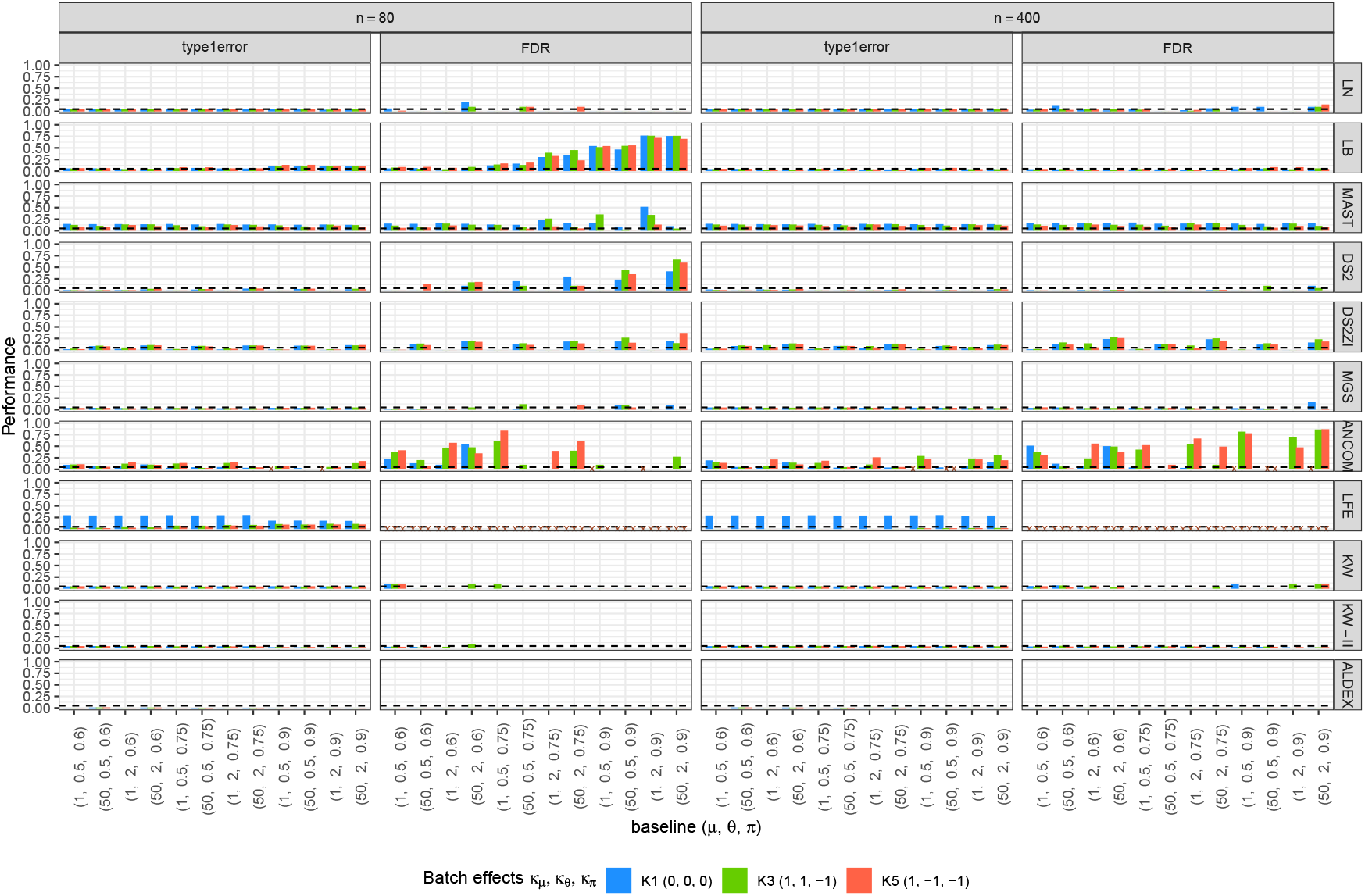
Type-I error rates and FDR for ZILN models. Columns correspond to sample sizes and evaluation criteria, rows are different tests, the *X*−axis represents baseline distributions, and colors indicate batch effects. The dotted horizontal lines show the significance level (5%). A failure in evaluation is marked as *×* to be discerned from zero. DS2 = DESeq2, DS2ZI = DESeq2-ZINBWaVE, ANCOM = ANCOM-BC2, LFE = LEfSe, ALDEX = ALDEx2.

MAST has type I error and FDR that are often higher than the nominal significance level even for a larger sample size. ANCOM-BC2 also frequently has inflated type I error and FDR, which is more evident under the batch-effects scenarios possibly due to model mis-specification. Its type I error and FDR are often amplified with a larger sample size, indicating that the error is not due to a finite sample size but could be a systematic bias. ANCOM-BC1, the original version of ANCOM-BC with structural zeros, has a considerable inflation of type I error for most of the scenarios, especially under the high zero-proportion scenarios. See Supplementary Figures 9–10. Identification of the structural zeros in ANCOM-BC1 could be unreliable under the high zero-proportion settings.

Otherwise, type I error is less stably controlled especially in the two-part models, such as LB and DESeq2-ZINBWaVE. This is true for LB especially when the zero-inflation (or zero proportion) parameter is high. This is likely a consequence of the nonzero part of those models relying on a small number of nonzero values, that causes high finite sample bias. For example, for a ZILN sample of size *n* = 80, *π* = 0.9 means that there are only 8 nonzero values on average and that large sample theory may not be applicable. The inflated or deflated type I error of those methods dissolves or attenuates when the sample size is large, further implying that finite sample bias is the culprit. Thus, we suggest that, when the sample size is not large, and proportion of zeros is high, two-part models are not recommended without knowledge that the posited distribution of the test agrees with the true underlying distribution.

DESeq2 has a very low type I error when used to model these zero-inflated data. Because it was designed for negative binomial distributions without zero-inflation, this may not be a surprising result. On the other hand, DESeq2-ZINBWaVE has on average higher type I error than DESeq2. However, it has higher-than-nominal type I errors for larger baseline nonzero mean values, and the inflation becomes even larger for a large sample, implying that the aberration may not be attributable to the finite sample bias. Designed for the scRNAseq and with its unstable control of type I error, the ZINB-WAVE extension of DESeq2 should be used with caution.

Furthermore, the type I error and FDR results under ZINB and ZIG models are presented in Supplementary Figures 14 and 17.

#### 3.3.2 Sensitivity in a small sample (*n* = 80)

The rejection rates at 5% cutoff for the signal genes under alternative distributions, or the sensitivity of the tests, are illustrated in Figure 5 for selected scenarios of the ZILN model and sample size of *n* = 80. The corresponding results for sample size of *n* = 400 are presented in Figure 6 of Section 3.3.3. The simulations based on the generative models other than ZILN are discussed in Section 3.3.4. The full results under all scenarios, i.e., all combinations of generative models, baseline distribution, batch effects, and disease effects, are provided in Supplementary Figure 9 to 16 in Supplementary Sections 5.1 to 5.4. We further vary either the significance level or the effects sizes to provide a more comprehensive landscape of the test performances in Sections 3.3.5. In what follows, sensitivity, illustrated in Figure 5, is discussed according to different disease effect scenarios (D2–D8).

**Fig. 5:**
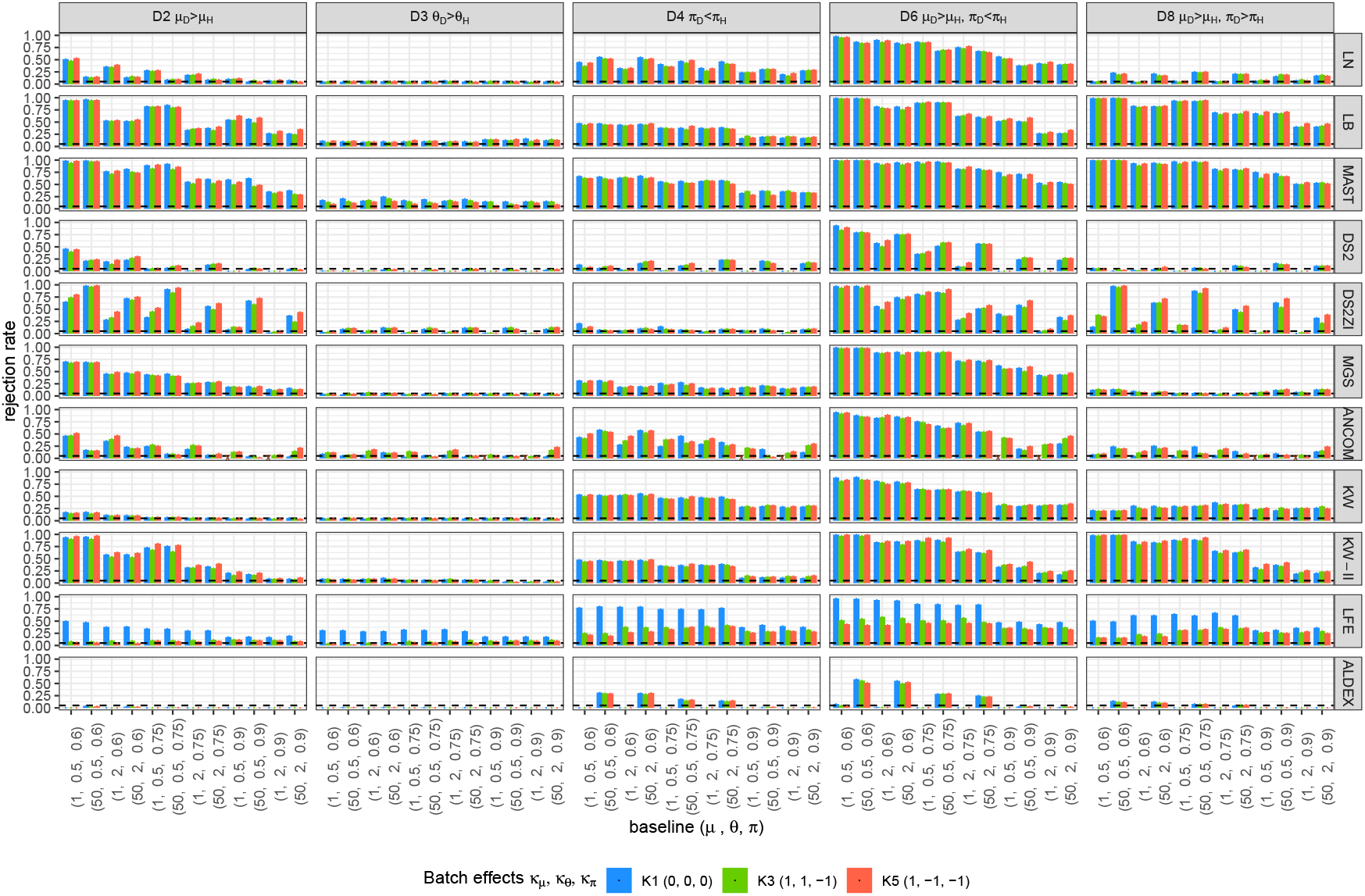
Sensitivity under alternative ZILN distributions for a small sample (*n* = 80). Columns and rows correspond to tests and alternative distributions, respectively, the *X*−axis represents baseline distributions, and colors represent batch effects. The dotted horizontal lines denote the significance level (5%). A failure in evaluation is marked as *×* to be discerned from zero. DS2 = DESeq2, DS2ZI = DESeq2-ZINBWaVE, ANCOM = ANCOM-BC2, LFE = LEfSe, ALDEX = ALDEx2.

**Fig. 6:**
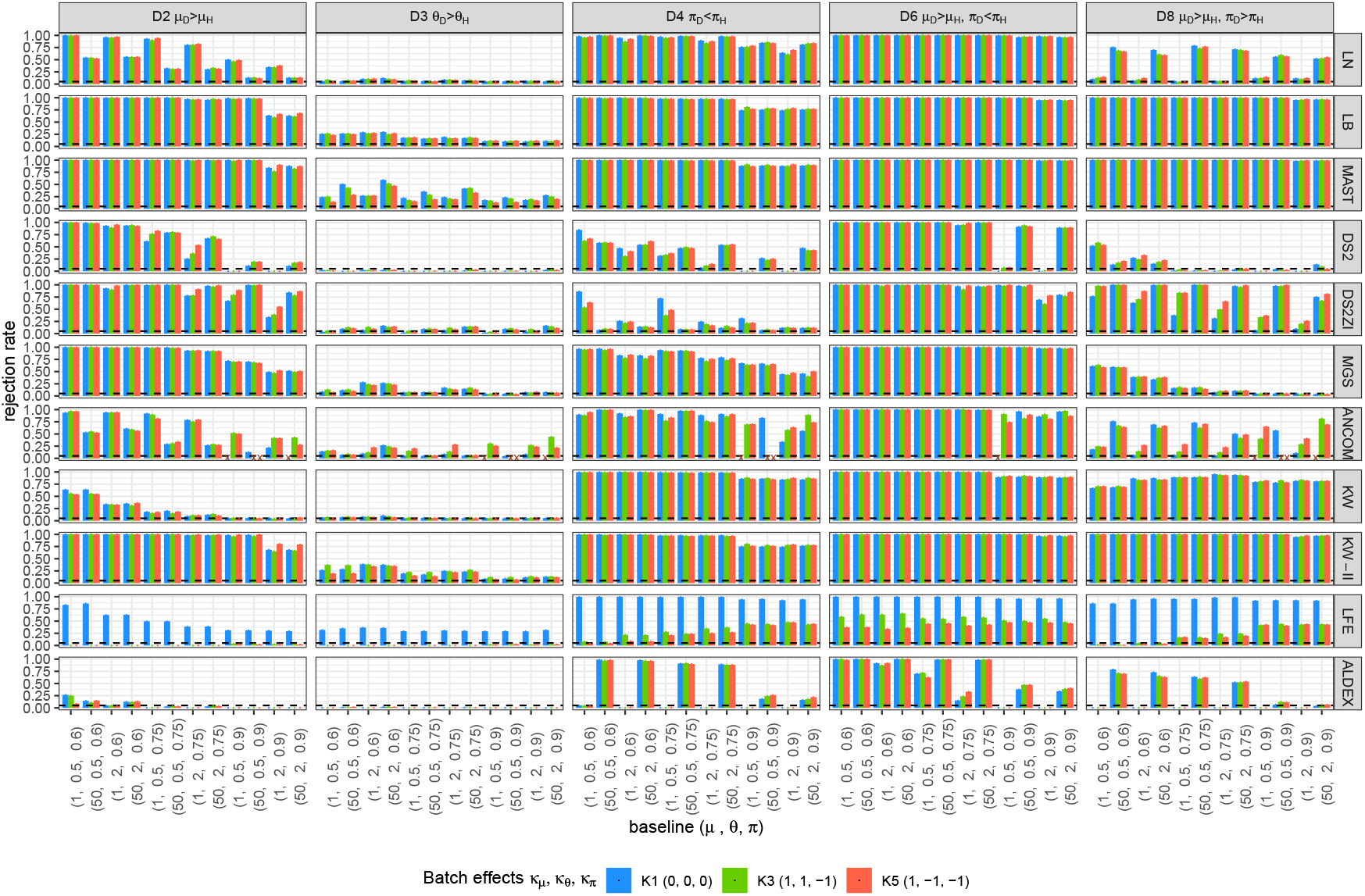
Sensitivity under alternative ZILN distributions for a large sample (*n* = 400). Columns and rows correspond to tests and alternative distributions, respectively, the *X*−axis represents baseline distributions, and colors represent batch effects. The dotted horizontal lines denote the nominal significance level (5%). A failure in evaluation is marked as *×* to be discerned from zero. DS2 = DESeq2, DS2ZI = DESeq2-ZINBWaVE, ANCOM = ANCOM-BC2, LFE = LEfSe, ALDEX = ALDEx2.

**D2 (***μ*_*D*_ > *μ*_*H*_ ). When the disease status is only associated with the difference in nonzero means (*μ*), many methods have high sensitivity for most of the baseline scenarios—LB, MAST, DESeq2-ZINBWaVE, MGS, LEfSe, KW-II—and methods such as LN and ANCOM-BC2 have moderate level of sensitivity over various settings. However, while the good amount of sensitivity of MAST, DESeq2-ZINBWaVE, LEfSe, and ANCOM-BC2 comes at a cost of inflated type I error (and FDR), the type I error of LB is relatively reasonably controlled for large sample size and that of MGS is well controlled. KW-II, that often has one of the highest sensitivities, has relatively weak sensitivity for the high zero-proportion scenarios. LN has good sensitivity for many baseline scenarios but lacks sensitivity when the zero-proportion is high (*π* = 0.9). This is due to the bias from model misspecification of the LN model. KW suffers from low sensitivity with even smaller *π*. DESeq2 has a reasonably good sensitivity when the zero inflation is not high (*π* ≤ 0.6). However, it often has lower than 5% sensitivity when the data are sparse. This again can be explained by DESeq2’s inability to model zero-inflation. Finally, ALDEx2 suffers from lack of power, sensitivity lower than 5%, which can be explained in part by the Dirichlet prior Dir 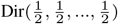 unduly dominating the expression levels of the relatively sparse and low-expressed compositional data.

**D3 (***θ*_*D*_ *> θ*_*H*_ ). Most tests lack power in detecting difference in *θ* (D3), which is expected as all the tests considered in this paper detect the marginal or conditional mean differences and *θ* difference alone does not affect the mean. However, there are quite a few methods that have sensitivity greater than 5%. Methods with inflated type I error, such as MAST, are expected to have rejection rates higher than 5%. The equal variance assumption that is implied in methods could also be a source of the inflation.

**D4 (***π*_*D*_ *> π*_*H*_ ). Differential disease effects only in *π* are captured by methods such as LB, LN, KW, LEfSe, and MAST. MAST and LEfSe’s higher sensitivity than the other methods is counterbalanced by inflated type I error rates. Under the setting D4, relatively low sensitivity for baseline *π* = 0.9 and high sensitivity for baseline *π* = 0.6 can be explained by the design of the experiments. When the baseline *π* is close to 1 or 0, the absolute difference (*π*_*D*_ −*π*_*H*_) between two groups is relatively smaller than that when the baseline *π* is close to 0.5.

**D6 (***μ*_*D*_ *> μ*_*H*_, *π*_*D*_ *< π*_*H*_ ). Sensitivities are higher for D6 (*μ*_*D*_ *> μ*_*H*_, *π*_*D*_ *< π*_*H*_) than for both D2 (*μ*_*D*_ *> μ*_*H*_) and D4 (*π*_*D*_ *< π*_*H*_), as D6 is expected to have larger marginal mean differences than D2 and D4. As a result, most tests have sensitivities ≥ 0.50 for *π* ≤ 0.9 including LN and KW.

**D8 (***μ*_*D*_ *> μ*_*H*_, *π*_*D*_ *> π*_*H*_ ). The disease effect scenario D8 is complicated, as the signal from *μ* difference and that from *π* difference offsets the effect on the marginal mean. Thus, tests based on marginal models, i.e. single-part models such as LN, KW, MGS, DESeq2, and ANCOM-BC, inherently cannot avoid low sensitivity under this scenario, because they do not separate two opposite signals from two parts. Consequently, they have lower rejection rates under D8 than under either D2 or D4. In contrast, two-part models (LB, MAST, KW-II) entertain the two distinct signals resulting in almost the same sensitivity as in D6.

##### Other scenarios involving *θ* (D5, D7, D9, D10)

Other scenarios involving *θ* such as D5 (*μ*_*D*_ *> μ*_*H*_, *θ*_*D*_ *> θ*_*H*_), D7 (*θ*_*D*_ *> θ*_*H*_, *π*_*D*_ *< π*_*H*_), D9(*μ*_*D*_ *> μ*_*H*_, *θ*_*D*_ *< θ*_*H*_), and D10(*θ*_*D*_ *< θ*_*H*_, *π*_*D*_ *< π*_*H*_) do not have remarkable differences in results than the corresponding scenarios without *θ* effects, i.e., D2, D4, D2, and D4, respectively (Supplementary Section 5.1). This is expected, because *θ* differences do not affect the marginal mean difference and the methods considered in this paper treat *θ* as a nuisance parameter.

##### Sensitivity in the presence of batch effects

The presence of batch effects affects sensitivity even when the batch information is incorporated in the tests. This could be due to the fact that batch effects are made multiplicatively in the generative models, while the models in tests only consider main effects of diseases and batches without their interaction. However, the unevenness of sensitivity across different batch-effect scenarios is neither dramatic nor systematic. The patterns, e.g. higher sensitivity for MAST and LB, lower sensitivity for large *π* values and so forth, discussed in earlier sections, still hold across different batch-effect scenarios.

##### Summary

In reality, differential expression only in nonzero mean (D2), only in zero proportion (D4), or in both nonzero mean and zero proportion with the opposite direction (D6), is of most interest and is more feasibly observed than the others. Thus, the LB and MGS tests that have high sensitivity under D2 and D6 and the LN test that has high sensitivity under D4 and D6 are noteworthy. KW and KW-II tests have high sensitivity under D6. However, KW has very low sensitivity for most of the settings of D2 and KW-II suffers from low sensitivity when *π* = 0.9 under all of D2, D4, and D6.

#### 3.3.3 Sensitivity in a large sample (*n* = 400)

As expected, rejection rates are higher in a larger sample (*n* = 400) and many tests under most scenarios have sensitivity close to 1. See Figure 6 The patterns for the larger sample size are mostly the same as those under the smaller sample size; MAST, LB, and MGS have the highest sensitivity under most scenarios, two-part models have higher sensitivity than single-part models when signals are in the opposite directions as in D8, and *θ* difference (D3) is not properly detected for most of the tests.

#### 3.3.4 Sensitivity under ZINB and ZIG

The patterns of rejection rates under ZINB and ZIG models are not very different from those under ZILN models. The full results are presented in Supplementary Figures 12 –16.

#### 3.3.5 Sensitivity in varying thresholds and effect sizes

To provide a broader view of the sensitivity of the tests we vary the significance level and the disease effect size, respectively. Figure 7 presents the sensitivity according to the cut-off values ranging from 0 to 0.2 under a few baseline distributions and disease effects scenarios of the ZILN model without batch effects. Each curve either dominates or is dominated by the others for most of the settings uniformly over the cut-off values in [0.0, 0.2], which suggests that the pattern of the previous results is mostly preserved with a different choice of cut-off values.

**Fig. 7:**
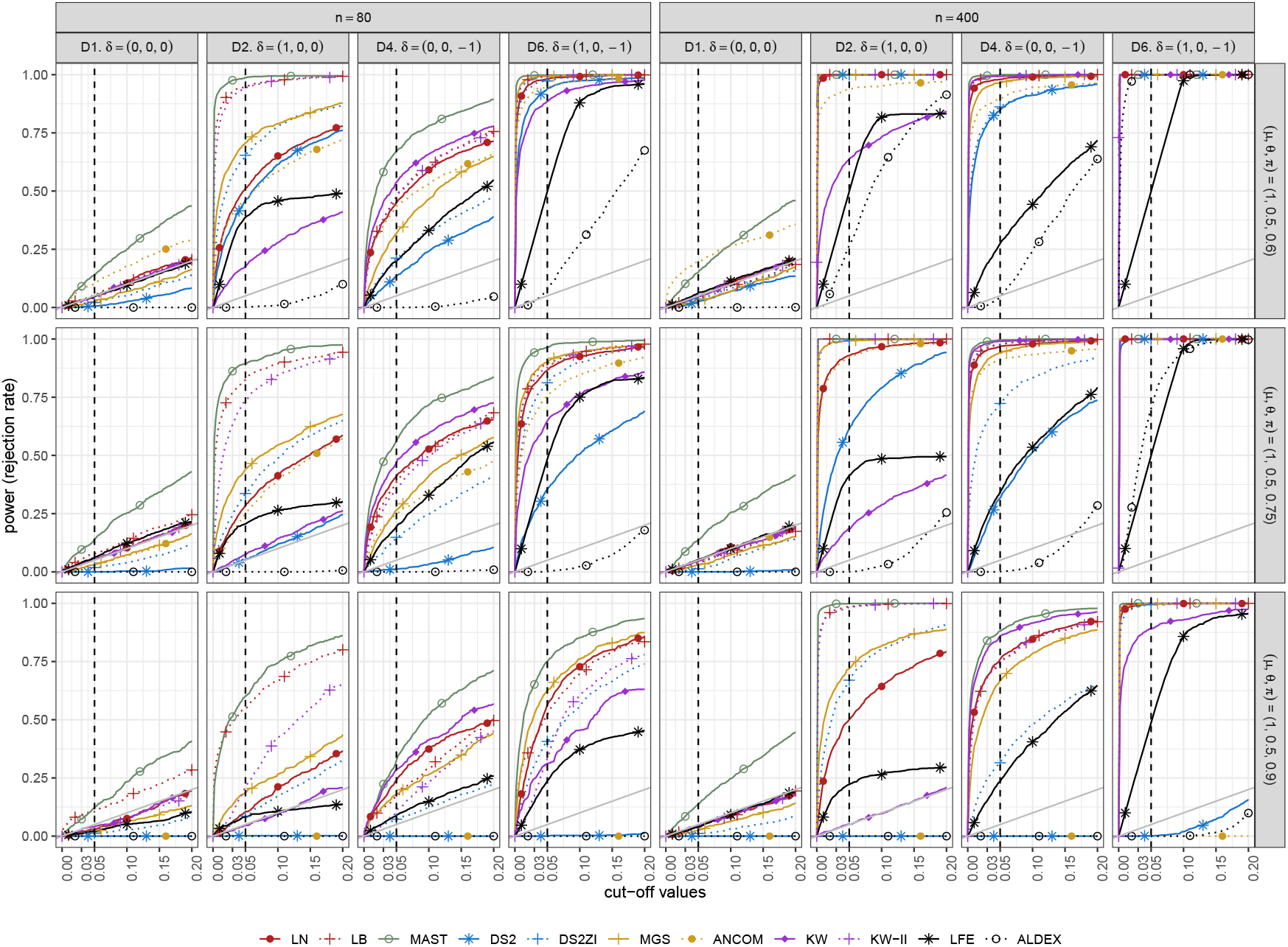
Sensitivity curves of differential expression tests according to different cut-off values. No batch effects are simulated in these scenarios. The gray solid diagonal lines denote the nominal significance level.

Figure 8 illustrates the sensitivity of the tests under different sizes of disease effects for a subset of the baseline scenarios and *n* = 80 without batch effects. The pattern of lower (higher) sensitivity for smaller (larger) disease effect sizes was expected. However, it is noteworthy that when there are only small disease effects on the non-zero mean (i.e., Scenario D11), some methods have virtually zero sensitivity (DESeq2 and KW) or very low sensitivity (LN), suggesting that to compensate for the lack of fit, the effect size must be sufficiently large.

**Fig. 8:**
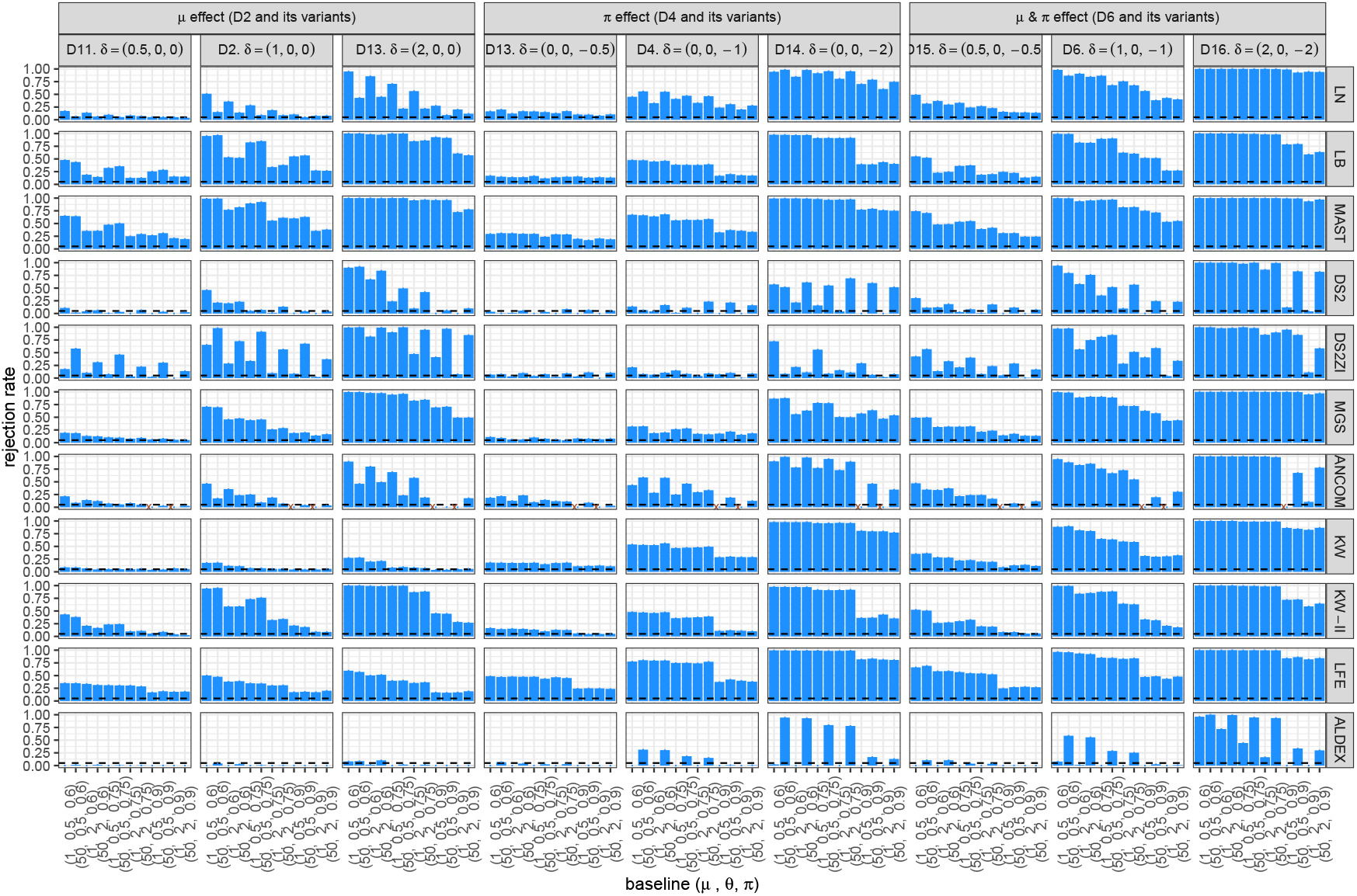
Sensitivity of the differential expression tests according to different effect sizes. No batch effects are simulated in these scenarios. A failure in evaluation is marked as *×* to be discerned from zero. DS2 = DESeq2, DS2ZI = DESeq2-ZINBWaVE, ANCOM = ANCOM-BC2, LFE = LEfSe, ALDEX = ALDEx2.

### 3.4 Semi-parametric simulation results

The semi-parametric simulation results (Figure 9) mostly confirm the parametric data simulation results. MAST, ANCOM-BC, and LEfSe have high type I error and FDR, and those for LB are high for small sized data (ZOE 2.0 pilot and IBD) but are well controlled for the larger data (ZOE 2.0). Type I error and FDR for the other methods are reasonably controlled, and especially those for ALDEx2 are lower than the threshold.

**Fig. 9:**
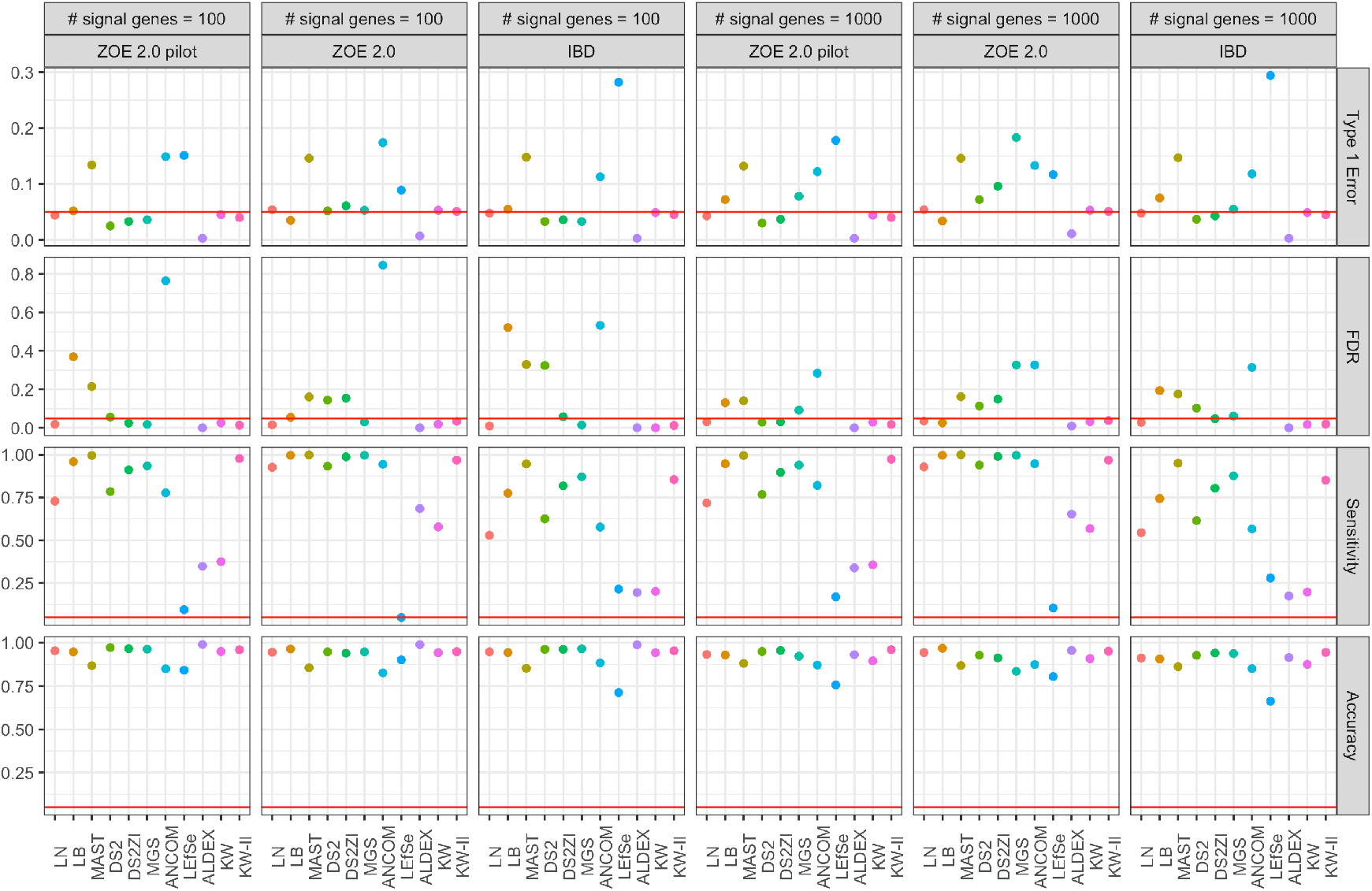
Performances of the analysis methods under semi-parametric simulations. 10,000 genes were selected, some of the genes, the portions of which are specified in the panel heads, were given artificial disease effects. DS2 = DESeq2, DS2ZI = DESeq2-ZINBWaVE, ANCOM = ANCOM-BC2.

Sensitivity is often high for most of the methods except KW, LEfSe, and ALDEx2. When the sample size is relatively large (e.g., ZOE 2.0) and the proportion of the signal genes is large, higher sensitivity is observed for most of the methods—LB, MAST, DESeq2, DESeq2-ZINBWaVE, MGS, ANCOM-BC2, LN. Among those, LB and LN have type I error and FDR less than or equal to the nominal level (5%), and DESeq2 and DESeq2-ZINBWaVE have type I error ≤ 10% and FDR ≤ 20%. Interestingly, MGS, that had good control of type I error, has type I error greater than 10% in this specific setting.

### 3.5 Application

#### 3.5.1 DE analysis of the oral microbiome data in ZOE 2.0 study

We apply the methods that were shown to have reasonable sensitivity with a controlled type I error and FDR in simulations to the oral microbiome data in ZOE 2.0 study (“ZOE 2.0 data”). Because the ZOE 2.0 data have batch effects—significantly different sequencing depths between the two sequencing waves—we do not apply MGS, and hence, LB and LN tests are selected for the analysis. The data are normalized according to the TPM format with an average scale of 20 million. Differences in expression levels in TPM for each gene between health (non-ECC) and disease (ECC) participants are tested after controlling for batch effects and age (coded in months). The data set includes 297 children of ages between 36-71 months (3–5 years old). There are 402,937 genes from 204 bacterial species. Genes with *<* 10% prevalence and average TPM *<* 0.2 were excluded from the analysis, resulting in 157,113 genes in the final analysis data set. For the LN test, the minimum positive value (0.007) is uniformly added to the TPM values. Because this application is for illustration purposes, we did not apply a multiple testing correction and report only crude p-values. For each gene in LB tests, two Wald test p-values of the disease effects for the two parts, as well as the global p-value, are reported.

The existence of DE genes (among 157,113 genes) associated with ECC was clearly suggested by the observed high peak on the left side of histogram of the nominal p values (Figure10A for LN; C for LB). The higher peak observed in LN tests (Figure10 A) than LB tests (Figure10 C) implies that there are more genes found to be statistically significant by the LN model than by the LB model. The discrete (logistic) model part yields more significant results than the continuous (beta) model part in LB tests (Figure10B), indicating that the significance of the Wald test statistic (Figure10C) is mostly driven by the discrete model part. We observed a weak relationship between the significant coefficients of the two model parts in LB (Figure10D), as the positive slope in the scatter plot between the two coefficients indicates the sample group with higher non-zero average (*μ*) of one gene also tends to having higher non-zero proportion (1 − *π*) of the same gene, where positive coefficients imply higher nonzero mean (*μ*) comes with higher nonzero proportion (1 − *π*), respectively. We also find that (1) there are no genes that are significantly DE in both parts and have opposite/conflict DE directions between the two parts of LB, and that (2) the significant DE genes based on the global p value of LB have same-sign coefficients in both parts. However for the globally significant DE genes, the coefficients of the continuous model part are close to zero, suggesting again that the significance is driven by the discrete model part. This weak relationship may strengthen or weaken the justification for the Wald statistic. In rare events where the signals from the two parts are both strong and with opposite signs, the Wald statistic can detect the signals that would have vanished if the two effects were marginalized. On the other hand, when the signal from only one part is strong, while the other is not, the Wald statistic may not be able to detect the strong signal after being diluted by the weak one, resulting to low sensitivity. In this case, using the minimum p-values from both parts with an adjusted significance level, i.e., twice the nominal level for the Bonferroni-type adjustment, could be an alternative strategy.

**Fig. 10:**
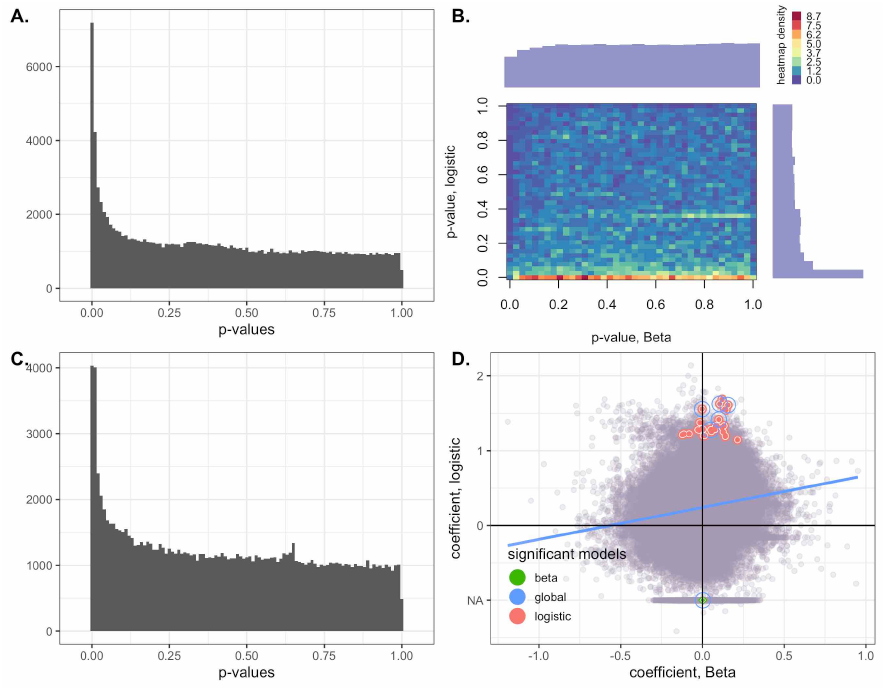
Application to the ZOE 2.0 data analysis results. A. Histogram of the p-values of the log-normal models. B. Histogram of the joint p-values of the logistic Beta models (logistic and Beta parts). C. Histogram of the single p-values of the logistic Beta models (Wald test statistics). D. Scatter plot of the coefficients of the LB models, with the circled dots representing the most significant genes—Wald test statistic *p <* 10^−5^ for the three types of Wald tests. Blue is for the global test, red is for the logistic part, and green is for the beta part. NA on the y-axis indicates that the logistic part was not estimated.

The number of significant DE genes (*p <* 10^−5^) are summarized in Figure 11. There are more number of significant genes according to the LN test (184) than the LB test (6 for the global test, 30 for the discrete part, and 1 for the continuous part). This is congruent with the fact that the LN test is more sensitive than the LB test under D4 (the differential disease effects in zero proportion) in Supplementary Figure 9. Most of the significant DE genes in the LB models are also reported as significant in the LN models. The ten genes with the lowest p-values from the LN models are: C8PIH7, C8PI10, C8PHV7, C8PEV7, C8PKZ2, C8PJY1, C8PG93, C8PKG9, C8PH26, and C8PJD1. The significant DE genes according to the LB Wald test were E0DI62, C8PHV7, C8PEV7, C8PI10, C8PIH7, and C8PHV8. The species and proteins associated with those genes and their functions listed in UniProt are present in Supplementary Table 11.

**Fig. 11:**
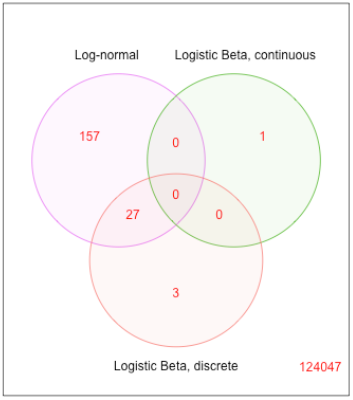
Venn diagram of genes with p-values are less than 10^−5^ for each evaluated model in the ZOE 2.0 data.

The results for the gene-species joint data and the results for the species marginal data are provided in Supplementary Sections 6. The patterns are overall similar to those obtained in the gene marginal data analysis; hiked frequency at the low p-value areas for the LN and global LB tests, significance mostly comes from the discrete part than the continuous part, the directions of the two parts in the significant taxonomic units are only weakly consistent with each other. The significant features (*p <* 10^−5^) in the gene-species joint data analysis using the LB models are E0DJ07 in Corynebacterium matruchotii; C8PHV7, C8PHV8, and C8PEV7 in Campylobacter gracilis; A3CQN5 in Streptococcus cristatus; G1WEB2 in Prevotella oulorum; and C7NCB2 in Leptotrichia shahii. It is noteworthy that the species information to annotate the gene-species level data may not be accurate in HUMAnN2 for oral bacterial species due to the built-in reference genome, e,g., for Leptotrichia and Streptococcus. Therefore, we further used species information in UniProt of the genes to annotate gene-species (Supplementary Table 11). The ten most significant gene-species from the LN tests were all associated with Campylobacter gracili and they include C8PHV7, C8PHV8, C8PEV7, C8PKG9, C8PI10, C8PH26, C8PHR6, C8PIH7, C8PFD0, and C8PG15 (Supplementary Table 11). The significant (*p <* 0.01) taxa from the species marginal data analysis using the LB models include Campylobacter gracilis, Streptococcus cristatus, Leptotrichia hofstadii, Lachnoanaerobaculum saburreum, Leptotrichia shahii, Streptococcus mutans, Campylobacter concisus, Prevotella oulorum (Supplementary Table 11). None of the species had *p <* 0.01 in the LN tests.

Streptococcus mutans, one of the identified significant DE species, is the most well-document dental caries-associated pathogen. The species most strongly associated with childhood dental caries in this analysis was Campylobacter gracilis, a gram-negative anaerobic bacillus, traditionally isolated from gingival crevices and dental biofilms accumulated close to the gingival margin (Vandamme et al., 1995). Oral campylobacters are enriched in genes for lactate metabolism, which plays an important role in the development and maintenance of acidic conditions in cariogenic biofilms as the predominant glucose-derived product, which is considered to be the main acid involved in caries formation (Iraola et al., 2014). The capacity of Campylobacter species to produce lactate may be contributing to the development and establishment of early childhood caries, as other microorganisms directly associated to caries disease like Streptococcus sp and Leptotrichia sp, which are benefited by this lactate-rich environment (Iraola et al., 2014; McLean et al., 2012). Chalmers et al. (2015) showed that Campylobacters gracilis is associated with severe early childhood caries at a frequency detection rate of 87.5% (Chalmers et al., 2015). Campylobacters gracilis’ active genes shown to have a significant association with ECC were associated with essential steps for: (1) bacterial growth (C8PIH7, encodes for an enzyme that catalyzes the first committed step in fatty acid synthesis) (Freiberg et al., 2006); (2) protein biosynthesis and transport (C8PI10, encodes for an enzyme that catalyzes the attachment of serine to its cognate transfer RNA molecule; C8PKZ2, encodes for the enzyme from biosynthesis of diverse amino acids leading to Llysine, Lthreonine, Lmethionine and Lisoleucine; C8PJD1, encodes for an amino acid biosynthesis pathway) (Cox et al., 2002; Tadrowski et al., 2016); (3) protein transport (C8PG93 encodes for twin-arginine translocation (Tat) pathway, which catalyzes the export of proteins from the cytoplasm across the inner/cytoplasmic membrane.) (Stephenson, 2005); (4) DNA replication and transcription (C8PJY1, encodes for key enzymes in the synthesis of nucleoside triphosphates molecular precursors of both DNA and RNA) (Chargaff, 2012); (5) biofilm formation or adhesion through gene C8PKG9 (encodes for NFACT-R 1 domain-containing protein) (Burroughs and Aravind, 2014) and; (6) energy conservation (C8PHR6 encodes for methylenetetrahydrofolate reductase (MTHFR) of acetogenic bacteria during reduction of carbon dioxide with molecular hydrogen to acetate) (Bertsch et al., 2015). Other genes associated with ECC were A3CQN5 (encodes for Ribosomal RNA small subunit methyltransferase A, which play the role of switch proteins in the ribosome assembly in Streptococcus sanguinis) and C7NCB2, which encodes for a multidrug and toxic compound extrusion (MATE) family of efflux pumps to actively transport of a solute across the membrane in Leptotrichia buccalis (Saier, 2000).

#### 3.5.2 DE analysis of the gut microbiome data in the IBD study

Next, we apply the LB and LN tests to the gut microbiome data in the IBD study (“IBD data”) to identify the differentially expressed genes associated with IBD. Out of 1,119,472 genes, 103,966 genes with prevalence rate *>* 0.1 and mean expression level *>* 10^−8^ in the relative RPKs were tested. Differences in expression levels in TPM for each gene between control (non-IBD, 26 (25%) participants) and cases (IBD, 78 (75%) participants) groups are tested after controlling for batch effects (binary-coded) and sex. The data set includes 104 patients (52 male and 52 female) who were between 5 and 74 years old at the time of diagnosis (for the cases). For the LN test, the minimum positive value (5 *×* 10^−10^) is uniformly added to the TPM values.

The p-values of each of the 103,966 genes for the LN and LB tests are summarized in Figure 12. Similarly to the ZOE data analysis results, the hike on the left end in both LB (Figure 12A) and LN (Figure 12C) indicates the existence of differentially expressed genes. However, in Figure 12B, only the continuous part has a conspicuous hike while the discrete part is mostly flat, implying that the signal lies massively in the continuous part, which is also confirmed in the scatter plot (Figure 12D). It is noteworthy that about 4% of genes have very high discrete model coefficients but are insignificant in two clusters around either 26 and −26 on the y-axis. These are a manifestation of the undesirable feature of Wald statistics called the Hauck-Donner effects, where larger disease effects may not always result in a larger statistic and, as a consequence, may yield lower sensitivity (Hauck Jr and Donner, 1977). This occurs when a phenotype group has prevalence rate of exactly zero or one, while the other group has prevalence rate away from zero and one. The likelihood-ratio test, permutation tests, Fisher’s exact test, regularization, and Bayesian approaches are the alternatives to the Wald test. Among the 523 candidate genes with such prevalence rate pattern, no genes were found significant at the significance level of the nominal P value 10^−5^ by the likelihood-ratio test and the Fisher’s exact test of which p-values are presented in Supplementary Figure 24.

**Fig. 12:**
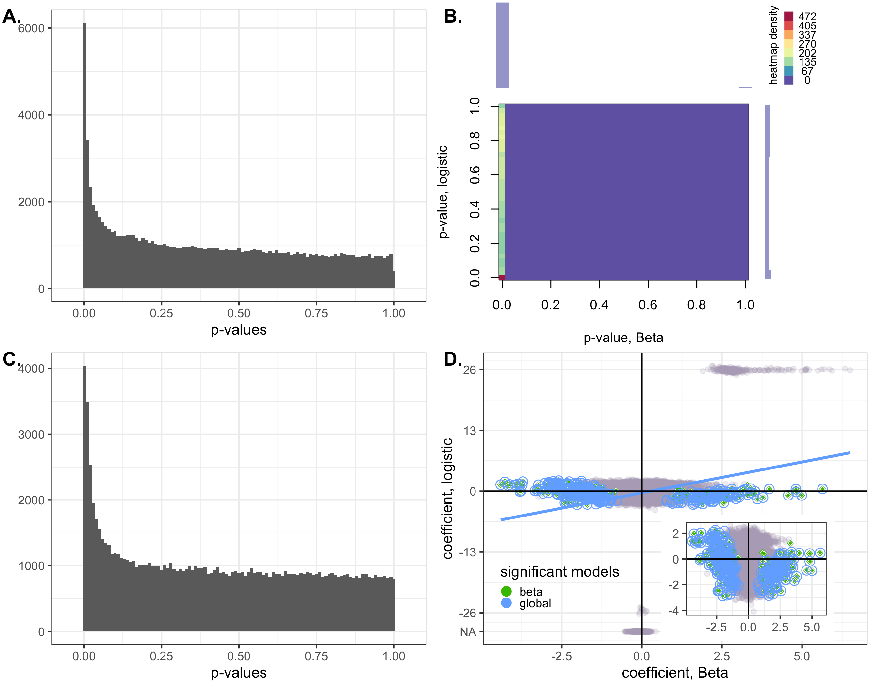
Application to the IBD data analysis results. A. Histogram of the p-values of the log-normal models. B. Histogram of the joint p-values of the logistic Beta models (logistic and Beta parts). C. Histogram of the single p-values of the logistic Beta models (Wald test statistics). D. Scatter plot of the coefficients of the LB models, with the circled dots representing the most significant genes—Wald test statistic *p <* 10^−5^. The NA results around *y* = *±*26 indicate that the logistic part was not estimated.

The numbers of the signficant genes in LN, LB-continuous, and LB-discrete models are given in Figure 13. The ten most statistically significant genes in the LN model are S3BI82, R5Q3H7, R5PRG3, R5Q1H1, R5QAG2, R5QE55, S3CE88, R5QEQ4, R5PLJ0, and, G2T243 while the top ten genes for the LB model are D4WIY6, Q0TKG5, R6W6W2, D1PDG3, Q17UW4, I9USK4, E2ZM16, R7NP61, B0NN15, and U2ZZD9. The species and proteins associated with those genes and their functions are presented in Supplementary Table 12

**Fig. 13:**
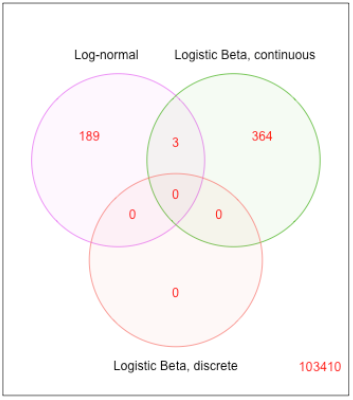
Venn diagram of genes with p-values are less than 10^−5^ for each evaluated model in the IBD data.

For the top 10 gene families identified as significant by the LN model, the first nine corresponded to Sutterella wadswothensis, Gram-negative, non-spore-forming rods of the Betaproteobacteria class that grow in a microaerophilic atmosphere or under anaerobic conditions, and which has been previously identified as significantly differentially abundant (prior to the correction for multiple hypothesis testing) in the original study (Lloyd-Price et al., 2019). The role of S. wadworthensis in IBD pathogenesis remains unresolved with studies both supporting (Gryaznova et al., 2021; Douglas et al., 2018) and not supporting this role (Mukhopadhya et al., 2011; Hiippala et al., 2016). The most significant gene family identified by the LN model, S3BI82, corresponded to the hydrogenase maturation factor HypA, a Ni-metallochaperone that binds to a Ni ion with micromolar affinity aiding in the assembly of [NiFe] hydrogenases, enzymes that catalyze the reversible hydrogen production and consumption of hydrogen. Hydrogenases are important factors that contribute to the metabolic versatility of Enterobacteriaceae, specifically their ability to utilize a large repertoire of terminal electron acceptors, which allows them to thrive in the inflamed gut (Hughes et al., 2021; Nguyen et al., 2020; Hughes et al., 2017).

The final significant gene family in the top 10 indicated by the LN model was a stage 0 sporulation protein A homolog found in Roseburia hominis. R. hominis, a prevalent butyrate producer, has been found consistently depleted in CD (Machiels et al., 2014; Patterson et al., 2017) and also identified as significant prior to the multiple hypothesis correction in the original study (Lloyd-Price et al., 2019). The 11th through 20th significant gene families assessed by the LN model included only a single additional genus, Alistipes, a member of the Bacteroidetes phylum. finegoldii is considered a protective species against colitis since its abundance was lower in mice with colitis, and reduced symptoms of colitis when administered with Prevotella falsenii, Bacteroides eggerthii, and Parabacteroides distasonis. Alistipes shahii was identified as significant prior to multiple hypothesis correction in the original study (Lloyd-Price et al., 2019).

The 10 gene families identified as significantly over or under represented by the LB model corresponded to Bacteroides ovatus, Escherichia coli, Prevotella copri, Faecalibacterium prausnitzii, Bacteroides faecis, Bacteroides xylanisolvens, and Bacteroides stercoris; all identified as significant in the original study (Lloyd-Price et al., 2019) prior to multiple hypothesis correction. Although the species Zygosaccharomyces rouxii harboring to Q17UW4 was not reported in the original study (Lloyd-Price et al., 2019), the gene family itself corresponded to a Cytochrome c oxidase subunit 2 which has been observed to increase significantly in UC (El Sayed and Sayed, 2019). Finally, the relevance of Faecalibacterium praunitzii as an important beneficial bacterium in IBD has been demonstrated in a number of studies (Sokol et al., 2008), together with the butyrate-producing R. hominis have been observed in the original stuy (Lloyd-Price et al., 2019) accounting for some of the strongest associations overall.

## 4 Discussion

In this study, we have provided a comprehensive evaluation of the main analysis methods for differential gene expression of metatranscriptomics data. This simulation study design is inspired by the human oral microbiome sequencing data, to which we investigated the goodness of fit of the generative models after scaling or transformation. The methods were evaluated in terms of control of type I error, sensitivity, and FDR. The semi-parametric simulations using both the oral and gut microbiome sequencing data were further performed as a complement of the parametric simulations. The best-performing methods were further used for detecting the differentially expressed genes in the ZOE 2.0 oral metatranscriptomics data and the IBD gut metatranscriptomics data. The microbial genes found significantly associated with ECC were reported and interpreted accordingly.

The simulation study offers guidance to microbiome investigators for choosing appropriate DE analysis methods. Our simulation framework could be further applied for the purposes of validating current and future DE methods that were not included in this manuscript. In what follows, we summarize the main findings and discuss the limitations of the simulations.

### Which method is the best in general?

In Table 2, we summarize the performance of each method evaluated in this study. The simulation results suggest that for metatranscriptomics data, LB and MGS have good sensitivity and good control of type I error under the scenarios that involve *μ*-differences (D2 and D6). However, the current version of MGS does not control for batch effects and MGS does not have good sensitivity under D4 (*π*-differences) and LB needs to be used with caution as it may have inflated type I error for high zero proportion with a small sample size. LN has a decent level of sensitivity when there is non-zero mean difference (for low baseline *π* values) or zero-proportion difference, or when both differences are present with the opposite directions. Both MAST and ANCOM-BC (both versions) have high power to detect non-zero mean differences, marginal mean differences, and the combinations of the two under many scenarios, but they do not properly control type I error. DESeq2 without ZINBWave has both low type I error and sensitivity for metatranscriptomics data, and the one with ZINBWave shows unstable control of type I error. KW as a nonparametric test, has generally lower sensitivity compared to other methods. As simulations are based on parametric generative models, the low sensitivity of KW is somewhat expected, but KW can still be considered when the distribution of the data at hand are thought to differ from the model that other tests assume to a great degree. LEfSe, which does not provide *p*-value, needs to be used with carefully selected thresholds to avoid inflated type-I error.

**Table 2.** Summary of performances. DESeq2ZI is the abbreviation of DESeq2-ZINBWaVE. Sensitivity (I) is sensitivity in parametric simulations; Sensitivity (II) is sensitivity in semi-parametric simulations. ○ models zero counts and considers the differential effects on zeros in the tests. △ models zero counts but the tests only the marginal or non-zero mean differences. × does not model zeros.

| method | type I error & FDR | sensitivity (I) | sensitivity (II) | zero model | other |
| --- | --- | --- | --- | --- | --- |
| LB | fair | high (D2, D4, D6, D8) | high | ○ | Inflated type I error for high $\pi$ and small $n$ in D2. |
| MGS | good | high (D2, D6) | high | △ | No batch control available. |
| LN | good | good (D4, D6) | mid-high | × | Low-powered for high $\pi$ in D2. |
| MAST | not controlled | high | high | ○ | type I error not controlled. |
| ANCOM-BC | not controlled | good (D4, D6) | mid-high | △ | type I error not controlled. |
| DESeq2 | good | low | mid-high | × |  |
| DESeq2ZI | unstable | good | high | △ | type I error not controlled |
| LEfSe | not controlled | good | low | × | $p$-value is not provided. |
| ALDEx2 | good | good (high $\mu$) | low | × | Fine tuning of the scale ($\mu$) is required. |
| KW | good | good (D4, D6) | low | × |  |
| KW2 | good | good (low $\pi$) | high | ○ | Low-powered when $\pi \geq 90\%$. |

### TPM transformation

According to our simulations, the log-normal distribution has a decent goodness-of-fit to the ZOE data after TPM transformation. This is consistent with the fact that MGS, which assumes zero-inflated log-normal distribution, is one of the most sensitive methods.

### Two-part models are beneficial in some cases

Two-part models have advantages in sensitivity over single-part ones when the signals come from two different sources with different directions (D8). Specifically this occurs when the non-zero mean difference and the difference in zero-proportion (or zero-inflation probability for ZINB model) both exist and they have the same sign (or the opposite directions in terms of the marginal mean). Even KW-II, not the highest-powered test under D2 or D4, often performs better than other single-part models when both signals are present.

### Distinct generative models

Some of the variational factors in this simulation may not have practical implications: e.g., generative models and batch effects. Simulation results showed that there were no substantial differences in sensitivity between different generative models, even if each generative model might have a different most powerful test. This might be due to the fact that all the generative distributions considered in this paper, ZILN, ZIG, and ZINB, have similar features such as zero-inflation and left-skewed unimodal distributions in their non-zero parts. Batch effect scenarios (K2 to K5) did not appreciably affect sensitivity even when the models were not correctly specified; for instance, the disease and batch effects were misspecified in the sense that independently generated multiplicative effects in *μ* of disease and batches resulted in interaction effects, whereas, the testing models assumed no interaction effects in this study.

### Limitation: distributional assumptions

The parametric simulation deals with a sizable number of distinct scenarios. However, it does not cover all possible data generative mechanisms. It is based on a combination of a few parametric generative models and a limited number of parameter sets. For example, the differential expression is based on fixed functions such as log-difference and logit-difference, we did not consider (multiplicative) interactions between disease effects and batch effects, and genes’ expressions were generated independently from each other. Each of these issues adds a chance that the simulation results may not plausibly represent the true data distributions in real-life experiments. However, we believe that this simulation results provide useful and practical insights regarding the behavior and performance of each test under certain settings, if the data are not too different from the models considered in this simulation. This is supported in part by the fact that the conclusions from the semi-parametric simulations were congruent with the parametric simulation results.

### Limitation: gene independence assumptions

It must be acknowledged that we assumed independence between genes in the parametric simulations. Although in reality, it is likely that some dependency or co-expression of genes is at play and it may affect the significance of potential gene set tests (Wu and Smyth, 2012) or multiple testing adjustment, such dependency would not affect the differential expression analysis at the individual gene level. Furthermore, all non-collective testing methods and the multiple testing procedures assume independence between genes, where the collective testing methods include MAST, MGS, and DESeq2, and p-values are calculated taking into account the dependence between genes using empirical Bayes. In other words, for each gene, the *p*-values obtained by individual gene tests on independently simulated data are valid; and so are the corresponding type I and type II error rates. Although, in real data settings, genes are dependent to some degree, we expect that, the high dimensionality of the data used here (i.e., a large number of simulated genes) likely introduced spurious correlations between features (Fan et al., 2014), our simulation results may not be far from a realistic scenario. Future work could explore more realistic settings where gene expression levels are correlated.

## Supporting information

All supplementary files

## Supplementary Materials

Further details of differential expression analysis methods and data generative models are provided in Supplementary Sections 1–2, respectively. Supplementary Section 3 entails supplementary information about the parametric simulation set-ups, and Supplementary Section 4 includes the goodness of fit results for the ZINB model. Extensive simulation results are provided in Supplementary Section 5. Additional results of methods’ application to the ZOE 2.0 data and the IBD data are contained in Supplementary Sections 6 and 7, respectively.

## Key Points

- Microbiome has emerged as an undeniable cornerstone for a multitude of health and disease outcomes. Besides metagenomics data, metatranscriptomics provides the opportunity to measure the activity of genes, instead of inferring gene expression from microbial genomes.
- How to choose the method for Differential Expression (DE) analysis at the gene level in metagenomics data is a key question, accounting for the high percentage of zeros, overdispersion, possibly compositional data structures. Sample size, existence of covariates or the batch variable, and the data distribution all affect the choice of DE analysis methods.
- We evaluate 10 methods with focus on statistical models that were specifically designed for zero-inflated over-dispersed counts or compositional data, and particularly the DE analysis at the more sparse and challenging gene level instead of the species level. The 10 methods are log-normal (LN), logistic-beta (LB), MAST, DESeq2, metagenomeSeq, ANCOM-BC, LEfSe, ALDEx2, Kruskal-Wallis, and two-part Kruskal-Wallis. We also discussed the comparison of normalization methods.
- We undertook a comprehensive evaluation and benchmarking of ten differential analysis methods for metatranscriptomics data. We used a combination of real and simulated data to evaluate performance (i.e., model fit, type I error, false discovery rate, and sensitivity). Though different methods show different advantages and limitations in different scenarios, logistic repression has a good well-balanced performance.
- The large scale of metagenomics data have been used, from either the oral health ECC studies or the gut health IBD studies.

## Data Availability

The real datasets analyzed in this manuscript are publicly available. ZOE 2.0 data are available in the dbGaP repository https://www.ncbi.nlm.nih.gov/gap under the umbrella study name Trans-Omics for Precision Dentistry and Early Childhood Caries or TOPDECC (accession: phs002232.v1.p1), and the IBD data are available at https://ibdmdb.org. The code for the procedure is available at https://github.com/Hunyong/microbiome2020/blob/master/Readme_KS_test.Rmd.

## Acknowledgements

None at this time.

