## Supplementary material for "Comprehensive evaluation of methods for differential expression analysis of metatranscriptomics data": All supplementary files

December 1, 2022

#### 1 Differential expression analysis methods

In this section, we present each of the DE analysis methods in detail. We first introduce notational conventions. Throughout this material, $Y_{i,g}$ denotes the expression level for the $g$th gene in the $i$th cell, $X_i \equiv (1, X_i^D, X_i^B)$ denotes the $i$th row of the design matrix, or a vector containing the intercept term, a binary disease status, and a binary batch indicator. Different models abusively use the same notation for parameters, as long as there is no confusion; e.g. regression coefficients $\beta_g$ are commonly used either in the LN model or the LB model, but they are shorthand for $\beta_g^{LN}$ and $\beta_g^{LB}$, respectively.

##### 1.1 Log-normal test

The Log-normal (LN) test relies on the assumption that the log-transformed expression is normally distributed as in (1). A small positive constant ($c$) is added to the gene expression to ensure that the log-transformed values are within a feasible range. In this simulation study 1 is uniformly added to expression levels ($c = 1$).

$$\log_2(Y_{i,g} + c) \sim N(\mu_{i,g}, \sigma_g), \tag{1}$$

where $\mu_{i,g} \equiv X_i^\top \beta_g$ with $\beta_g \equiv (\beta_g^0, \beta_g^D, \beta_g^B)^\top$.

The null and the alternative hypotheses for the $g$th gene are

- $H_0$: $\beta_g^D = 0$ and
- $H_1$: $\beta_g^D \neq 0$.

The test statistic for the $g$th gene is $T_g^{LN} = \left(\frac{\hat\beta_g^D}{se(\hat\beta_g^D)}\right)^2$ and follows a $\chi_1^2$ distribution under the null hypothesis asymptotically. The test rejects the null hypothesis if the test statistic is larger than $\chi_1^2(1-\alpha)$, or the $(1-\alpha)$th quantile of the $\chi^2$ distribution with one degree of freedom, where $\alpha$ is the significance level. Alternatively, the individual $p$-values are obtained as $p_g = 1 - F_{\chi_1^2}(T_g^{LN})$, where $F_d(t)$ is the distribution function of $d$ evaluated at $t$. The genes with $p$-values less than $\alpha$ are declared to have a statistically significant association with disease. This test is simply an analysis of covariance (ANCOVA) with an appropriately log-transformed dependent variable, and is easily implemented in most statistical software packages. The testing procedure, after obtaining a test statistic and the corresponding null distribution (e.g., p-values and rejection regions), is identical for rest of the methods, hence will be omitted unless needed.

#### 1.2 Logistic Beta test

The Logistic Beta model (LB) models relative expressions, $R_{i,g} \equiv Y_{i,g}/\sum_{h=1}^{G} Y_{i,h}$, instead of absolute expressions, $Y_{i,g}$. Because of the sum-to-one constraint of relative expressions, tests based on relative expression are structurally dependent. However, in microbiome data analyses, the number of tested genes is usually large enough and thus the dependence induced by the compositional structure is negligible.

The LB model is formulated (Peng *et al.*, 2016) as:

$$R_{i,g} \sim LB(\pi_{i,g}, \mu_{i,g}, \phi_g), \tag{2}$$

where $\pi_{i,g} = \text{expit}(X_i^\top \gamma_g)$ with $\gamma_g = (\gamma_g^0, \gamma_g^D, \gamma_g^B)^\top$, $\mu_{i,g} = \text{expit}(X_i^\top \beta_g)$ with $\beta_g \equiv (\beta_g^0, \beta_g^D, \beta_g^B)^\top$, $\phi_g$ denotes the dispersion parameter such that $var(R_{i,g}|R_{i,g} > 0) = \mu_{i,g}(1-\mu_{i,g})\phi_g$, and $\text{expit}(\cdot) := \frac{\exp(\cdot)}{\exp(\cdot)+1}$.

Note that this model can be decomposed into two orthogonal models:

$$1(R_{i,g} = 0) \sim \text{Bernoulli}(\pi_{i,g}), R_{i,g}|R_{i,g} > 0 \sim \text{Beta}(\mu_{i,g}, \theta_g), \tag{3}$$

where $1(\cdot)$ is the indicator function, $\mu_{i,g}$ is the mean of the Beta random variable and $\theta_g$ is the dispersion parameter. Orthogonality means that the estimate of $\pi_{i,g}$ and those of $\mu_{i,g}$ and $\theta_g$ are independent. Consequently, the test statistic can be obtained from these two separately estimated models. The maximum likelihood estimators (MLE) are used for estimation and an `R` package `gamlss` (Stasinopoulos and Rigby, 2007) was used for simulation in this study.

The null and the alternative hypotheses for the $g$th gene are

- $H_0$: $\beta_g^D = \gamma_g^D = 0$ and
- $H_1$: Either $\beta_g^D \neq 0$ or $\gamma_g^D \neq 0$.

Either a Wald-type or a likelihood test statistic can be used to test these hypotheses. Because they are asymptotically equivalent, here we only present a Wald test statistic:

$$T_g^{LB} = \left(\frac{\hat{\beta}_g^D}{se(\hat{\beta}_g^D)}\right)^2 + \left(\frac{\hat{\gamma}_g^D}{se(\hat{\gamma}_g^D)}\right)^2, \tag{4}$$

which follows a $\chi_2^2$ distribution under the null hypothesis asymptotically.

If only one of the two parts of LB is estimable, the test statistic is constructed based on the estimable component only and the reference distribution is $\chi_1^2$; e.g. when only the logistic model is estimable, $T_g^{LB} = \left(\frac{\hat{\gamma}_g^D}{se(\hat{\gamma}_g^D)}\right)^2$. The same approach was followed for the other two-part tests, including MAST and the two-part Kruskal-Wallis test.

#### 1.3 MAST

The"Model-based Analysis of Single-cell Transcriptomics" (MAST) (Finak *et al.*, 2015) was proposed specifically for differential expression analysis of scRNAseq data. This model, composed of a logistic regression model and a conditional log-normal model, regularizes parameter estimation and utilizes estimated cellular detection rates (CDR) as covariates as defined below. The model was designed to deal with zero-inflation which is driven by both technical and biological variabilities in scRNAseq data. Though zeros in microbiome sequencing data are believed to be generated mostly by biological reasons, the proportion of zeros is usually greater than that of conventional single-part parametric models such as Poisson, negative binomial, and log-normal. Thus, it is feasible to interrogate the performance of MAST in the context of microbiomal transcriptomics analysis.

The models in MAST can be summarized as

$$1(Y_{i,g} = 0) \sim \text{Bernoulli}(\pi_{i,g}), \log_2(Y_{i,g} + 1) \mid Y_{i,g} > 0 \sim N(\mu_{i,g}, \sigma_g^2), \tag{5}$$

where $\pi_{i,g} = \text{expit}(X_i^\top \gamma_g)$ with $\gamma_g \equiv (\gamma_g^0, \gamma_g^D, \gamma_g^B, \gamma_g^C)^\top$, $\mu_{i,g} = (X_i^\top \beta_g)$ with $\beta_g \equiv (\beta_g^0, \beta_g^D, \beta_g^B, \beta_g^C)^\top$, $X_i \equiv (1, X_i^D, X_i^B, X_i^C)$, and $X_i^C = \frac{1}{n}\sum_{i=1}^{n} 1(Y_{i,g} > k)$ is the CDR of the $i$th subject for background expression level $k$. In this simulation we set $k = 0$.

The parameters are estimated using a Bayesian framework, where $\gamma_g$ is regularized under weak informative prior and $1/\sigma_g$ is regularized using empirical Gamma prior. An `R` package `mast` is available (McDavid *et al.*, 2019).

The null and the alternative hypotheses for the $g$th gene are

- $H_0$: $\beta_g^D = \gamma_g^D = 0$ and
- $H_1$: Either $\beta_g^D \neq 0$ or $\gamma_g^D \neq 0$.

Either a Wald-type or a likelihood test statistic can be used to test these hypotheses. The Wald statistic is

$$T_g^{MAST} = \left(\frac{\hat{\beta}_g^D}{se(\hat{\beta}_g^D)}\right)^2 + \left(\frac{\hat{\gamma}_g^D}{se(\hat{\gamma}_g^D)}\right)^2, \quad (6)$$

with $\chi_2^2$ as its asymptotic null distribution. The testing procedure is exactly the same as that of the LB test once the coefficients and their standard errors are estimated.

#### 1.4 DESeq2

The DESeq2 (Love *et al.*, 2014) method is currently widely used for differential expression of RNAseq data. The underlying model of DESeq2 is a negative binomial distribution and it uses empirical Bayes for regularization.

The DESeq2 model can be summarized as

$$Y_{i,g} \sim NB(\mu_{i,g}, \theta_g), \quad (7)$$

where $\mu_{i,g} = s_{i,g}\nu_{i,g}$ is the mean parameter, $\theta_g$ is the dispersion parameter, $s_{i,g}$ is the size factor, $\nu_{i,g} = \exp(X_i^\top \beta_g)$ with $\beta_g \equiv (\beta_g^0, \beta_g^D, \beta_g^B, \beta_g^C)^\top$, and $X_i \equiv (1, X_i^D, X_i^B, X_i^C)$. The size factor is the parameter with which we adjust the sequencing depth. In this simulations we use the median-of-ratios method (Anders and Huber, 2010).

The parameters are estimated using maximum likelihood estimation and then $\theta_g$ and $\beta_g^D$ are regularized using an empirical Bayes approach. An `R` package `DESeq2` is available.

The null and the alternative hypotheses for the $g$th gene are

- $H_0$: $\beta_g^D = \gamma_g^D = 0$ and
- $H_1$: Either $\beta_g^D \neq 0$ or $\gamma_g^D \neq 0$.

A Wald test is used to test these hypotheses. The Wald statistic is given as

$$T_g^{DESeq2} = \left(\frac{\hat{\beta}_g^D}{se(\hat{\beta}_g^D)}\right)^2, \quad (8)$$

with $\chi_1^2$ as its asymptotic null distribution. The testing procedure is exactly the same as that of LB test, once the coefficients and their standard errors are estimated.

Because DESeq2 cannot accomodate high zero proportions, an extension was recently developed to enable the modeling of a greater number of zeros in the scRNAseq context (Van den Berge *et al.*, 2018). In this modified DESeq2 method, namely DESeq2-ZINBWaVE, first the zero-inflation parameter is estimated using the model,

$$Y_{i,g} \sim ZINB(\mu_{i,g}, \theta_g, \pi_{i,g}), \quad (9)$$

and each observation is assigned a weight of the posterior probability of non-zero-inflation,

$$\frac{(1-\pi_{i,g}) f_{ZINB}(y_{i,g}; \mu_{i,g}, \theta_g, 0)}{f_{ZINB}(y_{i,g}; \mu_{i,g}, \theta_g, \pi_{i,g})},$$

where $f_{ZINB}$ is the corresponding density of the ZINB distribution. For the size factor estimation in DESeq2-ZINBWaVE, we use the positive counts method. Then the conventional DESeq2 method is applied, as described earlier, including the weights. Whenever there is no ambiguity, “DESeq2” refers to the original method and “DESeq2-ZINBWaVE” to its extension.

#### 1.5 metagenomeSeq

MetagenomeSeq (MGS) is a differential abundance analysis method for metagenomics data (Paulson *et al.*, 2013a) and the corresponding bioconductor package, `metagenomeSeq` is available. MGS assumes zero-inflated log normal distribution. Furthermore, MGS uses an empirical Bayes shrinkage method for parameter estimation. Hence, MGS shares common modeling approaches with MAST; however, the two approaches are different in a few aspects. First, MAST uses CDR as a controlling factor in the model while MGS does not. Second, MAST provides tests on two parts of the model; i.e., two p-values are obtained from the zero-inflation part and the log-normal part in MAST. However, in MGS, after estimating the two-part model parameters, only the log-difference of the marginal mean is tested and a single p-value is given (Paulson, 2015). In our simulations, the `fitFeatureModel` function in the `R` package `metagenomeSeq` is used for implementation (Paulson *et al.*, 2013b). Although the MGS test can account for batch effects mathematically, the current `metagenomeSeq` software does not allow batch variables in the model. Thus, only results without batch effects will be reported in the simulation study in Section 4.

#### 1.6 ANCOM-BC

ANCOM-BC is a differential abundance analysis method for metagenomics data (Lin and Peddada, 2020). It shares the philosophy of its predecessor, ANCOM (Mandal *et al.*, 2015), in that it models the ratio of abundances between taxa. However, unlike ANCOM which is a rank-based approach, ANCOM-BC specifies the test statistic and its associated $p$-value for a large sample. In ANCOM-BC, the observed abundance $Y_{i,g}$ is assumed to be a realization of the unknown abundance $U_{i,g}$ of the whole ecosystem from where the sample is taken with possibly different sampling fraction $\eta_i$ for each sample. In other words, $E[Y_{i,g}|U_{i,g}] = \eta_i U_{i,g}$, where $U_{i,g}$ is a random variable with mean $\theta_g^D$ or $\theta_g^H$, depending on the membership of the sample $i$ to the disease ($D$) or health ($H$) group. Of note, ANCOM-BC is not limited to two-group problems but are designed for multi-group problems. Then it formulates $\log Y_{i,g} = \log \tilde{\eta}_i + \log \theta_{i,g} + \epsilon_{i,g}$, where $\tilde{\eta}_i$ is a slightly-redefined sampling fraction parameter due to the log-transformation, and $E[\epsilon_{i,g}] = 0$.

The hypotheses of ANCOM-BC are

- $H_0$: $\log \theta_g^D = \log \theta_g^H$ and
- $H_1$: $\log \theta_g^D \neq \log \theta_g^H$,

which are tested by the test statistic,

$$T_g^{ANCOM-BC} = \frac{\widehat{\log \theta_g^D} - \widehat{\log \theta_g^H} - \widehat{\log \tilde{\eta}}}{\sqrt{\{\hat{\sigma}_g^D\}^2 + \{\hat{\sigma}_g^H\}^2}},$$

where $\widehat{\log \theta_g^A}$ is the estimates of $\log \theta_g^A$, $\{\hat{\sigma}_g^A\}^2$ is the mean squared error for each group $A = D, H$, and $\widehat{\log \tilde{\eta}}$ is the estimate of the bias, $\log \tilde{\eta} \equiv E[\log \theta_g^D - \log \theta_g^H]$. The statistic follows the standard normal distribution per the large sample theory, and the authors defined a small sample version of the statistic of which distribution was not defined.

ANCOM-BC does a further procedure of detecting "the structural zero" which is defined to be the absence of a certain taxon in a specific group that is present in another group. Once the structural zero is detected, ANCOM-BC declares that the taxon is differentially abundant, giving $T_g^{ANCOM-BC} = \infty$ and $p$-value $= 0$. However, since such procedure often inflates the type-I error significantly in this simulation study, we add another version of ANCOM-BC that declares those sturctural zeros inconclusive (i.e., $p$-value = NA). We report the simulations results for second version as "ANCOM-BC2" and disclose the results of the original version, denoted as "ANCOM-BC1," in the Supplementary Materials.

#### 1.7 LEfSe

LEfSe (Linear discriminant analysis Effect Size) is commonly used for differential analysis of metagenomic biomarkers (Segata *et al.*, 2011). It assumes that the samples are labeled with a certain group that represents the main biological comparison class of interest. They may also include one or more subgroup labels that indicate within-group classifications where batch effects can be accounted for in our simulations. By combining standard tests for statistical significance, with extra tests encoding biological consistency and effect relevance, the method determines the features most likely to explain variations across groups.

There are three steps performed in order: the KW rank-sum test on groups, the pairwise Wilcoxon test between subgroups of different groups, and the LDA on the relevant features. In the first step, the factorial KW rank-sum test is applied to each feature according to the group label; the subgroup label is utilized for further stratifying when the information exists. As a result, only the features that reject the null hypothesis of identical value distribution among groups could be analyzed further. In the second step, the pairwise Wilcoxon test is applied to the extracted features belonging to subgroups of different groups, for testing whether all pairwise comparisons between subgroups in different groups significantly agree with the group level trend. If at least one comparison between subgroups has a p-value greater than the specified threshold, or if the sign of variation is not equal across all comparisons, the pairwise Wilcoxon test is not satisfied for that feature. The first two steps employ non-parametric tests because they are distribution-free approaches and much more robust to the underlying distribution of the data: the only assumption of the Wilcoxon and KW tests is that the distributions in each group are identically shaped with possible differences in the medians. Finally, in the third step, an LDA model is generated, by assigning the remaining features and subgroup labels as the independent variable and the group label as the dependent variables. This step is used to estimate their effect sizes, which are obtained by averaging the differences between group means with the differences between group means along the first linear discriminant axis, which equally weights features' variability and discriminatory power. The LDA score for each biomarker is obtained by computing the logarithm of this value after being scaled and induces the ranking of biomarker relevance, regardless of the absolute values of the LDA score. For robustness, LDA is additionally supported by bootstrapping and subsequent averaging. For implementation of this test, an R function lefser::lefser() is available.

#### 1.8 ALDEx2

ALDEx2 is a differential abundance analysis method for RNA-seq, 16S rRNA gene sequencing and differential growth datasets (Fernandes *et al.*, 2014). Instead of proposing new model, this method pays more attention on the data preprocessing, and it uses multiple instances to generate p-values.

If $Y_{i,g}$ denotes the read counts for the $g$th gene in the $i$th cell, suppose we want to use $K$ instances to generate the p-value. For each sample $i$, to generate the $k$th instance, ALDEx2 uses posterior Dirichlet distribution with an uninformative prior of $\frac{1}{2}$ to model the frequency of features with zero counts.

$$\left(Y_{i,1}^{(k)}, Y_{i,2}^{(k)}, \ldots, Y_{i,G}^{(k)}\right) = Dir\left(Y_{i,1} + \frac{1}{2}, Y_{i,2} + \frac{1}{2}, \ldots, Y_{i,G} + \frac{1}{2}\right) \tag{10}$$

where $Dir(\alpha_1, \alpha_2, \ldots, \alpha_G)$ denotes the Dirichlet distribution with paramter $\boldsymbol{\alpha} = (\alpha_1, \ldots, \alpha_K)$. Then, ALDEx2 uses CLR method to centralize the input data.

$$c_{i,g}^{(k)} = log_2\left(Y_{i,g}^{(k)}\right) - \frac{1}{G} log_2\left(\prod_{g=1}^{G} Y_{i,g}^{(k)}\right) \tag{11}$$

Then different models can be applied to the certralized data. Since we have two covariates in the model, we can use the linear model which is formulated as:

$$c_{i,g}^{(k)} \sim N\left(\mu_{i,g}^{(k)}, \sigma_g^{(k)}\right) \tag{12}$$

Where $\mu_{i,g} \equiv X_i^\top \boldsymbol{\beta}_g^{(k)}$ with $\beta_g^{(k)} \equiv \left(\beta_g^{0,(k)}, \beta_g^{D,(k)}, \beta_g^{B,(k)}\right)^\top$.

The null and the alternative hypotheses for the $g$th gene with $k$th instance is

- $H_0$: $\beta_g^{D,(k)} = 0$ and
- $H_1$: $\beta_g^{D,(k)} \neq 0$.

The test statistic for the $g$th gene with $k$th instances $T_g^{LN,(k)} = \left(\frac{\hat{\beta}_g^{D,(k)}}{se(\hat{\beta}_g^{D,(k)})}\right)^2$ and follows a $\chi_1^2$ distribution under the null hypothesis asymptotically. The test rejects the null hypothesis if the test statistic is larger than $\chi_1^2(1-\alpha)$, or the $(1-\alpha)$th quantile of the $\chi^2$ distribution with one degree of freedom, where $\alpha$ is the significance level. Alternatively, the individual $p$-values with $k$th instance are obtained as $p_g^{(k)} = 1 - F_{\chi_1^2}(T_g^{LN,(k)})$, where $F_d(t)$ is the distribution function of $d$ evaluated at $t$. The final p-value for gene $g$ is defined using $p_g = \frac{1}{K}\sum_{k=1}^{K} p_g^{(k)}$. And the genes with $p$-values less than $\alpha$ are declared to have a statistically significant association with disease.

#### 1.9 Kruskal-Wallis test

The Kruskal-Wallis (KW) test is equivalent to one-way analysis of variance (ANOVA) on ranks. KW is equivalent to Wilcoxon's rank sum (WRS) test, or Wilcoxon-Mann-Whitney test, for two sample problems and can accomodate comparisons of more than two samples (Kruskal and Wallis, 1952). Although the prototypical KW test was designed without consideration of covariates, it can be modified to account for possible batch effects (Hothorn *et al.*, 2006):

$$T_g^{KW} \equiv (n-1)\frac{\sum_{i=1}^{n}(\bar{r}_g^{db} - \bar{r}_g^{\cdot b})^2}{\sum_{i=1}^{n}(r_{i,g} - \bar{r}_g^{db})^2}, \tag{13}$$

where $r_{i,g} := \sum_{j=1}^{n}\{1(Y_{i,g} > Y_{j,g}) + \frac{1}{2}(Y_{i,g} = Y_{j,g})\} + 1$ is the rank of the $i$th subject's $g$th gene among $n$ subjects, $\bar{r}_g^{db} := \frac{\sum_{i=1}^{n} r_{i,g} 1(X_i^D = d, X_i^B = b)}{\sum_{i=1}^{n} 1(X_i^D = d, X_i^B = b)}$, $\bar{r}_g^{d\cdot} := \frac{\sum_{i=1}^{n} r_{i,g} 1(X_i^D = d)}{\sum_{i=1}^{n} 1(X_i^D = d)}$, and $\bar{r}_g^{\cdot\cdot} := \frac{1}{n}\sum_{i=1}^{n} r_{i,g}$.

The null and the alternative hypotheses for the $g$th gene are

- $H_0$: The ranked expression levels are independent of the phenotypic outcome controlling for batch effects,
- $H_1$: The complement of $H_0$.

The exact and approximate distributions of the statistic under the null hypothesis can be obtained through analysis or resampling (Hothorn *et al.*, 2006). However, when disease and batch strata are large, the statistic converges to $\chi_1^2$. Based on the null distribution, $p$-values are obtained for each gene.

For implementation of this test, an `R` function `coin::kruskal_wallis()` (Hothorn, 2019) is available. The `coin` package function allows only a single batch variable.

#### 1.10 two-part Kruskal-Wallis test

Nonparametric tests such as KW and Wilcoxon's rank-sum (WRS) test have minimal distributional assumptions. The lack of model-induced information often results in lack of power. While zero-inflation is a well-known characteristic of microbiome sequencing data, explicitly modeling the proportion of zeros can enhance the power in detecting differential expression. This additional assumption can be integrated into the nonparametric models using two-part model framework (Lachenbruch, 1976). Of note, the LB test and the MAST are also two-part models but they are fully parametric. Nonparametric two-part models have been used in other 'omics applications (Taylor and Pollard, 2009) and microbiome (Wagner *et al.*, 2011) data analysis. The binary part of these nonparametric models has been modeled using a conventional proportion test where no covariates are allowed. To incorporate covariate information in the binary model, a logistic regression model can be used. The KW or the WRS test can be used as the nonzero model–to allow for the inclusion of covariates in the model, the modified KW test can be used. In this paper we combine a logistic regression model and a KW test and name it two-part KW test. The binary component of the model is the same as that of LB model, i.e. Equation (3). The nonzero component's test statistic is derived based only on subjects with non-zero gene expressions and has the same formula given in KW test, i.e. Equation (13).

The $p$-values can be obtained by combining two $\chi_1^2$ statistics derived from each component:

$$T_g^{KWII} := W_g^a + W_g^b,$$

where $W_g^a$ is either a Wald test statistic $\left(\left(\frac{\hat{\beta}_g^D}{se(\hat{\beta}_g^D)}\right)^2\right)$ or a likelihood ratio statistic (two times the difference of log-likelihood of logistic regression models), and $W_g^b$ has the same form as Equation 13. The test statistic, $T_g^{KWII}$ follows a $\chi_1^2$ distribution under the null hypothesis asymptotically.

#### 2 Data generative models

The zero-inflated log-normal model is a mixture of log-normal distribution with a point mass at zero. The density is given as

$$f_{ZILN}(y) = \pi\ 1(y = 0) + (1 - \pi)\ \varphi(y, \mu, \mu\theta)1(y > 0), \tag{14}$$

where $\varphi(x, \mu, \sigma^2)$ is the log-normal density at $x$ with mean $\mu$ and variance $\sigma^2$, $\pi$ is the zero-inflation parameter, or $\pi \equiv \Pr(Y = 0)$, $\mu$ is the non-zero mean parameter (i.e. $\mu \equiv E[Y|Y > 0]$), and $\theta$ is the over-dispersion parameter so that $var[Y|Y > 0] = \mu^2\theta$

ZINB is an extension of the negative binomial distribution and is widely used to model count data with excess zeros. In many real world applications, if more zeros are observed than the negative binomial distribution assumes, the zero-inflated negative binomial distribution is suitable and has in fact become one of the most commonly used methods in count data analysis (Preisser *et al.*, 2012). This is true for omics data analysis including scRNAseq (Risso *et al.*, 2018; Van den Berge *et al.*, 2018) and microbiome data (Chen *et al.*, 2017; Xia *et al.*, 2018).

ZINB without covariates has three parameters, $(\mu, \theta, \pi)^\top$, with the following density:

$$f_{ZINB}(y) = \pi\ 1(y = 0) + (1 - \pi) \binom{y + \frac{1}{\theta} - 1}{y} \frac{(\mu\theta)^y}{(1 + \mu\theta)^{y+1/\theta}}, \tag{15}$$

$y = 0, 1, 2, ...$, where $\pi$ is the zero-inflation parameter, $\mu$ is the mean parameter assuming no zero-inflation (i.e. $E[Y] = \mu(1 - \pi)$), and $\theta$ is the over-dispersion parameter such that $var[Y] = \mu^2\pi(1 - \pi) + (1 - \pi)(\mu + \mu^2\theta)$.

We use the same notation $\xi \equiv (\mu, \theta, \pi)$ for each generative model as long as there is no ambiguity, and use a superscript denoting the model if distinction is needed.

The zero-inflated Gamma model is a mixture of a Gamma distribution and a point mass at zero. The density is given as

$$f_{ZIG}(y) = \pi\ 1(y = 0) + (1 - \pi)\ \frac{y^{\mu/\theta - 1} e^{-y/\theta}}{\Gamma(\mu/\theta)\theta^{\mu/\theta}} 1(y > 0), \tag{16}$$

where $\pi$ is the zero-inflation parameter or $\pi \equiv \Pr(Y = 0)$, $\mu$ is the non-zero mean parameter (i.e. $\mu \equiv E[Y|Y > 0]$), and $\theta$ is the over-dispersion parameter so that $var[Y|Y > 0] = \mu^2\theta$.

#### 3 Model-based simulation setup

##### 3.1 ZILN-simulation baseline parameters

##### 3.2 The baseline distribution parameters selected for ZILN- and ZIG-based simulations.

##### 3.3 ZOE 2.0 data parameter estimates for the ZINB models

The baseline parameters, disease effects, and batch effects for ZINB models are chosen (Web Table 3) based on the parameter estimates from the ZOE 2.0 data (Web Figure 1).

##### 3.4 Estimated parameters from the validation data

The estimated parameters of the ZILN (and ZIG) models from the validation data—the ZOE 2.0 pilot and the IBD data—are presented.

###### 3.4.1 Estimated parameters from the ZOE-pilot data.

Web Figure 2 and Web Table 5 illustrate the distribution of the ZILN parameter estimates in the ZOE-pilot data.

###### 3.4.2 Estimated parameters from the IBD data

Web Figure 3 and Web Table 6 illustrate the distribution of the ZILN parameter estimates in the IBD data.

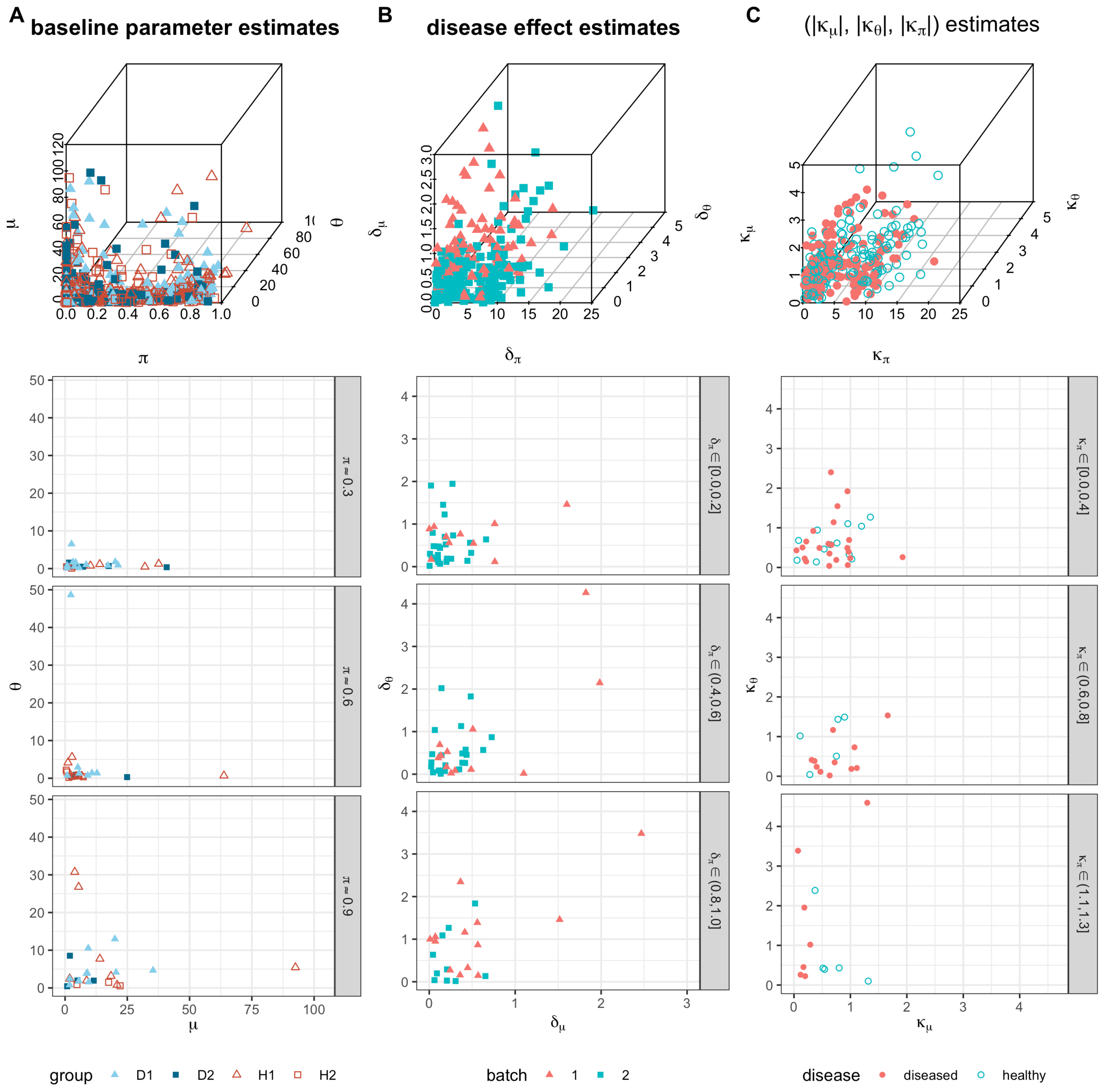


Web Figure 1: ZINB-parameter estimates from the ZOE 2.0 data. (A) baseline parameter estimates (TOP: $\pi > 0.8$, MIDDLE: $\pi \in (0.4, 0.8]$, BOTTOM: $\pi \leq 0.4$), (B) disease effect parameter estimates, (C) batch effect parameter estimates. The quartiles of the estimated parameters are provided in Web Table 2.

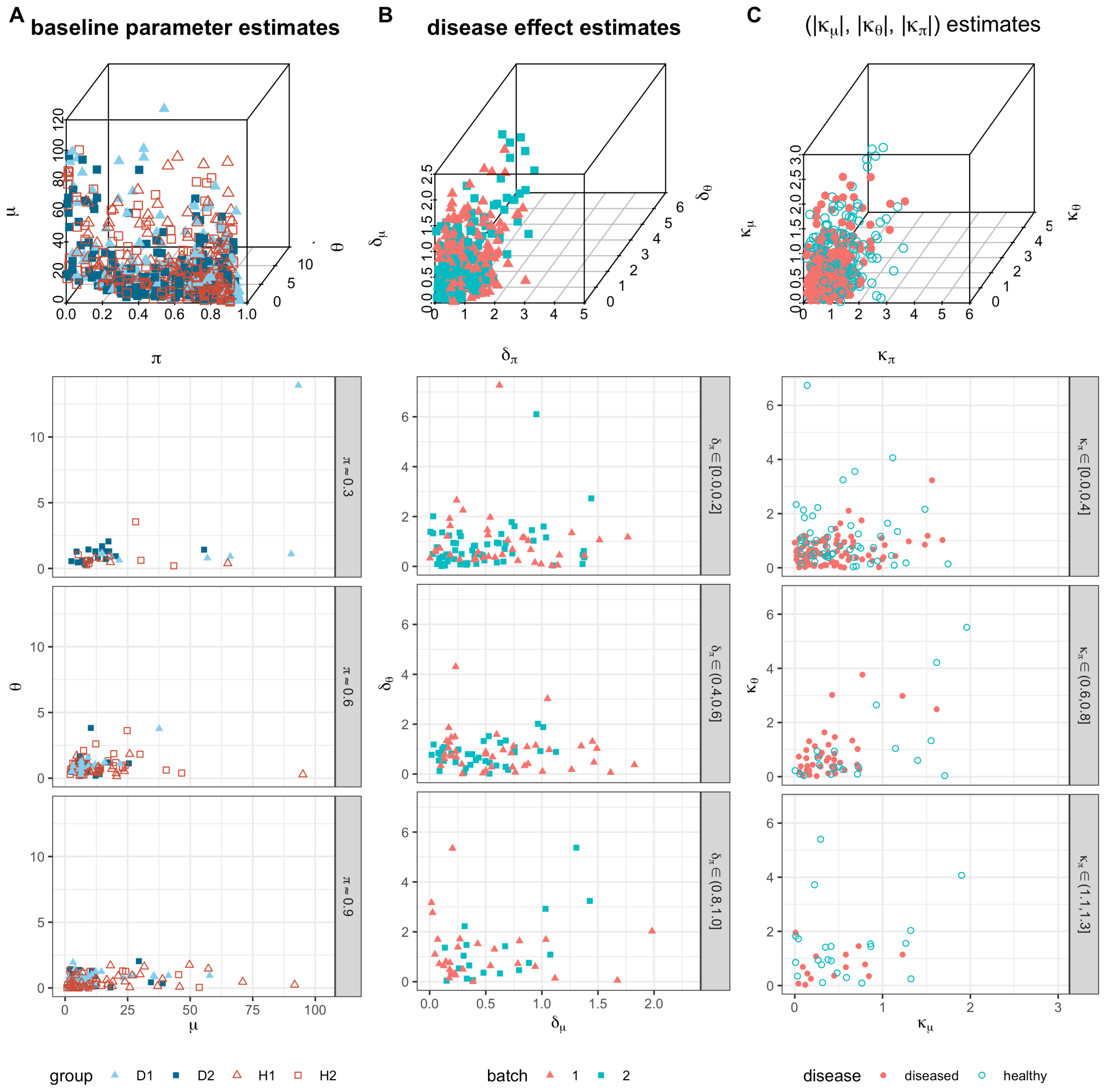


Web Figure 2: The ZILN parameter estimates for genes in the ZOE-pilot data
Column A: parameter estimates of baseline ZILN distributions from the ZOE-pilot data with the 3-dimensional scatter plot on the top row and each of the subsequent rows representing $\pi$ estimates being within 0.03 from 0.9, 0.6, and 0.3.
Column B: disease effect estimates based on ZILN models from the ZOE-pilot data in absolute values $(|\delta_\mu|, |\delta_\theta|, |\delta_\pi|)$
Column C: batch effect estimates based on ZILN models from the ZOE-pilot data in absolute values $(|\kappa_\mu|, |\kappa_\theta|, |\kappa_\pi|)$. The quartiles of the estimated parameters are provided in Web Table 5.

| No. | $\mu$ | $\theta$ | $\pi$ |
|---|---|---|---|
| B01-03 | 1, 10, 50 | 0.5 | 0.3 |
| B04-06 | 1, 10, 50 | 2.0 | 0.3 |
| B07-09 | 1, 10, 50 | 0.5 | 0.6 |
| B10-12 | 1, 10, 50 | 2.0 | 0.6 |
| B13-15 | 1, 10, 50 | 0.5 | 0.65 |
| B16-18 | 1, 10, 50 | 2.0 | 0.65 |
| B19-21 | 1, 10, 50 | 0.5 | 0.7 |
| B22-24 | 1, 10, 50 | 2.0 | 0.7 |
| B25-27 | 1, 10, 50 | 0.5 | 0.75 |
| B28-30 | 1, 10, 50 | 2.0 | 0.75 |
| B31-33 | 1, 10, 50 | 0.5 | 0.8 |
| B34-36 | 1, 10, 50 | 2.0 | 0.8 |
| B37-39 | 1, 10, 50 | 0.5 | 0.85 |
| B40-42 | 1, 10, 50 | 2.0 | 0.85 |
| B43-45 | 1, 10, 50 | 0.5 | 0.9 |
| B46-48 | 1, 10, 50 | 2.0 | 0.9 |
| B49-51 | 1, 10, 50 | 0.5 | 0.95 |
| B52-54 | 1, 10, 50 | 2.0 | 0.95 |

Web Table 1: Baseline ZILN parameters

| parameters | 1Q | 2Q | 3Q |
|---|---|---|---|
| $\mu$ | 0.80 | 2.60 | 8.00 |
| $\theta$ | 0.40 | 0.80 | 2.90 |
| $\pi$ | 0.00 | 0.10 | 0.63 |
| $\delta_\mu$ | 0.10 | 0.30 | 0.60 |
| $\delta_\theta$ | 0.20 | 0.50 | 1.10 |
| $\delta_\pi$ | 0.50 | 1.30 | 6.60 |
| $\kappa_\mu$ | 0.40 | 0.90 | 1.40 |
| $\kappa_\theta$ | 0.30 | 0.60 | 1.30 |
| $\kappa_\pi$ | 1.20 | 2.80 | 8.90 |

Web Table 2: The ZINB parameter estimate distribution of genes in the ZOE 2.0 data

#### 3.5 Estimated parameters for other aspects of the data

##### 3.5.1 Parameters for the gene expression of each gene-species combination

Web Figure 4 and Web Table 7 illustrate the distribution of the ZILN parameter estimates of the gene-species combinations in the ZOE 2.0 data.

The estimated parameters (Web Figure 4) of the gene expression distribution for gene-species combinations are well covered by the parameter sets in Table 1.

Web Figure 5 and Web Table 8 illustrate the distribution of the ZINB parameter estimates of gene-species combinations in the ZOE 2.0 data.

The estimated parameters (Web Figure 5) of the gene expression distribution for gene-species combinations are well covered by the parameter sets in Web Table 3.

##### 3.5.2 Estimated parameters for the total gene expression of each species

Web Figure 6 and Web Table 9 illustrate the distribution of the ZILN parameter estimates of species in the ZOE 2.0 data.

Web Figure 7 and Web Table 10 illustrate the distribution of the ZINB parameter estimates of species in the ZOE 2.0 data.

Estimated parameters (Web Figure 6) of gene expression distribution for species are well covered by the parameter sets in Table 1.

Estimated parameters (Web Figure 7) of gene expression distribution for species can be mostly covered by the parameter sets in Web Table 3 except when $\pi$ is large and the overdispersion parameter $\theta$ is very irregular.

| No. | $\mu$ | $\theta$ | $\pi$ |
|---|---|---|---|
| B01-03 | 1, 10, 50 | 1 | 0.3 |
| B04-06 | 1, 10, 50 | 5 | 0.3 |
| B07-09 | 1, 10, 50 | 1 | 0.6 |
| B10-12 | 1, 10, 50 | 5 | 0.6 |
| B13-15 | 1, 10, 50 | 1 | 0.65 |
| B16-18 | 1, 10, 50 | 5 | 0.65 |
| B19-21 | 1, 10, 50 | 1 | 0.7 |
| B22-24 | 1, 10, 50 | 5 | 0.7 |
| B25-27 | 1, 10, 50 | 1 | 0.75 |
| B28-30 | 1, 10, 50 | 5 | 0.75 |
| B31-33 | 1, 10, 50 | 1 | 0.8 |
| B34-36 | 1, 10, 50 | 5 | 0.8 |
| B37-39 | 1, 10, 50 | 1 | 0.85 |
| B40-42 | 1, 10, 50 | 5 | 0.85 |
| B43-45 | 1, 10, 50 | 1 | 0.9 |
| B46-48 | 1, 10, 50 | 5 | 0.9 |
| B49-51 | 1, 10, 50 | 1 | 0.95 |
| B52-54 | 1, 10, 50 | 5 | 0.95 |

Web Table 3: Baseline ZINB-parameters

| No. | $\mu$ | $\theta$ | $\pi$ |
|---|---|---|---|
| B01-03 | 1, 10, 50 | 0.5 | 0.3 |
| B04-06 | 1, 10, 50 | 2.0 | 0.3 |
| B07-09 | 1, 10, 50 | 0.5 | 0.6 |
| B10-12 | 1, 10, 50 | 2.0 | 0.6 |
| B13-15 | 1, 10, 50 | 0.5 | 0.65 |
| B16-18 | 1, 10, 50 | 2.0 | 0.65 |
| B19-21 | 1, 10, 50 | 0.5 | 0.7 |
| B22-24 | 1, 10, 50 | 2.0 | 0.7 |
| B25-27 | 1, 10, 50 | 0.5 | 0.75 |
| B28-30 | 1, 10, 50 | 2.0 | 0.75 |
| B31-33 | 1, 10, 50 | 0.5 | 0.8 |
| B34-36 | 1, 10, 50 | 2.0 | 0.8 |
| B37-39 | 1, 10, 50 | 0.5 | 0.85 |
| B40-42 | 1, 10, 50 | 2.0 | 0.85 |
| B43-45 | 1, 10, 50 | 0.5 | 0.9 |
| B46-48 | 1, 10, 50 | 2.0 | 0.9 |
| B49-51 | 1, 10, 50 | 0.5 | 0.95 |
| B52-54 | 1, 10, 50 | 2.0 | 0.95 |

Web Table 4: Baseline ZIG-parameters

| parameters | 1Q | 2Q | 3Q |
|---|---|---|---|
| $\mu$ 5.30 | 10.80 | 25.90 | |
| $\theta$ 0.40 | 0.70 | 1.10 | |
| $\pi$ 0.48 | 0.72 | 0.86 | |
| $\delta_\mu$ 0.20 | 0.40 | 0.70 | |
| $\delta_\theta$0.30 | 0.70 | 1.30 | |
| $\delta_\pi$ 0.20 | 0.50 | 0.90 | |
| $\kappa_\mu$ 0.20 | 0.40 | 0.70 | |
| $\kappa_\theta$0.20 | 0.60 | 1.10 | |
| $\kappa_\pi$ 0.30 | 0.70 | 1.20 | |

Web Table 5: The ZILN parameter estimate distribution of genes in the ZOE-pilot data

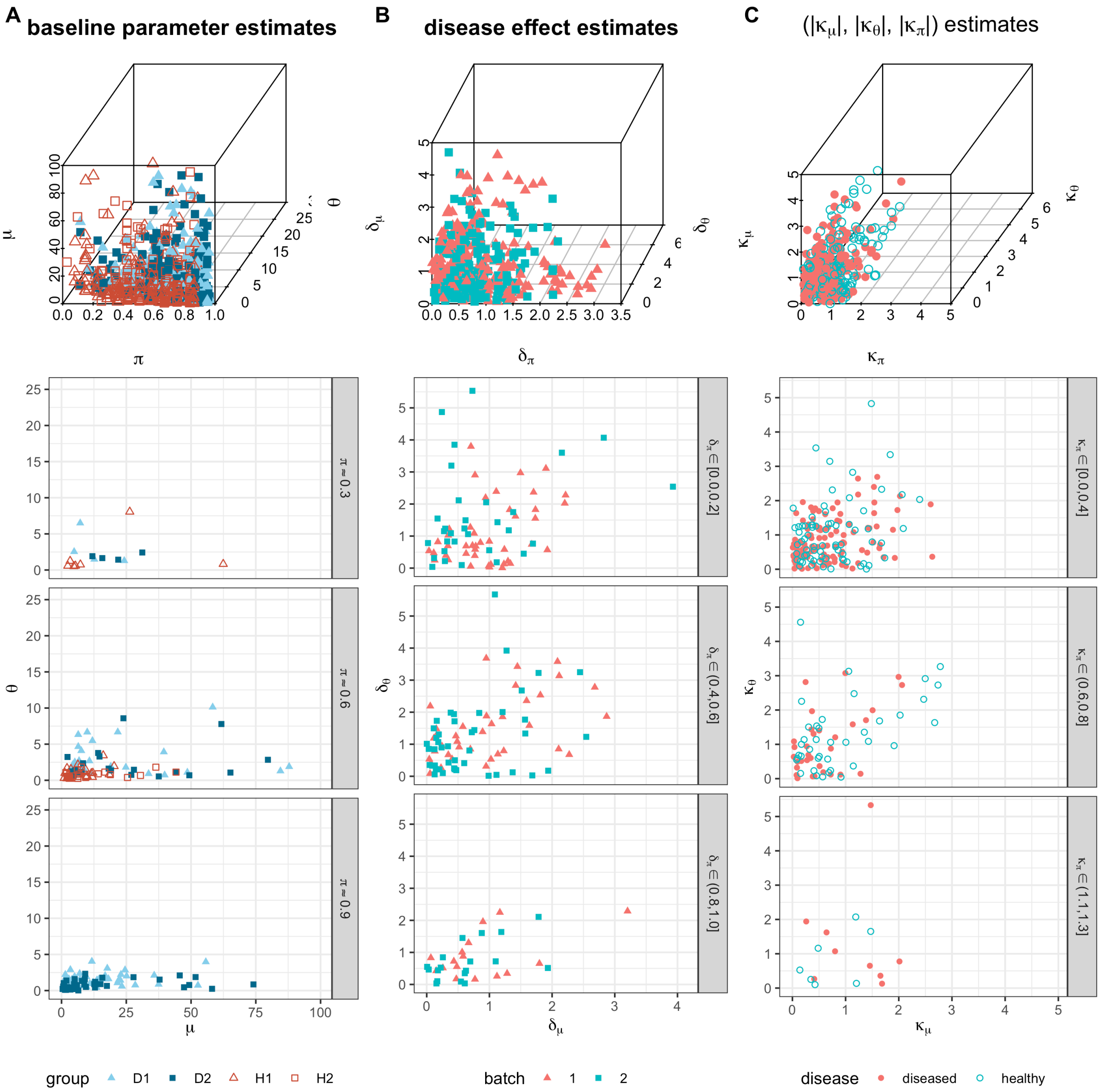


Web Figure 3: The ZILN parameter estimates for genes in the IBD data.
Column A: parameter estimates of baseline ZILN distributions from the IBD data with the 3-dimensional scatter plot on the top row and each of the subsequent rows representing $\pi$ estimates being within 0.03 from 0.9, 0.6, and 0.3.
Column B: disease effect estimates based on ZILN models from the IBD data in absolute values $(|\delta_\mu|, |\delta_\theta|, |\delta_\pi|)$
Column C: batch effect estimates based on ZILN models from the IBD data in absolute values $(|\kappa_\mu|, |\kappa_\theta|, |\kappa_\pi|)$. The quartiles of the estimated parameters are provided in Web Table 6.

| parameters | 1Q | 2Q | 3Q |
|---|---|---|---|
| $\mu$ | 0.00 | 0.10 | 0.20 |
| $\theta$ | 0.50 | 1.00 | 1.80 |
| $\pi$ | 0.56 | 0.75 | 0.86 |
| $\delta_\mu$ | 0.30 | 0.70 | 1.30 |
| $\delta_\theta$ | 0.40 | 0.90 | 1.60 |
| $\delta_\pi$ | 0.30 | 0.60 | 1.00 |
| $\kappa_\mu$ | 0.30 | 0.60 | 1.00 |
| $\kappa_\theta$ | 0.30 | 0.80 | 1.40 |
| $\kappa_\pi$ | 0.30 | 0.50 | 0.80 |

Web Table 6: The ZILN parameter estimate distribution of genes in the IBD data

| parameters | 1Q | 2Q | 3Q |
|---|---|---|---|
| $\mu$ | 3.40 | 7.70 | 21.50 |
| $\theta$ | 0.70 | 1.10 | 1.70 |
| $\pi$ | 0.39 | 0.67 | 0.83 |
| $\delta_\mu$ | 0.10 | 0.20 | 0.50 |
| $\delta_\theta$ | 0.20 | 0.50 | 0.90 |
| $\delta_\pi$ | 0.20 | 0.30 | 0.50 |
| $\kappa_\mu$ | 0.30 | 0.60 | 1.00 |
| $\kappa_\theta$ | 0.30 | 0.70 | 1.30 |
| $\kappa_\pi$ | 0.40 | 0.70 | 1.30 |

Web Table 7: The ZILN parameter estimate distribution of gene-species combinations in the ZOE 2.0 data

| parameters | 1Q | 2Q | 3Q |
|---|---|---|---|
| $\mu$ | 2.00 | 5.90 | 16.70 |
| $\theta$ | 0.40 | 0.90 | 2.10 |
| $\pi$ | 0.00 | 0.37 | 0.71 |
| $\delta_\mu$ | 0.10 | 0.30 | 0.60 |
| $\delta_\theta$ | 0.20 | 0.50 | 1.20 |
| $\delta_\pi$ | 0.30 | 0.80 | 3.30 |
| $\kappa_\mu$ | 0.40 | 0.80 | 1.30 |
| $\kappa_\theta$ | 0.30 | 0.70 | 1.40 |
| $\kappa_\pi$ | 0.70 | 1.60 | 6.70 |

Web Table 8: The ZINB parameter estimate distribution of gene-species combinations in the ZOE 2.0 data

| parameters | 1Q | 2Q | 3Q |
|---|---|---|---|
| $\mu$ | 5.80 | 13.10 | 39.40 |
| $\theta$ | 0.50 | 0.80 | 1.40 |
| $\pi$ | 0.25 | 0.53 | 0.79 |
| $\delta_\mu$ | 0.10 | 0.20 | 0.40 |
| $\delta_\theta$ | 0.20 | 0.50 | 0.90 |
| $\delta_\pi$ | 0.20 | 0.30 | 0.50 |
| $\kappa_\mu$ | 0.30 | 0.50 | 1.00 |
| $\kappa_\theta$ | 0.30 | 0.60 | 1.00 |
| $\kappa_\pi$ | 0.30 | 0.70 | 1.20 |

Web Table 9: The ZILN parameter estimate distribution of species in the ZOE 2.0 data

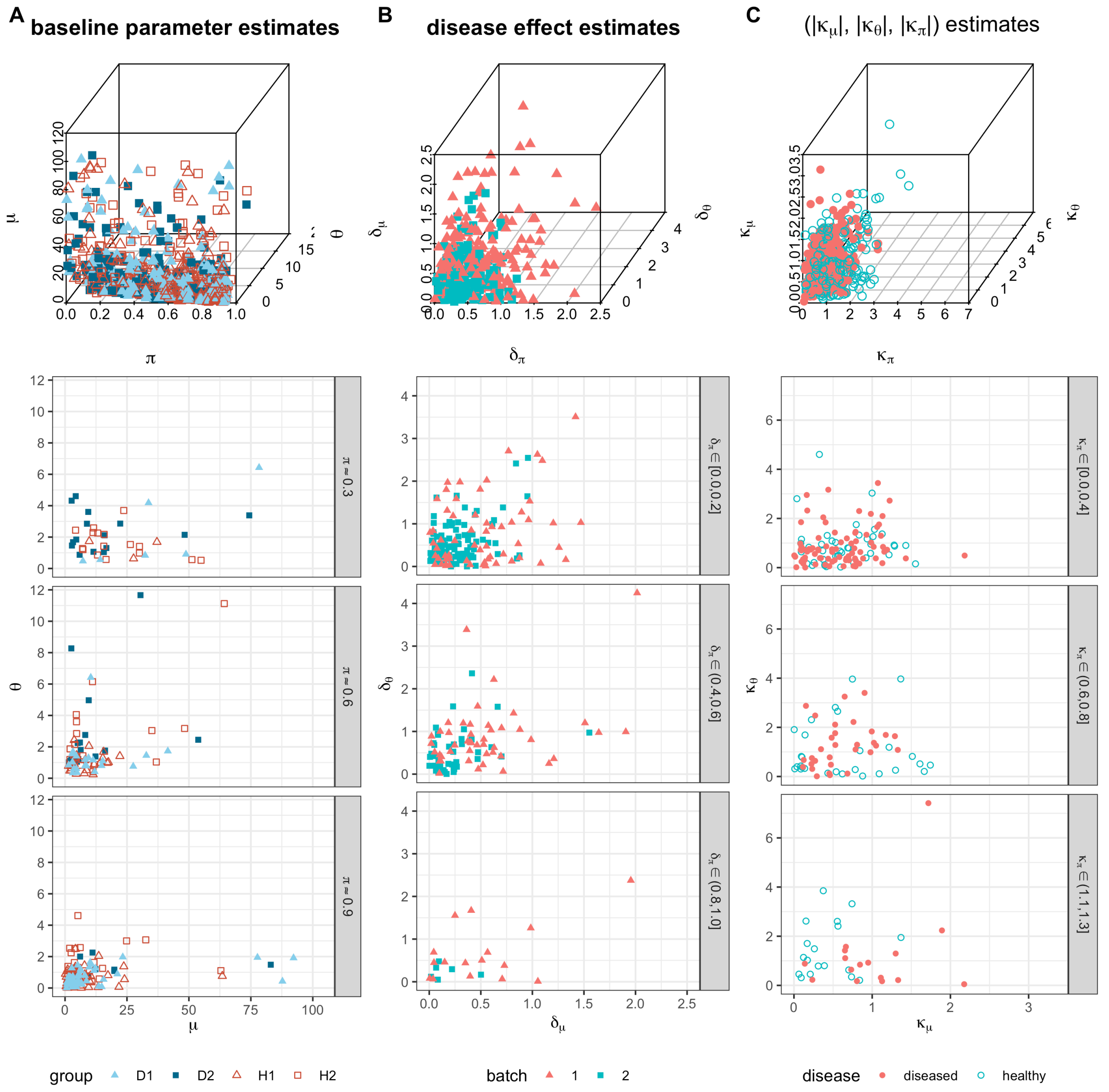


Web Figure 4: The ZILN parameter estimates for gene-species combination
Column A: parameter estimates of baseline ZILN distributions from the ZOE 2.0 data with the 3-dimensional scatter plot on the top row and each of the subsequent rows representing $\pi$ estimates being within 0.03 from 0.9, 0.6, and 0.3.
Column B: disease effect estimates based on ZILN models from the ZOE 2.0 data in absolute values $(|\delta_\mu|, |\delta_\theta|, |\delta_\pi|)$
Column C: batch effect estimates based on ZILN models from the ZOE 2.0 data in absolute values $(|\kappa_\mu|, |\kappa_\theta|, |\kappa_\pi|)$. The quartiles of the estimated parameters are provided in Web Table 7.

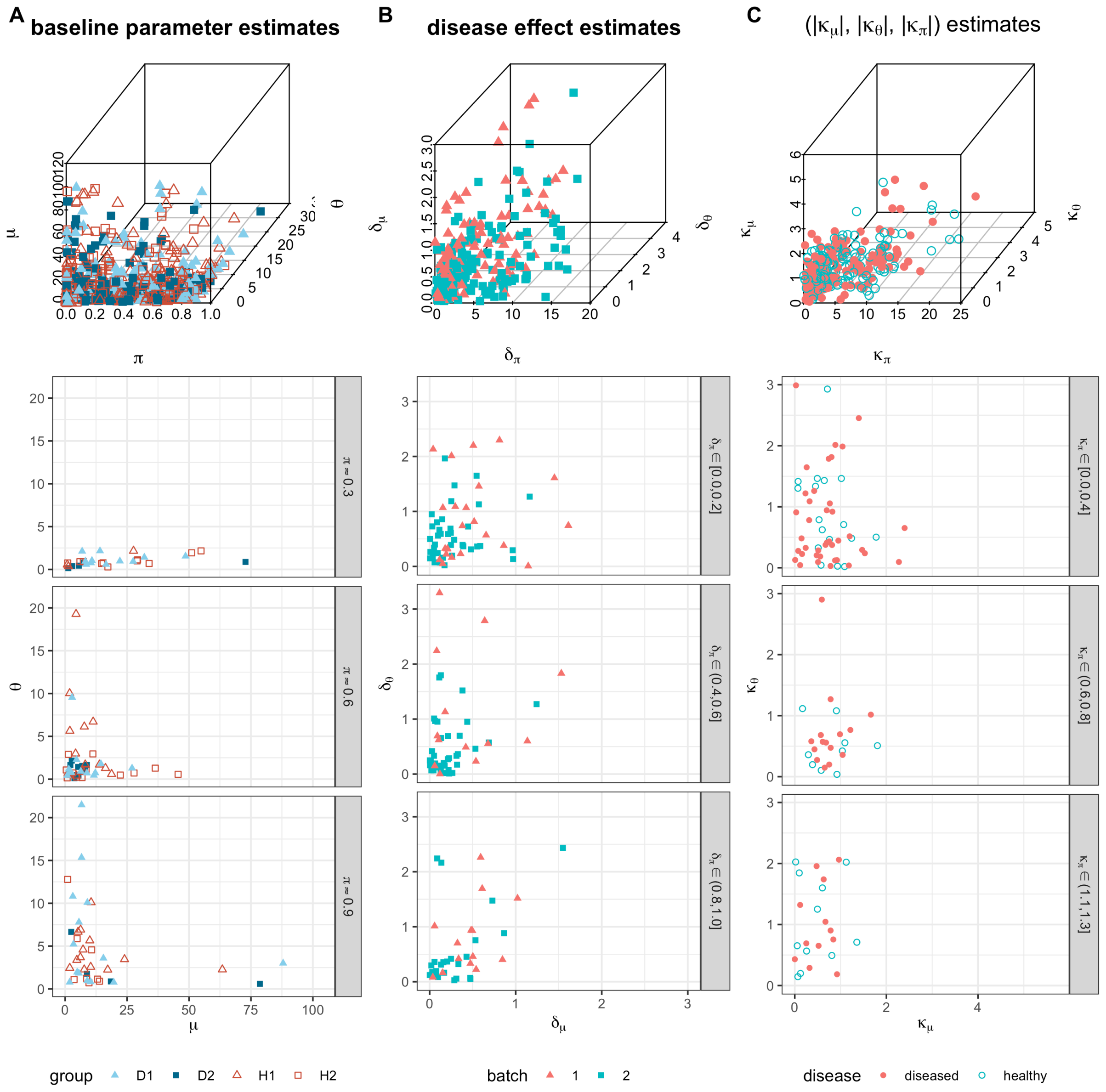


Web Figure 5: ZINB parameter estimates for gene-species combinations
Column A: parameter estimates of baseline ZINB distributions in the ZOE 2.0 data presented with a 3-dimensional scatter plot on the top row and each of the subsequent rows represents $\pi$ estimates being within 0.03 from 0.9, 0.6, and 0.3.
Column B: disease effect estimates based on ZINB models from the ZOE 2.0 data in absolute values $(|\delta_\mu|, |\delta_\theta|, |\delta_\pi|)$
Column C: batch effect estimates based on ZINB models from the ZOE 2.0 data in absolute values $(|\kappa_\mu|, |\kappa_\theta|, |\kappa_\pi|)$. The quartiles of the estimated parameters are provided in Web Table 8.

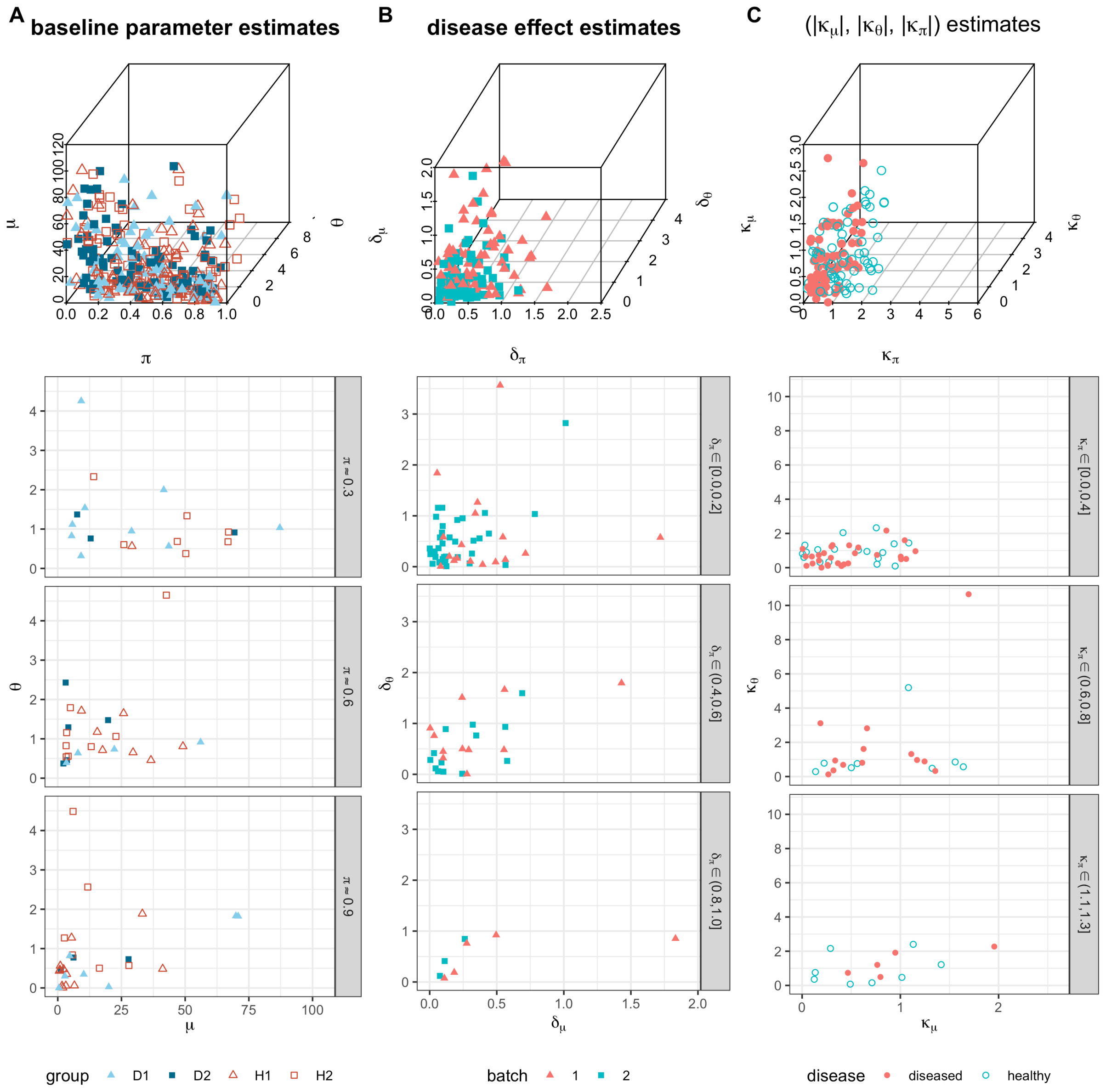


Web Figure 6: ZILN parameter estimates for species

Column A: parameter estimates of baseline ZILN distributions in the ZOE 2.0 data are presented with a 3-dimensional scatter plot on the top row and each of the subsequent rows represents $\pi$ estimates being within 0.03 from 0.9, 0.6, and 0.3.

Column B: disease effect estimates based on ZILN models from the ZOE 2.0 data in absolute values $(|\delta_\mu|, |\delta_\theta|, |\delta_\pi|)$

Column C: batch effect estimates based on ZILN models from the ZOE 2.0 data in absolute values $(|\kappa_\mu|, |\kappa_\theta|, |\kappa_\pi|)$. The quartiles of the estimated parameters are provided in Web Table 9.

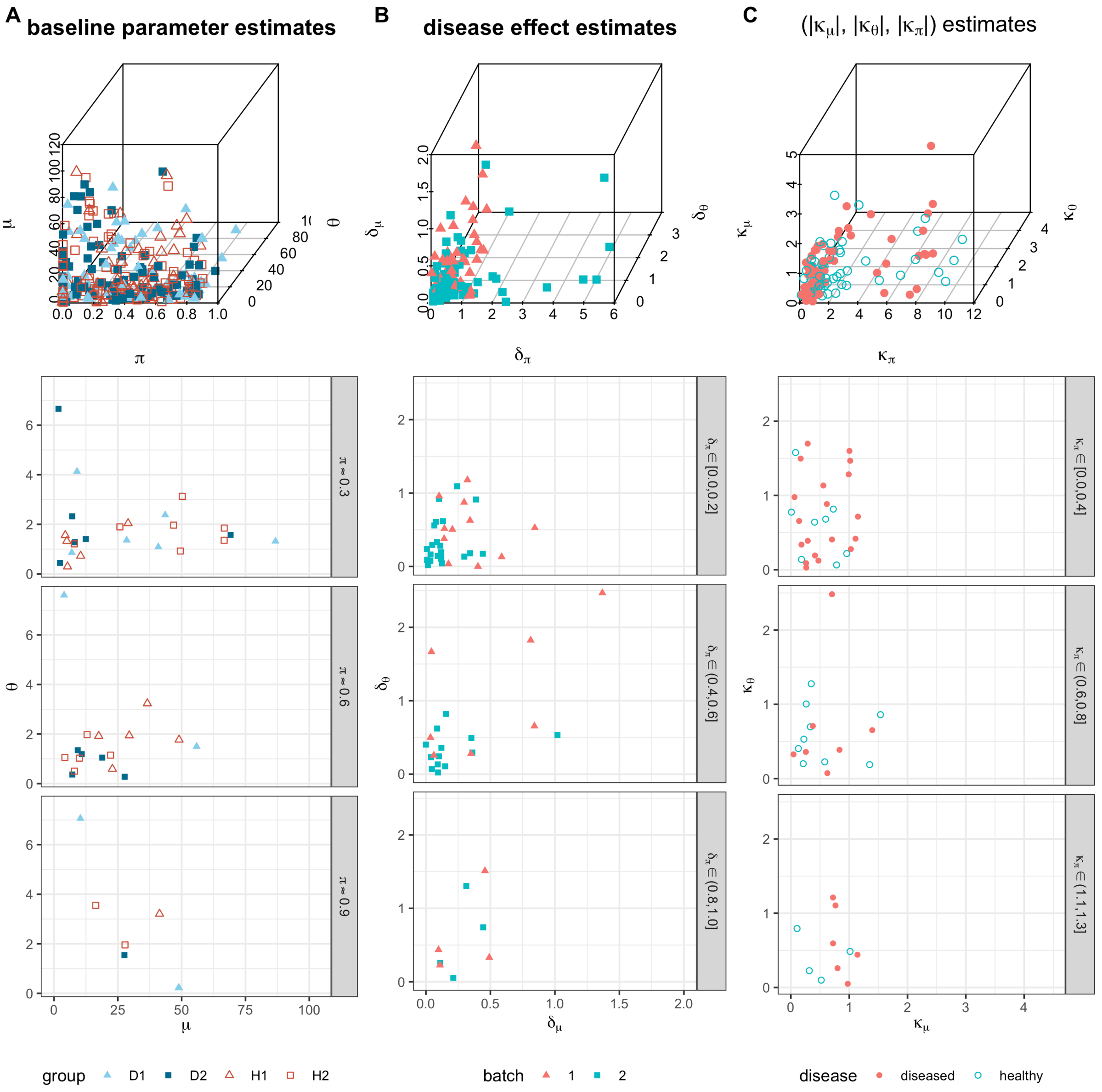


Web Figure 7: ZINB parameter estimates for species
Column A: parameter estimates of baseline ZINB distributions in the ZOE 2.0 data are presented with a 3-dimensional scatter plot on the top row and each of the subsequent rows represents $\pi$ estimates being within 0.03 from 0.9, 0.6, and 0.3.
Column B: disease effect estimates based on ZINB models from the ZOE 2.0 data in absolute values $(|\delta_\mu|, |\delta_\theta|, |\delta_\pi|)$
Column C: batch effect estimates based on ZINB models from the ZOE 2.0 data in absolute values $(|\kappa_\mu|, |\kappa_\theta|, |\kappa_\pi|)$. The quartiles of the estimated parameters are provided in Web Table 10.

| parameters | 1Q | 2Q | 3Q |
|---|---|---|---|
| $\mu$ | 4.40 | 11.30 | 30.60 |
| $\theta$ | 0.80 | 1.50 | 2.80 |
| $\pi$ | 0.14 | 0.38 | 0.67 |
| $\delta_\mu$ | 0.10 | 0.20 | 0.40 |
| $\delta_\theta$ | 0.20 | 0.40 | 0.80 |
| $\delta_\pi$ | 0.20 | 0.50 | 1.10 |
| $\kappa_\mu$ | 0.30 | 0.60 | 1.00 |
| $\kappa_\theta$ | 0.20 | 0.60 | 1.00 |
| $\kappa_\pi$ | 0.50 | 1.10 | 2.20 |

Web Table 10: The ZINB parameter estimate distribution of species in the ZOE 2.0 data

### 4 Goodness of fit results

#### 4.1 Goodness of fit for the ZINB models

Web Figure 8 presents the goodness of fit results of the ZINB model for two randomly chosen genes in the ZOE 2.0 data. The ZINB distribution provides a decent approximation to the RPK and TPM transformations, although it does not fit well to the arcsin transformed data.

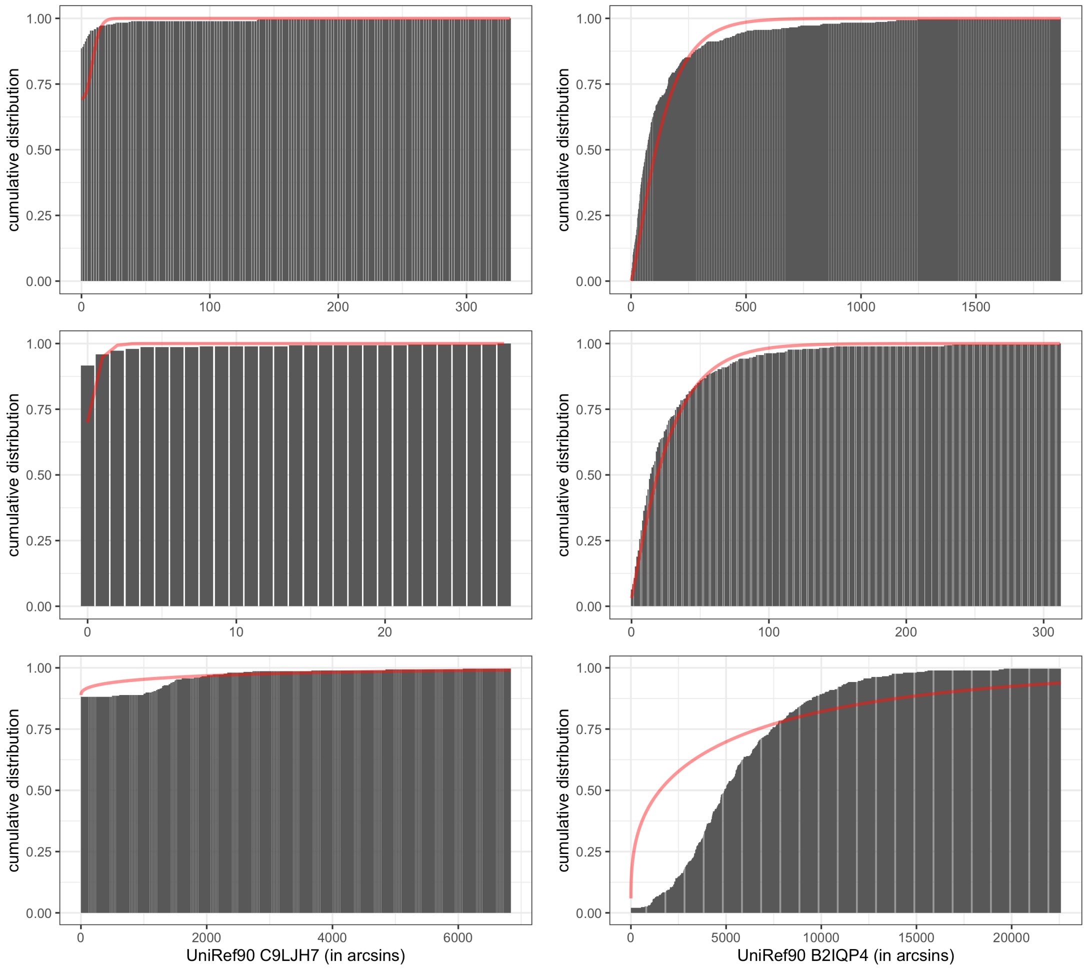


Web Figure 8: The empirical distribution (bars) and the estimated ZINB distribution (curves) based on RPK (top), TPM (middle), and arcsin (bottom) transformation of two randomly chosen genes (LEFT: C9LJH7, RIGHT: B2QP4).

### 5 Full model-based simulation results

#### 5.1 Full results under ZILN models

##### 5.1.1 Full results under ZILN models—Sensitivity

See Figures 9 and 10.

##### 5.1.2 Type I error and FDR under ZILN model under D2 scenario

See Figures 11.

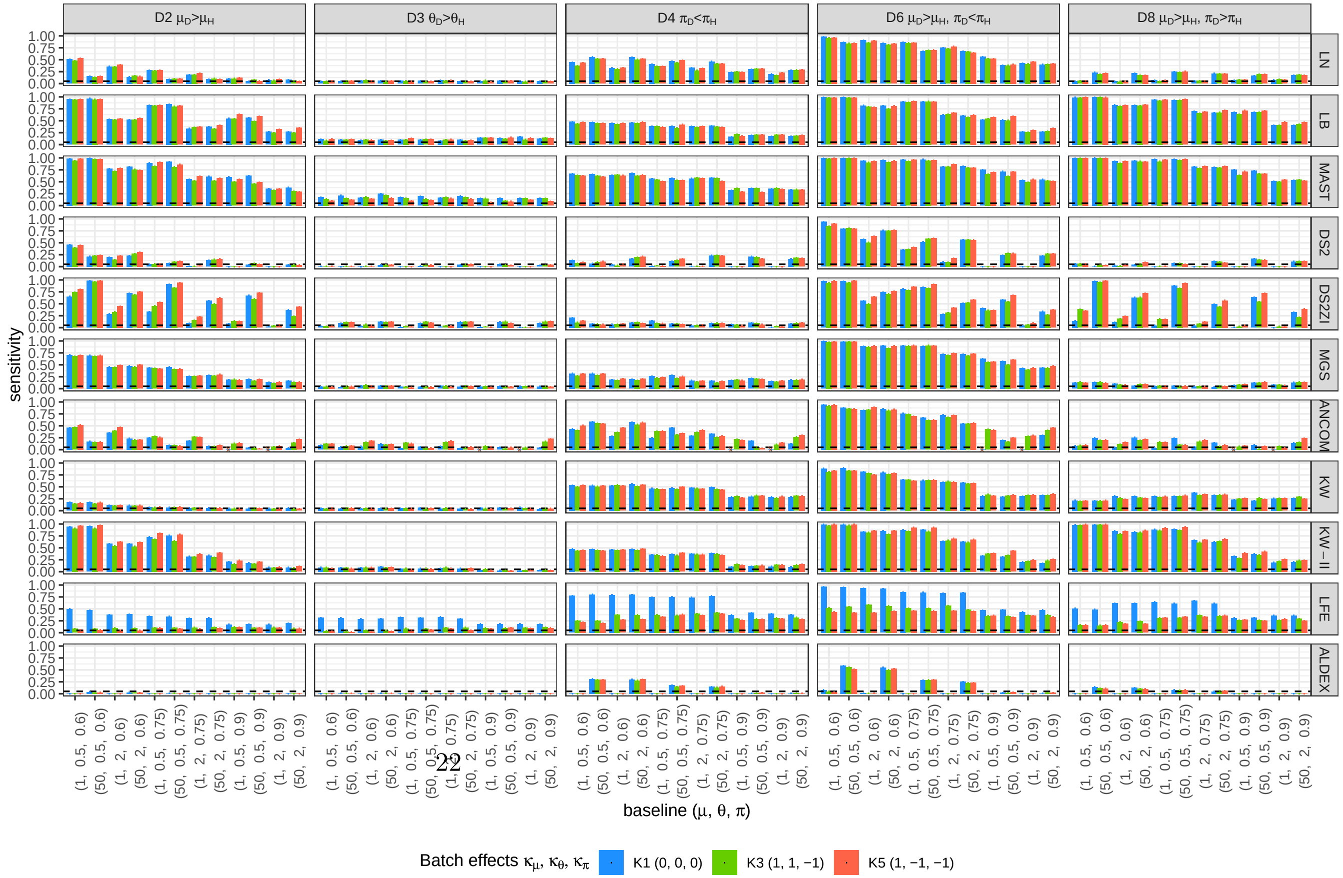




Web Figure 9: Full results under ZILN model for a sample size of 80. DS2 = DESeq2, DS2ZI = DESeq2-ZINBWaVE, ANCOM.sz = ANCOM-BC1, ANCOM = ANCOM-BC2, LFE = LEfSe, ALDEX = ALDEx2.

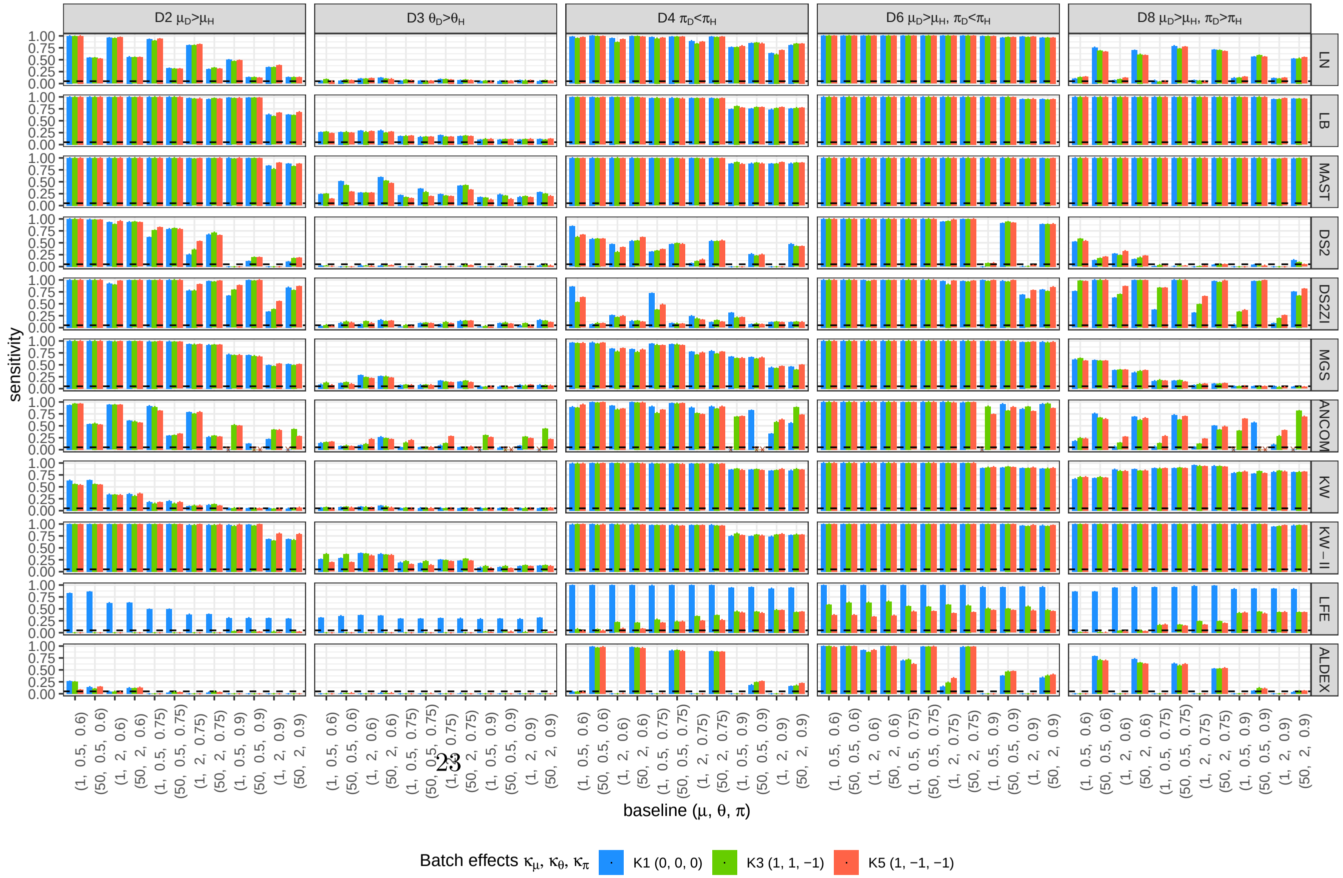




Web Figure 10: Full results under ZILN model for a sample size of 400. DS2 = DESeq2, DS2ZI = DESeq2-ZINBWaVE, ANCOM.sz = ANCOM-BC1, ANCOM = ANCOM-BC2, LFE = LEfSe, ALDEX = ALDEx2.

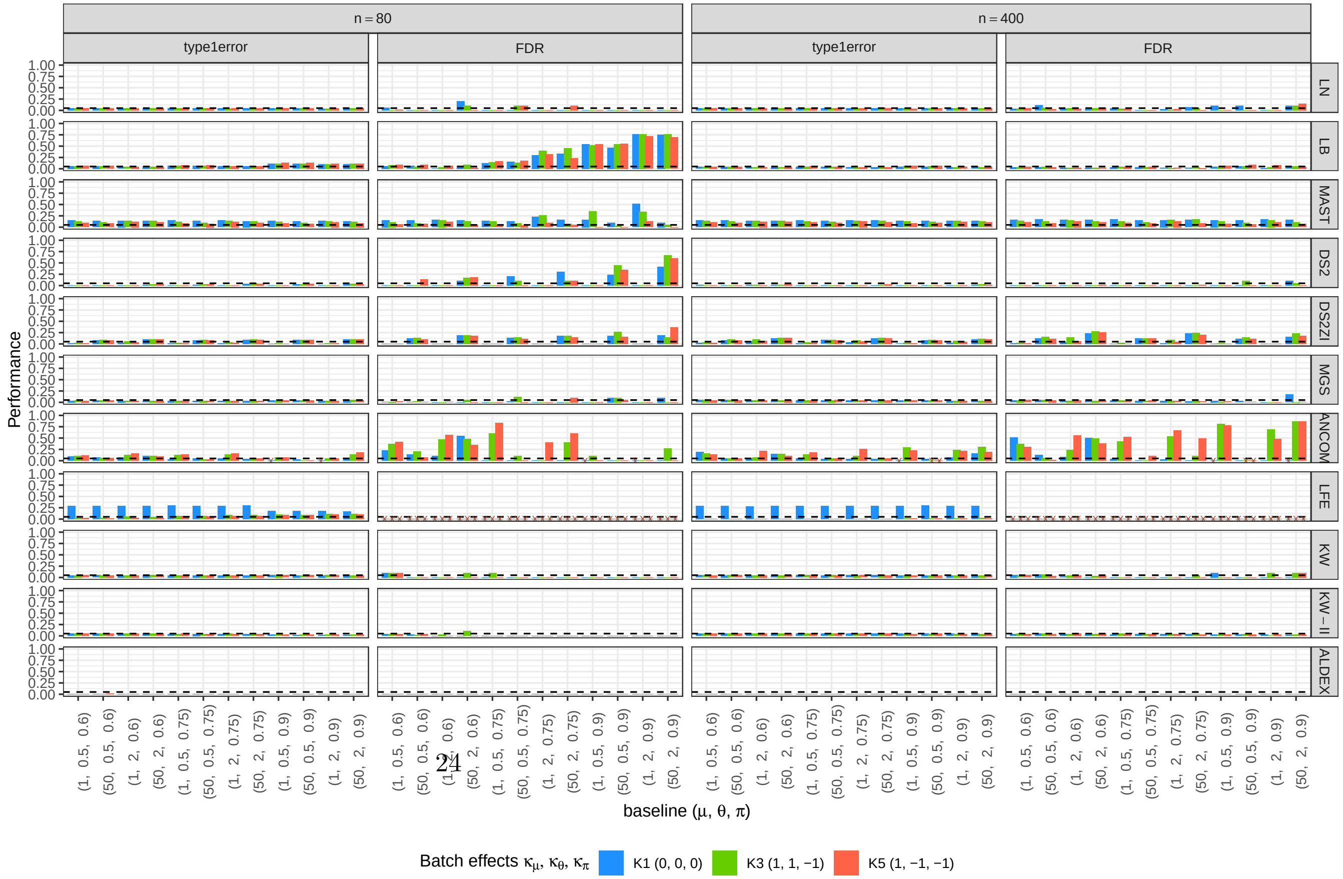




Web Figure 11: Type I error and FDR results under ZILN model under D2 scenario. DS2 = DESeq2, DS2ZI = DESeq2-ZINBWaVE, ANCOM.sz = ANCOM-BC1, ANCOM = ANCOM-BC2, LFE = LEfSe, ALDEX = ALDEx2.

#### 5.2 Full results under ZINB models

#### 5.3 Full results under ZINB models—Sensitivity

See Figures 12 and 13.

##### 5.3.1 Type I error and FDR under ZINB model under D2 scenario

See Figures 14.

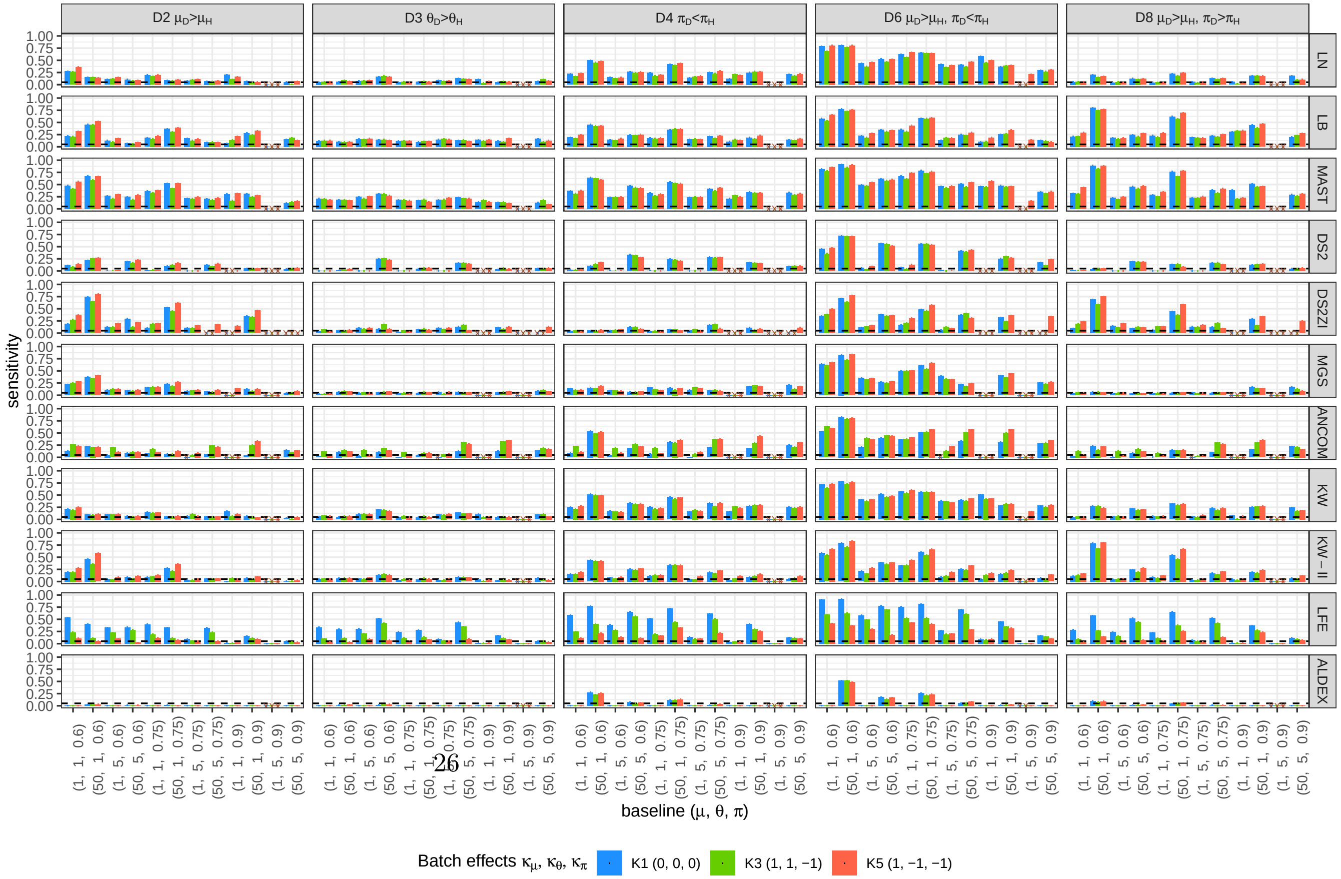


Web Figure 12: Full results under ZINB model for a sample size of 80. DS2 = DESeq2, DS2ZI = DESeq2-ZINBWaVE, ANCOM.sz = ANCOM-BC1, ANCOM = ANCOM-BC2, LFE = LEfSe, ALDEX = ALDEx2.

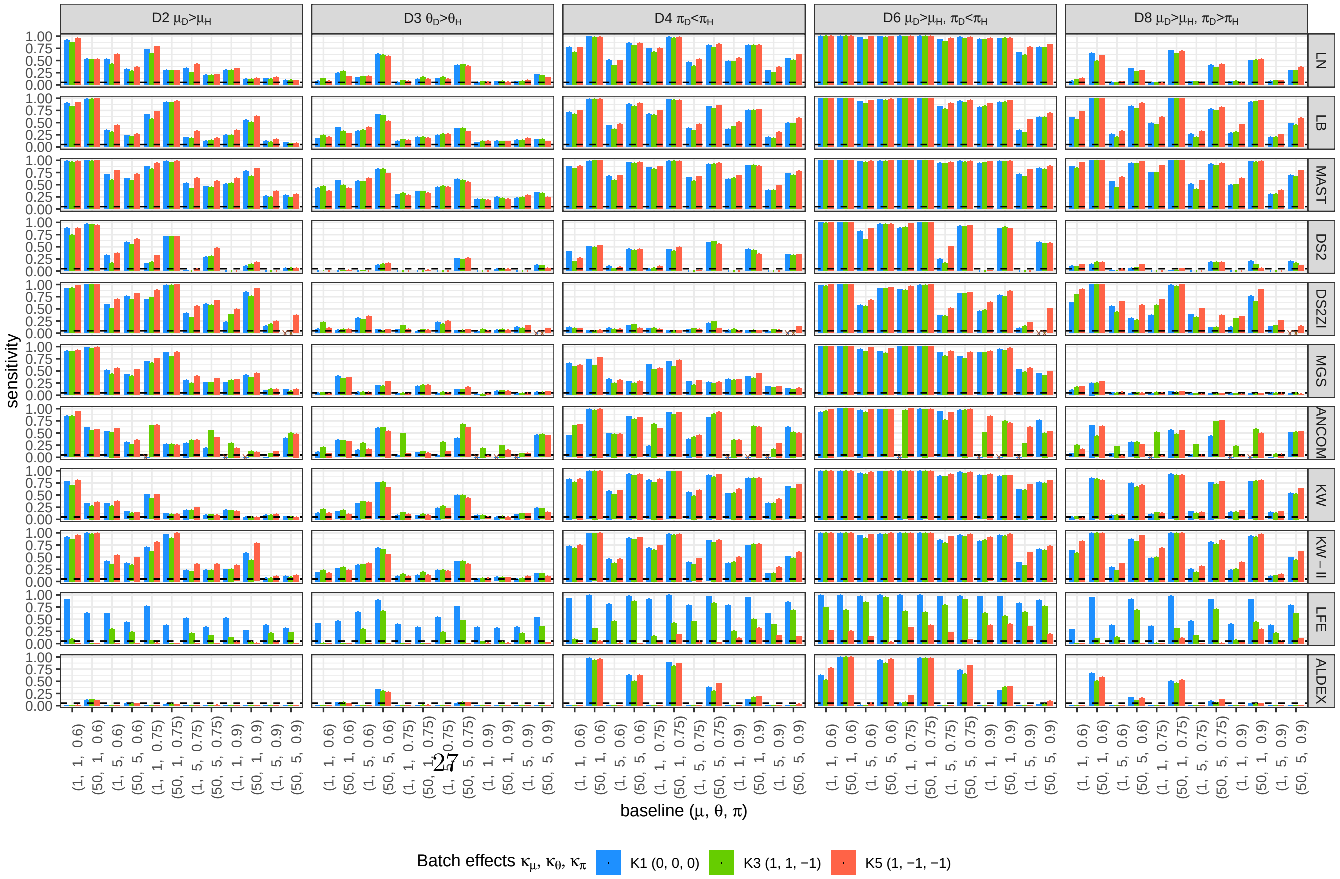

Web Figure 13: Full results under ZINB model for a sample size of 400. DS2 = DESeq2, DS2ZI = DESeq2-ZINBWaVE, ANCOM.sz = ANCOM-BC1, ANCOM = ANCOM-BC2, LFE = LEfSe, ALDEX = ALDEx2.

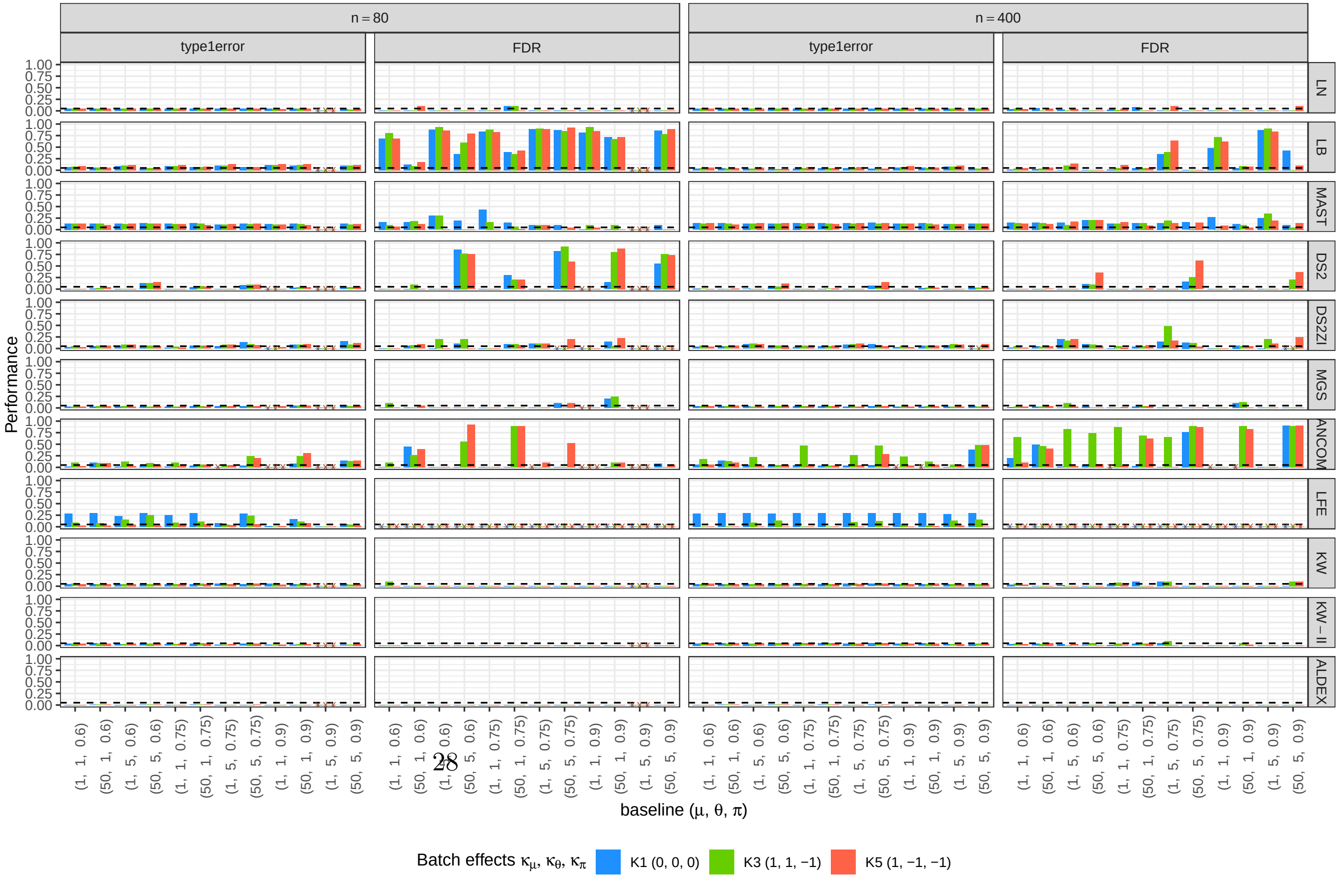




Web Figure 14: Type I error and FDR results under ZINB model under D2 scenario. DS2 = DESeq2, DS2ZI = DESeq2-ZINBWaVE, ANCOM.sz = ANCOM-BC1, ANCOM = ANCOM-BC2, LFE = LEfSe, ALDEX = ALDEx2.

##### 5.4 Full results under ZIG models

##### 5.5 Full results under ZIG models—Sensitivity

See Figures 15 and 16.

##### 5.6 Type I error and FDR results under ZIG model under D2 scenario

See Figure 17.

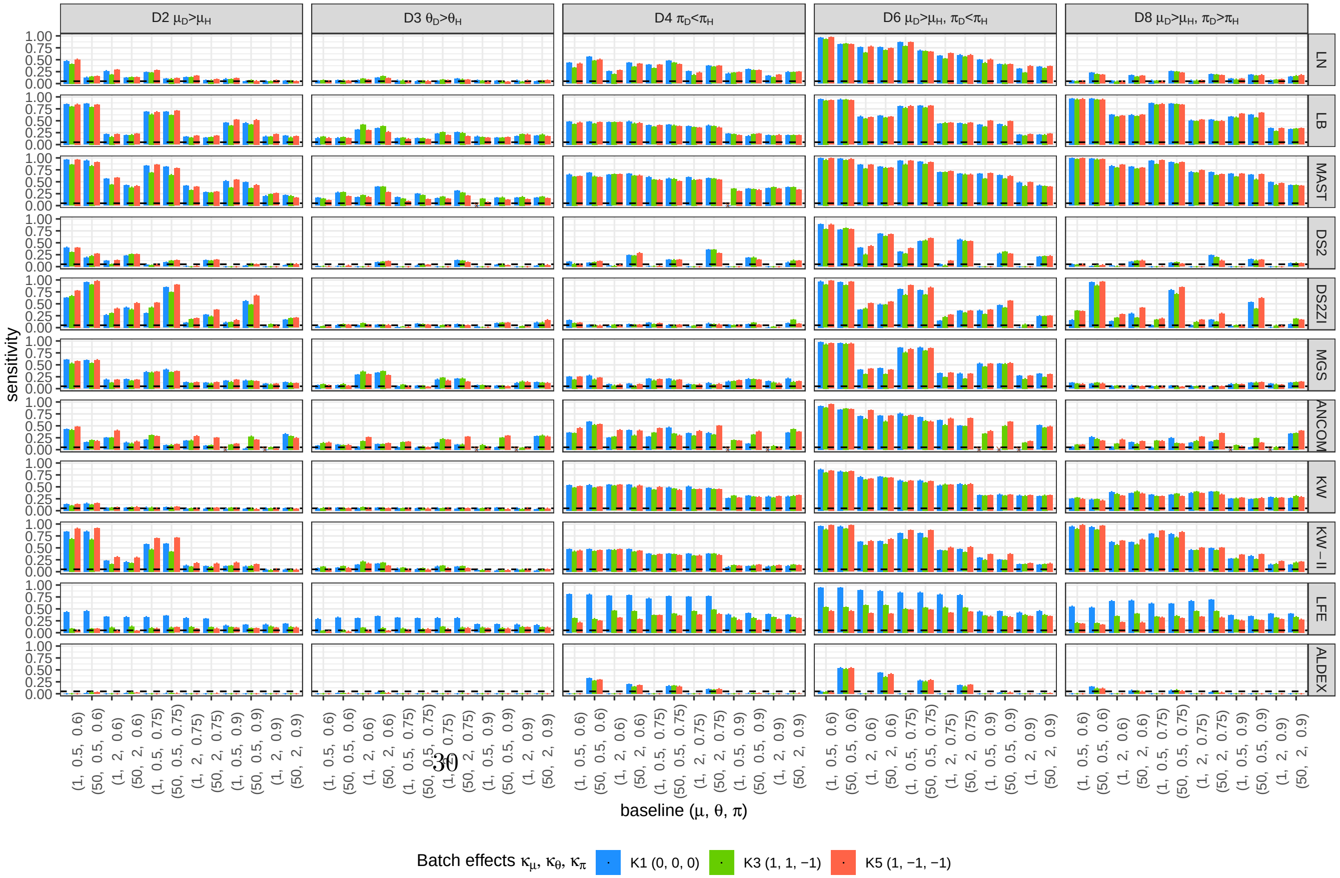




Web Figure 15: Full results under ZIG model for a sample size of 80. DS2 = DESeq2, DS2ZI = DESeq2-ZINBWaVE, ANCOM.sz = ANCOM-BC1, ANCOM = ANCOM-BC2, LFE = LEfSe, ALDEX = ALDEx2.

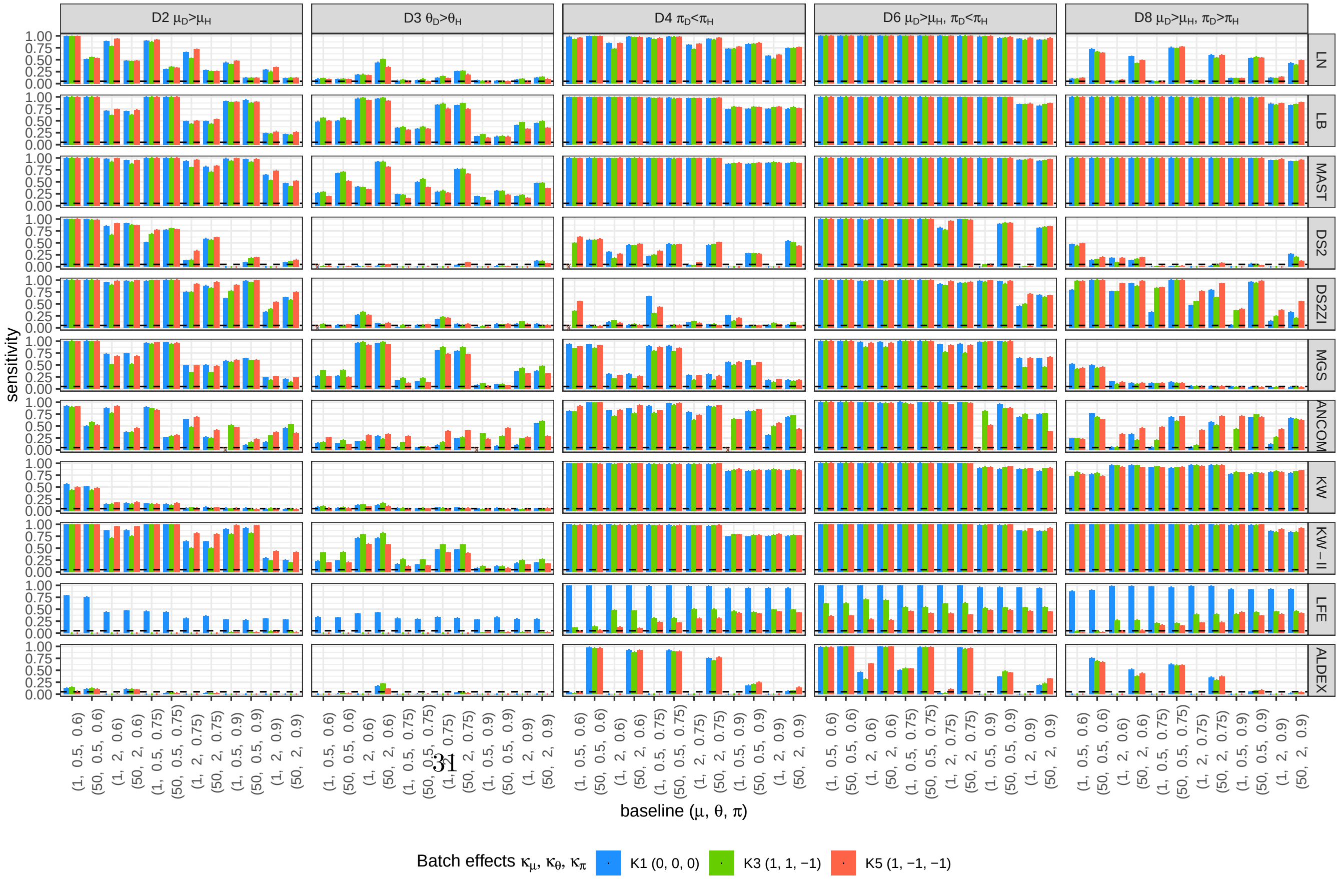


Web Figure 16: Full results under ZIG model for a sample size of 400. DS2 = DESeq2, DS2ZI = DESeq2-ZINBWaVE, ANCOM.sz = ANCOM-BC1, ANCOM = ANCOM-BC2, LFE = LEfSe, ALDEX = ALDEx2.

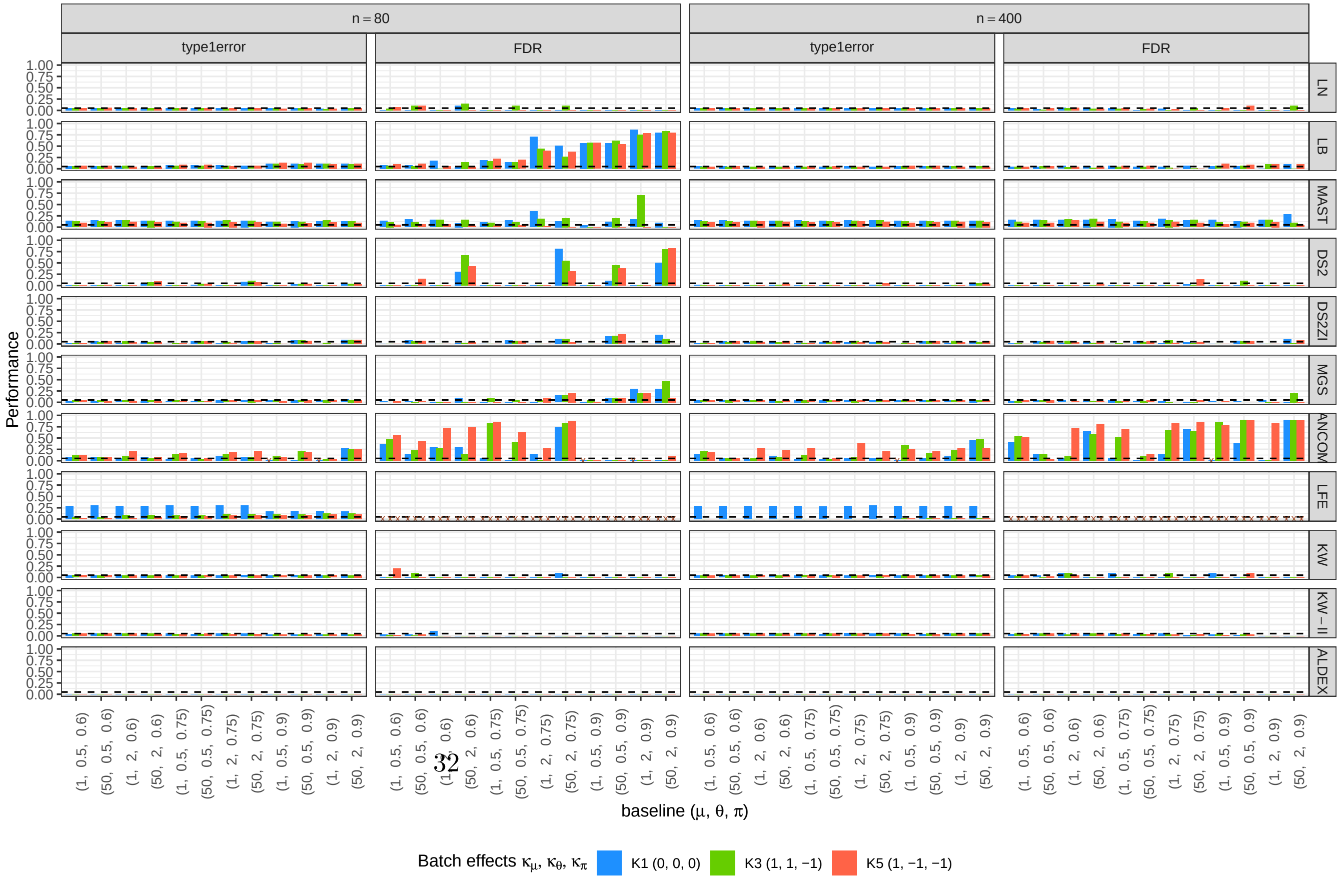




Web Figure 17: Type I error and FDR results under ZIG model under D2 scenario. DS2 = DESeq2, DS2ZI = DESeq2-ZINBWaVE, ANCOM.sz = ANCOM-BC1, ANCOM = ANCOM-BC2, LFE = LEfSe, ALDEX = ALDEx2.

### 6 Application to the ZOE 2.0 data

This section includes the analysis results from the ZOE 2.0 data in terms of species and gene-species combination. The same normalization and screening procedures are applied to these analyses except for the screening thresholds. As for the abundance-based screening, gene-species with TPM less than 0.2 were not tested and species were screened with the threshold of TPM being less than 2. Out of 535,299 gene-species combinations, after screening, 188,957 gene-species were tested. Out of 209 species, after screening, 97 species were tested.

#### 6.1 Application to the ZOE 2.0 data - gene-species

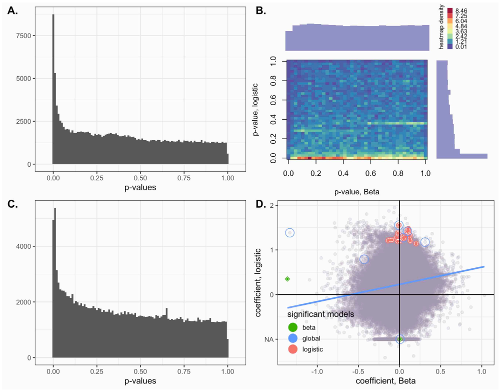


Web Figure 18: A. Histogram of the p-values of the log-normal models (gene-species combination). B. Histogram of the joint p-values of the logistic Beta models (logistic and Beta parts; gene-species combination). C. Histogram of the single p-values of the logistic Beta models (Wald statistics; gene-species combination). D. Scatter plot of the coefficients of the LB models (gene-species combination), with the circled dots representing the most significant gene-species—Wald statistic $p < 10^{-5}$.

The ten most significant gene-species combinations according to the LN model are C8PHV7 Campylobacter gracilis, C8PEV7 Campylobacter gracilis, C8PKG9 Campylobacter gracilis, C8PI10 Campylobacter gracilis, C8PH26 Campylobacter gracilis, C8PIH7 Campylobacter gracilis, C8PHR6 Campylobacter gracilis, C8PHV8 Campylobacter gracilis, C8PFD0 Campylobacter gracilis, and C8PG15 Campylobacter gracilis.

The seven gene-species combinations, of which global p-value is less than $10^{-5}$ according to the LB model, are E0DJ07 E0DJ07 Corynebacterium matruchotii, C8PHV7 Campylobacter gracilis, A3CQN5 Streptococcus cristatus, C8PEV7 Campylobacter gracilis, G1WEB2 Prevotella oulorum, C7NCB2 Leptotrichia shahii, and C8PHV8 Campylobacter gracilis.

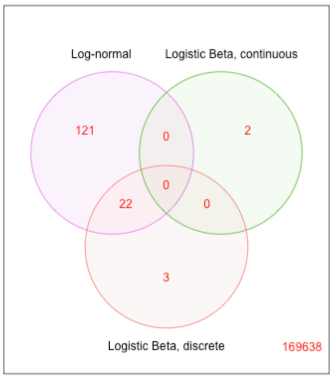


Web Figure 19: Venn diagram of the gene-species of which p-values are less than $10^{-5}$ for each model.

The species and proteins mapped to and the functions of those genes can be found in the Web Table 11

#### 6.2 Application to the ZOE 2.0 data - species

The eight species, of which global p-value is less than $10^{-2}$ according to the LB model, are Campylobacter gracilis, Streptococcus cristatus, Leptotrichia hofstadii, Lachnoanaerobaculum saburreum, Leptotrichia shahii, Streptococcus mutans, Campylobacter concisus, and Prevotella oulorum.

#### 6.3 Application to the ZOE 2.0 data - gene profiles

Web Table 11: Profiles of the top significant genes and gene-species based on the UniProt database UniProt Consortium (2019). * genes from the list of top significant genes, † genes from the list of top significant gene-species.

| gene ID<br>test reference | gene name<br>protein<br>function | organism |
|---|---|---|
| A3CQN5<br>† | rsmA<br>Ribosomal RNA small subunit methyltransferase A.<br>Specifically dimethylates two adjacent adenosines (A1518 and A1519) in the loop of a conserved hairpin near the 3'-end of 16S rRNA in the 30S particle. May play a critical role in biogenesis of 30S subunits. | Streptococcus sanguinis (strain SK36) |

| | | |
|---|---|---|
| C7NCB2 | Lebu_0017 | Leptotrichia buccalis (strain ATCC 14201 / DSM 1135 / JCM 12969 / NCTC 10249 / C-1013-b) |
| † | MATE efflux family protein. | |
| | GO - Molecular function. antiporter activity, xenobiotic transmembrane transporter activity. | |
| C8PEV7 | CAMGR0001_2596 | Campylobacter gracilis RM3268 |
| *† | Uncharacterized protein. | |
| | - | |
| C8PFD0 | CAMGR0001_2483 | Campylobacter gracilis RM3268 |
| † | Uncharacterized protein. | |
| | - | |
| C8PG15 | CAMGR0001_0808 | Campylobacter gracilis RM3268 |
| † | Transcriptional regulator, LysR family. | |
| | - | |
| C8PG93 | CAMGR0001_0886 | Campylobacter gracilis RM3268 |
| * | Tat pathway signal sequence domain protein. | |
| | - | |
| C8PH26 | CAMGR0001_2214 | Campylobacter gracilis RM3268 |
| *† | Uncharacterized protein. | |
| | - | |
| C8PHR6 | CAMGR0001_0512 | Campylobacter gracilis RM3268 |
| † | Methylenetetrahydrofolate reductase. | |
| | - | |
| C8PHV7 | CAMGR0001_0553 | Campylobacter gracilis RM3268 |
| *† | Uncharacterized protein. | |
| | - | |
| C8PHV8 | CAMGR0001_0554 | Campylobacter gracilis RM3268 |
| *† | Uncharacterized protein. | |
| | - | |
| C8PI10 | serS | Campylobacter gracilis RM3268 |
| *† | Serine–tRNA ligase. | |
| | Catalyzes the attachment of serine to tRNA(Ser). Is also able to aminoacylate tRNA(Sec) with serine, to form the misacylated tRNA L-seryl-tRNA(Sec), which will be further converted into selenocysteinyl-tRNA(Sec). | |
| C8PIH7 | accD | Campylobacter gracilis RM3268 |
| *† | Acetyl-coenzyme A carboxylase carboxyl transferase subunit beta. | |
| | Component of the acetyl coenzyme A carboxylase (ACC) complex. Biotin carboxylase (BC) catalyzes the carboxylation of biotin on its carrier protein (BCCP) and then the CO2 group is transferred by the transcarboxylase to acetyl-CoA to form malonyl-CoA. | |
| C8PKG9 | CAMGR0001_0190 | Campylobacter gracilis RM3268 |

| | | |
|---|---|---|
| *† | NFACT-R_1 domain-containing protein.<br>- | |
| C8PKZ2<br>* | asd<br>Aspartate-semialdehyde dehydrogenase.<br>Catalyzes the NADPH-dependent formation of L-aspartate-semialdehyde (L-ASA) by the reductive dephosphorylation of L-aspartyl-4-phosphate. | Campylobacter gracilis RM3268 |
| C8PJD1<br>* | ilvC<br>Ketol-acid reductoisomerase (NADP(+)).<br>Involved in the biosynthesis of branched-chain amino acids (BCAA). Catalyzes an alkyl-migration followed by a ketol-acid reduction of (S)-2-acetolactate (S2AL) to yield (R)-2,3-dihydroxy-isovalerate. In the isomerase reaction, S2AL is rearranged via a Mg-dependent methyl migration to produce 3-hydroxy-3-methyl-2-ketobutyrate (HMKB). In the reductase reaction, this 2-ketoacid undergoes a metal-dependent reduction by NADPH to yield (R)-2,3-dihydroxy-isovalerate. | Campylobacter gracilis RM3268 |
| C8PJY1<br>* | pyrH<br>Uridylate kinase.<br>Catalyzes the reversible phosphorylation of UMP to UDP. | Campylobacter gracilis RM3268 |
| E0DI62<br>* | HMPREF0299 5372<br>Sua5/YciO/YrdC/YwlC family protein<br>- | Corynebacterium matruchotii ATCC 14266 |
| E0DJ07<br>† | HMPREF0299 5672<br>Phosphoserine transaminase<br>- | Corynebacterium matruchotii ATCC 14266 |
| G1WEB2<br>† | HMPREF9431_02084<br>Uncharacterized protein.<br>- | Prevotella oulorum F0390 |

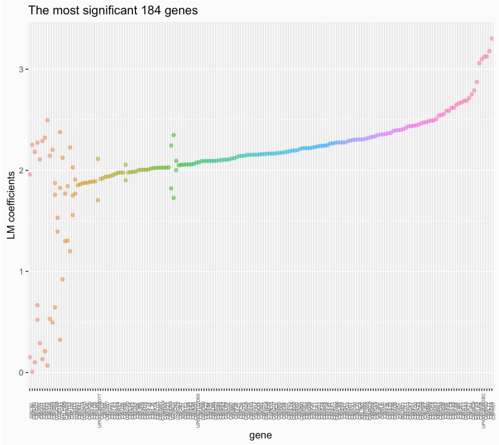


Web Figure 20: LN model coefficients of the gene-species for each of the most significant genes

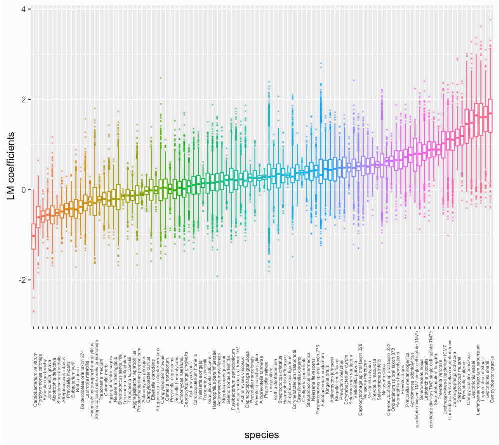


Web Figure 21: LN model coefficients of the gene-species for each species

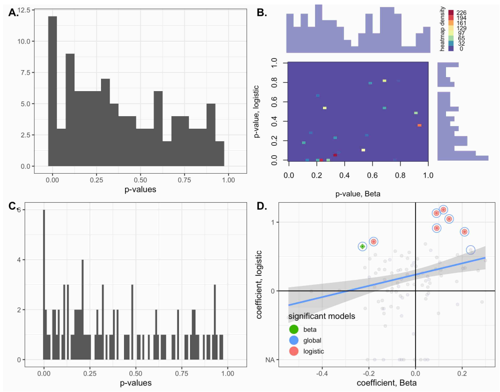


Web Figure 22: A. Histogram of the p-values of the log-normal models (for species). B. Histogram of the joint p-values of the logistic Beta models (logistic and Beta parts; for species). C. Histogram of the single p-values of the logistic Beta models (Wald statistics; for species). D. Scatter plot of the coefficients of the LB models (for species), with the circled dots representing the most significant species—Wald statistic $p < 0.01$.

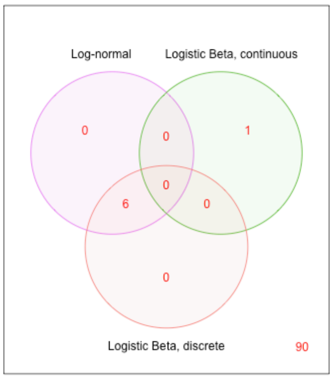


Web Figure 23: Venn diagram of the species of which p-values are less than 0.01 for each model.

### 7 Application to the IBD data

#### 7.1 Application to the IBD data - gene profiles

Web Table 12: Profiles of the top significant genes based on the UniProt database UniProt Consortium (2019). * genes from the list of top significant genes.

| gene ID<br>test reference | gene name<br>protein<br>function | organism |
|---|---|---|
| R5PRG3<br>* | BN489_01283<br>PlsC domain-containing protein<br>acyltransferase activity | Sutterella wadsworthensis CAG:135 |
| R5QAG2<br>* | BN489_02406<br>Polyamine ABC transporter<br>ATP binding | Sutterella wadsworthensis CAG:135 |
| R5QE55<br>* | BN489_00474<br>Leucine efflux protein<br>amino acid transport | Sutterella wadsworthensis CAG:135 |
| R5PLJ0<br>* | BN489_01810<br>HTH gntR-type domain-containing protein<br>DNA binding | Sutterella wadsworthensis CAG:135 |
| R5Q3H7<br>* | BN489_01382<br>Uncharacterized protein<br>- | Sutterella wadsworthensis |
| R5Q1H1<br>* | BN489_00687<br>Uncharacterized protein<br>- | Sutterella wadsworthensis CAG:135 |
| S3CE88<br>* | HMPREF1476_01419<br>Uncharacterized protein<br>transmembrane transport | Sutterella wadsworthensis HGA0223 |
| R5QEQ4<br>* | BN489_00642<br>Uncharacterized protein<br>- | Sutterella wadsworthensis CAG:135 |
| G2T243<br>* | RHOM_03530<br>Stage 0 sporulation protein A homolog<br>May play the central regulatory role in sporulation. It may be an element of the effector pathway responsible for the activation of sporulation genes in response to nutritional stress. Spo0A may act in concert with spo0H (a sigma factor) to control the expression of some genes that are critical to the sporulation process. | Roseburia hominis |
| S3BF40<br>* | HMPREF1476_00737<br>CN hydrolase domain-containing protein<br>nitrogen compound metabolic process | Sutterella wadsworthensis HGA0223 |

| | | |
|---|---|---|
| R5Q7C5<br>* | BN489_02016<br>Uroporphyrin-III C/tetrapyrrole methyltransferase<br>methyltransferase activity | Sutterella wadsworthensis CAG:135 |
| R5PTS5<br>* | BN489_00648<br>Uncharacterized protein<br>- | Sutterella wadsworthensis CAG:135 |
| D4IJ04<br>* | AL1_02120<br>Uncharacterized protein<br>- | Alistipes shahii WAL 8301 |
| R5PRP2<br>* | BN489_00146<br>Acetylglutamate kinase<br>ATP binding, kinase activity, cellular amino acid biosynthetic process | Sutterella wadsworthensis CAG:135 |
| R5PJG3<br>* | BN489_00982<br>Uncharacterized protein<br>- | Sutterella wadsworthensis CAG:135 |
| R5PM43<br>* | BN489_01889<br>Uncharacterized protein<br>- | Sutterella wadsworthensis CAG:135 |
| R5W0F1 (R5W0F1_9BACT)<br>* | (Obsolete)<br>(Obsolete)<br>- | Alistipes sp. CAG:53 |
| R5PWS5<br>* | BN489_02130<br>Uncharacterized protein<br>amino acid transport | Sutterella wadsworthensis |
| R5PNF6<br>* | BN489_02005<br>Uncharacterized protein<br>sulfurtransferase activity | Sutterella wadsworthensis |
| D4WIY6<br>* | rplC<br>50S ribosomal protein L3<br>rRNA binding, structural constituent of ribosome, translation | Bacteroides ovatus SD CMC 3f |
| Q0TKG5<br>* | aes<br>Acetyl esterase<br>Displays esterase activity towards short chain fatty esters (acyl chain length of up to 8 carbons). Able to hydrolyze triacetylglycerol (triacetin) and tributyrylglycerol (tributyrin), but not trioleylglycerol (triolein) or cholesterol oleate. Negatively regulates MalT activity by antagonizing maltotriose binding. Inhibits MelA galactosidase activity. | Escherichia coli O6:K15:H31 |
| D1PDG3<br>* | PREVCOP_05253<br>ISPg3, transposase<br>- | Prevotella copri DSM 18205 |
| Q17UW4<br>* | COII<br>Cytochrome c oxidase subunit 2 (Fragment) | Zygosaccharomyces rouxii (Candida mogii) |

| | |
|---|---|
| | Component of the cytochrome c oxidase, the last enzyme in the mitochondrial electron transport chain which drives oxidative phosphorylation. The respiratory chain contains 3 multisubunit complexes succinate dehydrogenase (complex II, CII), ubiquinol-cytochrome c oxidoreductase (cytochrome b-c1 complex, complex III, CIII) and cytochrome c oxidase (complex IV, CIV), that cooperate to transfer electrons derived from NADH and succinate to molecular oxygen, creating an electrochemical gradient over the inner membrane that drives transmembrane transport and the ATP synthase. Cytochrome c oxidase is the component of the respiratory chain that catalyzes the reduction of oxygen to water. Electrons originating from reduced cytochrome c in the intermembrane space (IMS) are transferred via the dinuclear copper A center (CU(A)) of subunit 2 and heme A of subunit 1 to the active site in subunit 1, a binuclear center (BNC) formed by heme A3 and copper B (CU(B)). The BNC reduces molecular oxygen to 2 water molecules using 4 electrons from cytochrome c in the IMS and 4 protons from the mitochondrial matrix. |
| E2ZM16<br>* | HMPREF9436_02727 Faecalibacterium cf. prausnitzii KLE1255<br>DUF3887 domain-containing protein<br>Enzyme and pathway databases: FCF748224-HMP:GTSS-1911-MONOMER |
| R7NP61 (R7NP61_9BACE)<br>* | (Obsolete) Bacteroides sp. CAG:98<br>(Obsolete) iron-regulated protein A<br>- |
| U2ZZD9 (U2ZZD9_VIBAL)<br>* | (Obsolete) Vibrio alginolyticus<br>(Obsolete) partial hypothetical protein<br>- |
| R6W6W2<br>* | BN607_03102 Bacteroides faecis CAG:32<br>Uncharacterized protein<br>- |
| I9USK4<br>* | HMPREF1074_02638 Bacteroides xylanisolvens CL03T12C04<br>Uncharacterized protein<br>- |
| B0NN15<br>* | BACSTE_00845 Bacteroides stercoris ATCC 43183<br>Uncharacterized protein<br>- |

#### 7.2 Application to the IBD data - alternative tests

Alternative tests to the logistic regression of the LB test were implemented using the likelihood-ratio and Fisher's exact tests. The likelihood-ratio test statistic is obtained by twice the difference in the log-likelihoods of the full model and the reduced model. The full model is the same as specified in the main body of paper (Section 6), and the reduced model simply lacks the disease feature from the full model. The p-value is obtained from the reference distribution of $\chi^2_1$. Fisher's exact test may not be as

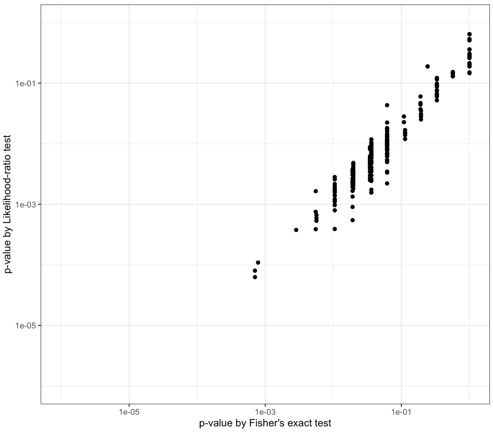


Web Figure 24: P-values of the likelihood-ratio and Fisher's exact tests for genes with one of the subgroup prevalence rate at the boundary in the IBD data

powerful as the likelihood-ratio test because it can only be applied in the case of a two-by-two table, thus not being able to account for any covariates. Figure 24 confirms that the p-values from the Exact test are overall larger than those of the likelihood-ratio test. None of the two tests identified significant genes at $10^{-5}$ significance level.
